# Nanoscale resolution imaging of the whole mouse embryos and larval zebrafish using expansion microscopy

**DOI:** 10.1101/2021.05.18.443629

**Authors:** Jueun Sim, Chan E Park, In Cho, Kyeongbae Min, Minho Eom, Seungjae Han, Hyungju Jeon, Hyun-Ju Cho, Eun-Seo Cho, Ajeet Kumar, Yosep Chong, Jeong Seuk Kang, Kiryl D. Piatkevich, Erica E. Jung, Du-Seock Kang, Seok-Kyu Kwon, Jinhyun Kim, Ki-Jun Yoon, Jeong-Soo Lee, Edward S. Boyden, Young-Gyu Yoon, Jae-Byum Chang

**Affiliations:** Department of Materials Science and Engineering, Korea Advanced Institute of Science and Technology, Daejeon, Korea; Department of Biomedical Engineering, Sungkyunkwan University, Suwon, Republic of Korea; School of Electrical Engineering, Korea Advanced Institute of Science and Technology, Daejeon, Republic of Korea; Brain Science Institute, Korea Institute of Science and Technology, Seoul, Republic of Korea; Disease Target Structure Research Center, Korea Research Institute of Bioscience and Biotechnology, Daejeon, Republic of Korea; Dementia DTC R&D Convergence Program, Korea Institute of Science and Technology, Seoul, Republic of Korea; Department of Biological Science, Korea Advanced Institute of Science and Technology, Daejeon, Republic of Korea; Department of Hospital Pathology, Uijeongbu St. Mary’s Hospital, College of Medicine, The Catholic University of Korea, Uijeongbu, Republic of Korea; John A. Paulson School of Engineering and Applied Sciences, Harvard University, Cambridge, MA, USA; School of Life Sciences, Westlake University, Hangzhou, Zhejiang, China; Westlake Laboratory of Life Sciences and Biomedicine, Westlake University, Hangzhou, Zhejiang, China; Institute of Basic Medical Sciences, Westlake Institute for Advanced Study, Hangzhou, Zhejiang, China; Department of Mechanical and Industrial Engineering, The University of Illinois at Chicago, Chicago, IL, USA; Graduate School of Medical Science and Engineering, Korea Advanced Institute of Science and Technology, Daejeon, Republic of Korea; Division of Bio-Medical Science and Technology, KIST School, Korea University of Science and Technology, Seoul, Republic of Korea; KRIBB school, University of Science and Technology, Daejeon, Republic of Korea; Howard Hughes Medical Institute, Cambridge, MA, USA.; McGovern Institute, Massachusetts Institute of Technology, Cambridge, MA, USA.; Departments of Brain and Cognitive Sciences, Media Arts and Sciences, and Biological Engineering, Massachusetts Institute of Technology, Cambridge, MA, USA.; KAIST Institute for Health Science and Technology, Daejeon, Republic of Korea

## Abstract

Nanoscale resolution imaging of whole vertebrates is required for a systematic understanding of human diseases, but this has yet to be realized. Expansion microscopy (ExM) is an attractive option for achieving this goal, but the expansion of whole vertebrates has not been demonstrated due to the difficulty of expanding hard body components. Here, we demonstrate whole-body ExM, which enables nanoscale resolution imaging of anatomical structures, proteins, and endogenous fluorescent proteins (FPs) of whole zebrafish larvae and mouse embryos by expanding them fourfold. We first show that post-digestion decalcification and digestion kinetics matching are critical steps in the expansion of whole vertebrates. Then, whole-body ExM is combined with the improved pan-protein labeling approach to demonstrate the three-dimensional super-resolution imaging of antibody- or FP-labeled structures and all major anatomical structures surrounding them. We also show that whole-body ExM enables visualization of the nanoscale details of neuronal structures across the entire body.

## INTRODUCTION

A systematic understanding of biological systems requires an unbiased investigation into entire vertebrate bodies, not just specific organs, at sub-cellular resolutions.^1–3^ For instance, in developmental biology, a high spatial resolution is essential for discerning changes in subcellular structures during different developmental stages, whereas full spatial coverage is needed for organ morphogenesis analysis from a systematic perspective.^4^ This type of high-resolution panoptic imaging is also crucial for researching nervous systems, as changes in cellular organelles or nanoscale neuronal structures must be observed across the entire body.^5, 6^ Various imaging techniques, including tissue clearing,^5–12^ computed tomography,^13^ and electron microscopy,^14^ have been developed to visualize the whole organism. Computed tomography and tissue clearing do not rely on physical sectioning, allowing for non-invasive, three-dimensional histological analysis.^2, 13^ These techniques have a resolution of 0.5–2 μm, which is sufficient for imaging micron-scale morphological and molecular structures. Using these techniques, neural projections from the brain to internal organs have been imaged throughout the vertebrate body.^5, 6, 9^ Electron microscopy, on the other hand, can achieve a resolution of up to 3 nm on ultra-thin specimens.^15^ Electron microscopy has been combined with automated sectioning and successfully imaged the nanoscale details of neuronal structures, even individual synaptic vesicles, over a hundred microns^15, 16^ However, what is missing is a technique that enables the imaging of centimeter-size entire vertebrate bodies with the sub-100 nm resolution required for resolving the nanoscale details of cellular organelles and dense neural structures.

Expansion microscopy (ExM) is an attractive candidate for satisfying such needs. ExM can image proteins or mRNA with a lateral resolution of 60 nm by physically expanding the specimens.^17–40^ As the ExM procedure also optically clears specimens, images of specimens with a thickness of hundreds of microns could be obtained without thin-sectioning, which is required for other super-resolution microscopy techniques.^41^ ExM has been successfully used on a variety of specimens, including cultured cells^17–30^ animal organ slices,^21–35^ entire mouse organs,^35^ *Drosophila* brains,^36, 37^ and zebrafish larvae brains.^38^ ExM has also been used on entire organisms, including parasite microorganisms^36, 39^ and *Caenorhabditis elegans*.^40^ However, subcellular-resolution imaging of whole vertebrate bodies with bones by expanding the bodies has yet to be demonstrated. A possible solution is to divide the vertebrate bodies into distinct organs, expand them, and stitch the images together; however, this approach has some limitations. First, structures spanning the entire body, such as axons inside axon bundles, cannot be stitched correctly due to their small size and high density, which require resolutions beyond the diffraction limit of the light. Second, during specimen preparation, structures at the interfaces between organs or inter-organ connections may be destroyed. Third, the three-dimensional morphologies, locations, and orientations of organs may shift. Most importantly, no technique for expanding hard body parts, such as bones and spines, has been demonstrated, which is required to study neural connections from the brain to the body.

In this paper, we demonstrate a new ExM technique, whole-body ExM, which enables the imaging of whole mouse embryos and zebrafish larvae with a 60 nm resolution by expanding them more than fourfold. The key to enabling the uniform expansion of whole mouse embryos and zebrafish larvae is bone decalcification. The decalcification process must be applied after the gelation and proteinase digestion of the specimens to expand the whole mouse embryos and zebrafish larvae more than fourfold. In addition, the organs’ digestion kinetics must be matched to prevent crack formation at the interfaces between relatively hard and soft organs during digestion. Pan-protein staining through fluorophore NHS (*N*-hydroxysuccinimidyl) ester staining, antibody staining, and genetically encoded fluorescent proteins (FPs) are compatible with the whole-body ExM process. Whole-body ExM is first applied to zebrafish larvae to visualize the nanoscale details of anatomical structures that cannot be resolved with diffraction-limited microscopy. This process is then applied to mouse embryos to visualize the nanoscale details of anatomical and neuronal structures from the head to the tail.

## RESULTS

### Uniform expansion of all vertebrates

The whole-body ExM process started with the formaldehyde fixation of specimens. If necessary, specimens were stained with antibodies to label specific proteins following chemical fixation. Then, the specimens were treated with 6-((acryloyl)amino)hexanoic acid (AcX) to make all proteins in the specimens gel anchorable and then embedded in a swellable hydrogel. After gelation, specimens were digested with proteinase K for 12 h and treated with NHS esters of both hydrophilic and hydrophobic fluorophores. The specimens were then treated with a mixture of collagenase I, II, and IV for 24 h to prevent crack formation due to the different expansion speeds between organs during the digestion process. Following the collagenase mixture treatment, the specimens were further digested with proteinase K. Then, specimens were treated with ethylenediaminetetraacetic acid (EDTA) for 24 h to decalcify the bones and cartilage. After the decalcification process, the specimens were expanded in deionized (DI) water more than fourfold (**Fig. 1a**).

**Fig. 1:**
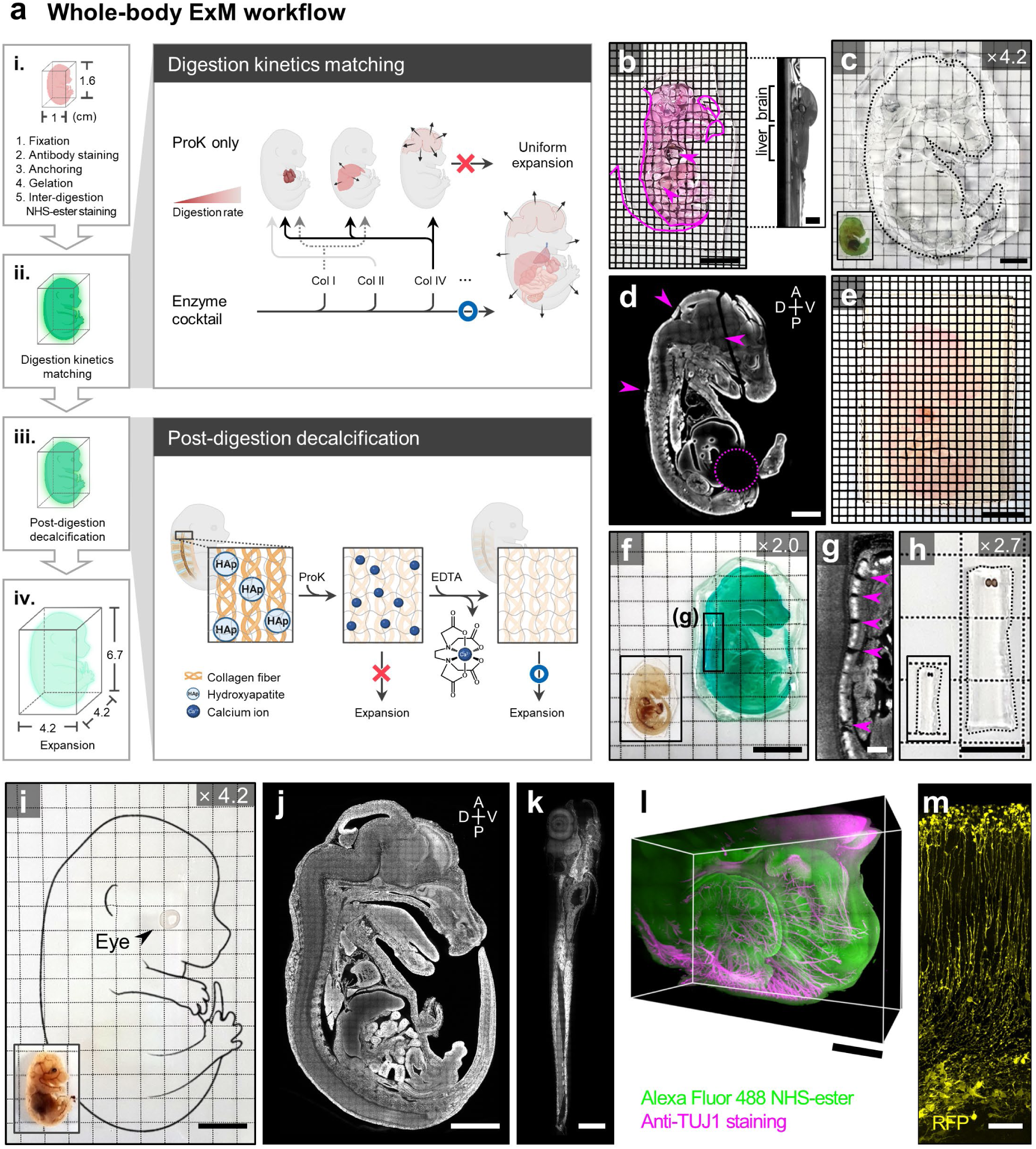
Whole-body ExM procedures and results. (**a**) Schematic illustrating the whole-body ExM workflow from sample preparation to whole vertebrate expansion: **i**, fixation, antibody staining (optional), anchoring, gelation, and inter-digestion NHS-ester staining; **ii**, digestion kinetics matching; **iii**, post-digestion decalcification; and **iv**, expansion. Each of the boxes in the middle describes the two key strategies for expanding the whole vertebrates. (**b**) Representative photographs of a mouse embryo slice digested with proteinase K for 72 h. Magenta arrows and lines indicate partial distortions and cracks in the specimen due to differences in digestion rates between the internal organs. The right box displays the side view of the sample. (**c**) Representative photograph of an expanded mouse embryo slice without digestion kinetics matching. The dotted lines indicate cracks in the specimen. The inset image displays the specimen before expansion. (**d**) Confocal microscopy image of the specimen in **c**. Magenta arrows indicate cracks in the specimen. A dotted circle indicates the area where the tissues have been severely damaged and lost. (**e**) Representative photographs of a mouse embryo slice digested with proteinase K for 24 h, collagenase mixture for 24 h, and proteinase K for 24 h. (**f**) Representative photograph of an expanded mouse embryo slice without post-digestion decalcification. The inset image displays the specimen before expansion. Compared to the original embryo size, the specimen showed only a twofold increase in length. (**g**) Confocal microscopy image of the boxed region in **f**. Magenta arrows indicate cracks between the bone joints due to incomplete bone expansion. (**h**) Representative photograph of an expanded zebrafish larva without post-digestion decalcification. The inset image displays the specimen before expansion. (**i**) Photograph of an E15.5 whole mouse embryo after expansion. The inset image displays the specimen before expansion. Compared to the original embryo size, the final expansion of the mouse embryo showed a more- than-fourfold increase in length. (**j**) Confocal microscopy image of an expanded E15.5 mouse embryo slice. (**k**) Confocal microscopy image of an expanded 6 dpf whole larval zebrafish. (**l**) Confocal microscopy image of the facial region of the whole-body ExM-processed mouse embryo. Anti-TUJ1 staining (magenta) was used to label facial nerve networks, while Alexa Fluor 488 NHS ester (green) was used to outline the morphology of the biological landmarks. (**m**) Confocal microscopy image of RFP-positive neural stem cells (yellow) in the whole-body ExM-processed mouse embryo. Samples: (**b**) E15.5 mouse embryo slices, 500 μm thick; (**c**–**d**) E15.5 mouse embryo slices, 2 mm thick; (**e**) E15.5 mouse embryo slices, 500 μm thick; (**f**–**g**) E15.5 mouse embryo slices, 1 mm thick; (**h**) 6 pdf zebrafish larva; (**i**) E15.5 whole-mouse embryo; (**j**) E15.5 mouse embryo slices, 500 μm thick; (**k**) 6 pdf zebrafish larva; (**l**) E13.5 half-mouse embryo; and (**m**) E14.5 mouse embryo slices, 500 μm thick. Scale bars: (**b**) 5 mm; inset in (**b**) 2 mm, (**c**– **d**) 1 cm, (**e**) 5 mm, (**f**) 1 cm, (**g**) 500 μm, (**h**) 5 mm, (**i**–**j**) 1 cm, (**k**) 250 μm, (**l**) 500 μm, and (**m**) 20 μm. All length scales are presented in pre-expansion dimensions.

The two key segments of the whole-body ExM process were digestion kinetics matching and post-digestion decalcification. The expansion of thick mouse organs other than the mouse brain is not trivial and requires optimization for each organ.^32^ We first attempted to develop an expansion protocol that could allow for the uniform expansion of all organs. We discovered that repeated digestion with proteinase K at a high temperature resulted in the uniform fourfold expansion of a variety of adult mouse organ slices with a thickness of 1 mm, including the heart, kidney, liver, lung, small intestine, spleen, and stomach (**Supplementary Fig**. **1**). However, unlike the expansion of single organs, the expansion of whole vertebrate bodies required the consideration of digestion kinetics. When a hydrogel was synthesized inside a mouse embryo and treated with proteinase K, the organs in the embryo showed different expansion factors during the digestion process, as shown in **Fig. 1b**. The heterogeneous expansion resulted in cracks at the interfaces between relatively more and less expanded organs. After the full expansion in DI water, such cracks were more evident, especially around heavily calcified body parts, such as the head and spine, as shown in **Fig. 1c**–**d** (see also **Supplementary Fig**. **2**). However, when a hydrogel-embedded embryo was treated with proteinase K for 24 h, a mixture of collagenase I, II, and IV for 24 h, and then again with proteinase K for longer than 24 h, the mouse embryo did not show such cracks, as shown in **Fig. 1e**. The collagenase mixture treatment might digest collagens in organs, especially organs with a high content of collagens, including bones, and render those organs more accessible to proteinase K. The use of a mixture of three collagenases was essential; when a single type of collagenase was used, cracks still developed.

More importantly, the uniform expansion of whole vertebrate bodies needed post-digestion decalcification. Since the first demonstration of expansion-based tissue imaging in 2015, the expansion of hard body parts, such as bones, has not yet been demonstrated. In the hydrogel-based clearing of bones, EDTA, which is a chelating agent of calcium ions, is applied prior to hydrogel synthesis.^42, 43^ Thus, we first tested the pre-gelation EDTA treatment. We treated fixed mouse embryos with 0.5 M EDTA for longer than three weeks and then proceeded to gelation, proteinase K treatment, collagenase mixture treatment, proteinase K treatment again, and then expansion in DI water. The resulting specimens were expanded only twofold, which was half of the expansion factor of the bare hydrogels (**Fig. 1f**). In addition, confocal microscopy imaging showed cracks around the calcified body parts, such as the spine (**Fig. 1g**). The pre-gelation EDTA enabled only a 2.7-fold expansion for 6 dpf (days post-fertilization) zebrafish larvae, which had fewer calcified body parts than mouse embryos, as shown in **Fig. 1h**. Extending the EDTA treatment for more than 3 months did not still enable the full expansion of larvae. We then applied EDTA treatment after digestion. The mouse embryos were embedded in a hydrogel, digested with proteinase K, collagenase mixture, and proteinase K again, and then treated with a decalcification buffer containing 0.3 M EDTA and 2 M NaCl. Sodium chloride was added to the decalcification buffer to prevent the expansion of the digested embryos and larvae before the complete removal of calcium. This protocol expanded E15.5 mouse embryos and 8 dpf zebrafish larvae 4.1-fold, which was similar to the expansion factor of bare hydrogels. The full expansion of the larvae with post-digestion decalcification indicated that the proteinase treatment rendered the calcium more accessible to EDTA and thus enabled their complete removal.

Through these two processes, whole mouse embryos up to E15.5 in stage and zebrafish larvae up to 8 dpf were successfully expanded, as shown in **Fig. 1i**. After expansion, the zebrafish larvae and mouse embryos were almost completely transparent. Zebrafish larvae older than 8 dpf were not tested; however, zebrafish juveniles 2 months in age were successfully expanded using the same process (data not shown). Mouse embryos typically show a high level of autofluorescence, and a step for removing autofluorescence is required in the clearing of mouse embryos;^44^ however, the autofluorescence levels in expanded mouse embryos were negligible due to the digestion of tissues and the volumetric dilution of molecules. The whole-body ExM procedure was compatible with pan-protein staining using fluorophore NHS esters (**Fig. 1j**–**l**), antibody staining (**Fig. 1l**), and genetically encoded fluorescent proteins (**Fig. 1m**).

### Optimization of fluorophore NHS ester staining

In addition to the whole-body ExM process, we also optimized pan-protein staining using fluorophore NHS esters to better visualize the protein structures inside mouse embryos and zebrafish larvae. Recently developed ExM techniques utilizing pan-molecular staining can visualize diverse cellular structures.^17, 26, 29^ However, the effects of the physicochemical properties of fluorophores on the range of structures that can be visualized have not been studied. We found that protein structures labeled with fluorophore NHS esters depended on the hydrophobicity of the fluorophores. Hydrophilic fluorophores, such as Alexa Fluor 488, labeled the nucleoplasm and showed uniform staining in the cytoplasm. Hydrophobic fluorophores, such as Cy3 and ATTO 647N, labeled lipid-rich structures, including membrane-bound cellular organelles, such as nuclear membranes, the endoplasmic reticulum (ER), Golgi apparatus, and mitochondria, as well as structures bearing multiple membranes, such as myelinated axons. The use of both hydrophilic and hydrophobic fluorophores enabled the identification of specific cellular organelles using pan-protein labeling (see **Supplementary Notes 1**–**2**, **Supplementary Figs. 3–4**, and **Supplementary Video 1** for a detailed discussion). Next, we optimized the fluorophore NHS ester labeling process to label more anatomically relevant structures in diverse mouse organs. When fluorophore NHS esters were applied after brief proteinase K digestion, the process we termed inter-digestion staining, the most diverse structures were visualized (see **Supplementary Note 3** and **Supplementary Fig. 4**). We chose ATTO 647N over Cy3 as a hydrophobic fluorophore because of its greater fluorescence signal retention after digestion. Using a transgenic mouse line, we confirmed that the mitochondria were stained with ATTO 647N (see **Supplementary Note 4** and **Suppplementary Figs. 5–6**). We applied the inter-digestion process of fluorophore NHS ester labeling to various mouse organs, such as the liver, esophagus, stomach, small intestine, kidney, lung, heart, and testis. In all these organs, the inter-digestion staining using fluorophore NHS esters resolved the nanoscale structures, demonstrating that this method could be utilized to visualize all important anatomical structures over the whole body (see **Supplementary Note 5**, **Supplementary Fig**. **7**, and **Supplementary Video 2** for digestive organs; see **Supplementary Note 6**, **Supplementary Figs**. **8**–**9**, and **Supplementary Videos 3**–**5** for other representative organs).

### Whole-body ExM imaging of whole zebrafish larvae

We first tested whole-body ExM’s ability to visualize the ultrastructures over the whole organisms of 6 dpf zebrafish larvae (**Fig. 2a**; see **Supplementary Video 6** for annotated images). We used an objective with an NA of 1.15 for imaging, and the effective lateral resolution was 62 nm. Consistent with the staining patterns observed in the mouse organs, Alexa Fluor 488 was useful for determining their overall morphologies, and ATTO 647N was useful for revealing detailed organelle structures (**Supplementary Fig**. **10**). In the sensory systems, the nanoscale details of major sensory organs, such as the individual microvillus of the olfactory sensory epithelium (**Fig. 2b**,**c**; see **Supplementary Fig**. **11a**–**d** for more detailed analysis and **Supplementary Video 7** for a z-stack image) and the individual cilium and cells of the trunk neuromast (**Fig. 2d**; see **Supplementary Fig**. **11e**–**h** for more detailed analysis and **Supplementary Video 8** for a z-stack image) were visualized in the Alexa Fluor 488 channel. In addition, the otoliths and sensory patches of the inner ear were also clearly shown in the same channel. Individual otolithic membrane layers composed of the otolith and three cell layers of the otic sensory epithelium were difficult to identify in the Alexa Fluor 488 channel but were clearly resolved in the ATTO 647N channel (**Fig. 2e**,**f**). For cilia present in the hair cell bundles of the inner ear (dotted rectangles in **Fig. 2f**), the average center-to-center distance between neighboring cilia is smaller than 150 nm,^45^ and the distinction of individual cilia requires a resolution beyond that of optical microscopy. As shown in the inset of **Fig. 2f**, the whole-body ExM resolution was sufficiently high, allowing the visualization of the individual cilium of the hair cell bundles and the distinguishing between kinocilium (relatively long cilium; white arrows in **Fig. 2f**) and stereocilia (relatively short cilia; white arrowheads in **Fig. 2f**) based on their length (see also **Supplementary Video 9**). In the digestive system, most features of organ morphogenesis that are commonly observed in this developmental stage were clearly observed, along with the detailed shape of major internal organs, including the intestinal bulb, liver, pancreas, and esophagus (**Fig. 2g**). Interestingly, structures presumably of bacteria were also labeled and observed inside the intestinal bulb (inset of **Fig. 2g**; see **Supplementary Fig**. **12a**–**e** for detailed analysis and **Supplementary Video 10** for a z-stack image) due to the non-specific labeling nature of pan-protein staining. Whole-body ExM clearly visualized the microvilli of the intestinal epithelium (**Fig. 2h**; see **Supplementary Fig**. **12f**–**g**, and **Supplementary Video 11** for a z-stack image) and the details of the mid-intestine, including goblet cells, brush border, and secretory vesicles (**Supplementary Video 12**). In the urinary system, the pronephros (the earliest stage of kidney development; **Fig. 2i**), pronephric duct, and motile cilia in the pronephric duct along the whole body were shown (**Supplementary Fig**. **13**).

**Fig. 2:**
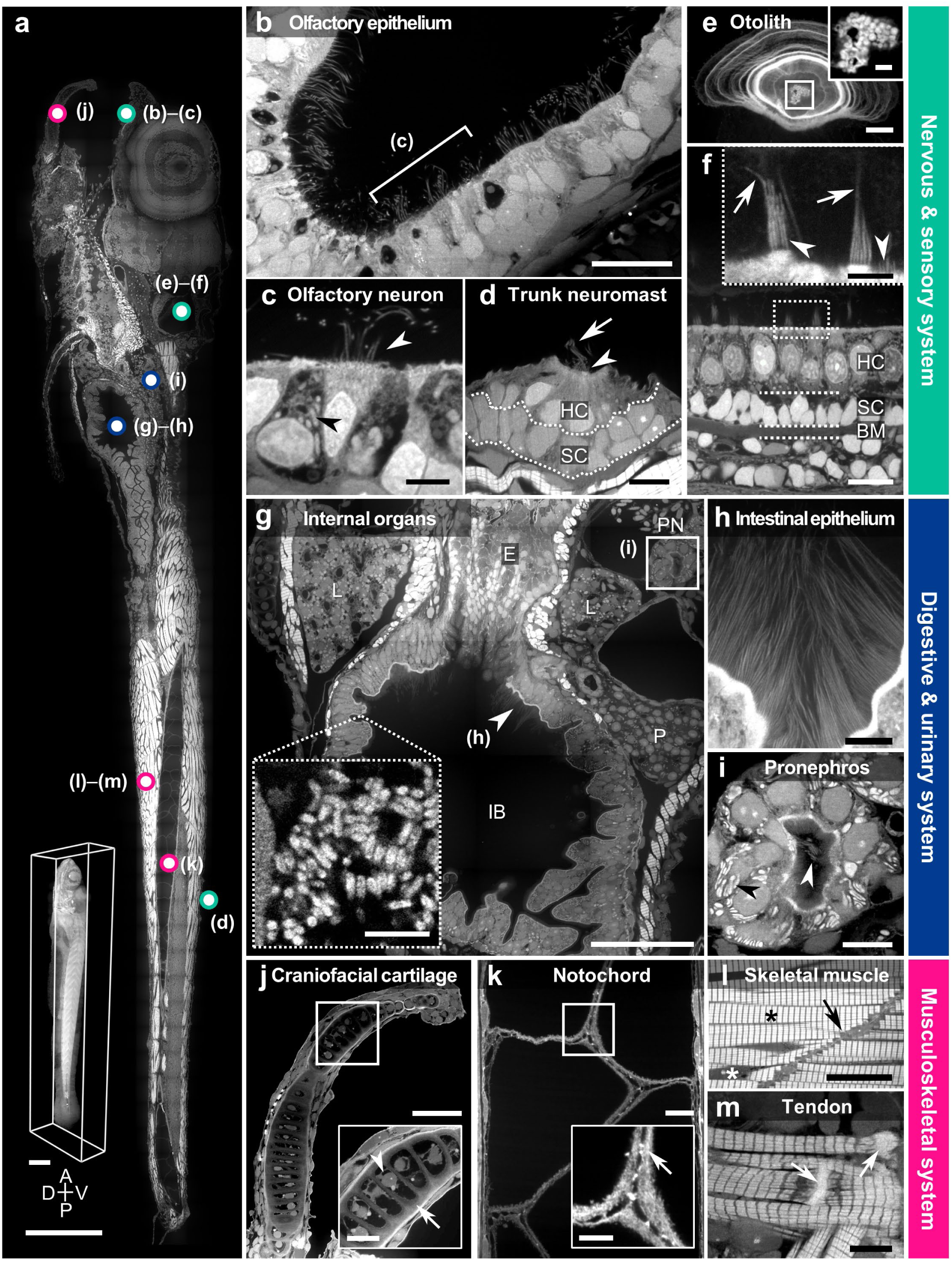
Whole-body ExM imaging of a variety of key landmarks in zebrafish larvae. (**a**) Confocal microscopy image of an expanded 6 dpf larval zebrafish. The inset displays a volumetric rendering of an expanded 6 dpf larval zebrafish. (**b**–**c**) Olfactory sensory epithelium. The bracket in **b** indicates several olfactory rod cells. (**c**) Olfactory neurons. Black arrowhead, mitochondria; white arrowhead, cilia. (**d**) Trunk neuromast. HC, hair cell; SC, supporting cell. Arrow, kinocilium; arrowhead, stereocilia. (**e**) Sagittal view of the otolith in which the layered biomatrix is intensely stained with ATTO 647N NHS ester. The boxed region shows the inner globular structures and is shown in the magnified view in the inset box. (**f**) Frontal view of the otic sensory epithelium (utricular macula). Hair cells (HC), supporting cells (SC), and the basilar membrane (BM) are marked. The inset shows the magnified view of the boxed region (taken from a different sample) and reveals the brush-like sensory hair cell bundles. Arrows, kinocilia; arrowheads, stereocilia. (**g**) Internal organs that belong to the digestive or urinary systems. E, esophagus; L, liver; IB, intestinal bulb; P, pancreas; and PN, pronephros. The dotted box highlights bacterial colonies attached to the intestinal bulb epithelium (observed in a different sample). (**h**) Unusually long (around 20 μm) microvilli found in the anterior intestinal bulb region adjacent to the esophageal intestinal junction. (**i**) Magnified view of the boxed region in **g** showing the frontal view of the pronephros. White arrowhead, motile cilia; black arrowhead, mitochondria. (**j**) Craniofacial cartilage. The inset highlights a chondrocyte (arrowhead) encapsulated by a cartilage matrix (arrow). (**k**) Notochord organized by vacuoles, sheath cells, and tightly apposed notochordal vacuolated cells. The inset highlights the cell membrane (arrow) between neighboring notochordal cells. (**l**) Magnified view of the skeletal muscles. White asterisk, nucleus; black asterisk, muscle fibers; black arrow, connective tissue. (**m**) Tendons (arrows) between the cartilaginous bone and the muscle fibers. Except for **j**, which was observed with an 8 dpf larva, the remaining images were observed with 6 dpf larvae. Labels: (**a**), (**b**), (**c**), (**d**), (**g**) and inset in (**g**), (**h**), (**i**), Alexa Fluor 488 NHS ester; (**e**), inset in (**e**), (**f**), inset in (**f**), (**j**), inset in (**j**), (**k**), and inset in (**k**), (**l**), (**m**), ATTO 647 NHS ester. Scale bars: (**a**) 250 μm, (**b**) 5 μm, (**c**) 2 μm, (**d**) 5 μm, (**e**) 5 μm, inset in (**e**) 1 μm, (**f**) 5 μm, inset in (**f**) 2 μm, (**g**) 100 μm, inset in (**g**) 5 μm, (**h**– **i**) 10 μm, (**j**) 50 μm, inset in (**j**) 10 μm, (**k**) 10 μm, inset in (**k**) 5 μm, (**l**) 20 μm, and (**m**) 10 μm. All length scales are presented in pre-expansion dimensions.

Whole-body ExM also successfully visualized complex musculoskeletal tissues (**Supplementary Video 13**), such as calcified cartilage (**Fig. 2j**; see **Supplementary Fig**. **14** for the whole pharyngeal arches), body trunk—including the notochord (**Fig. 2k** for the notochord; see **Supplementary Fig**. **15** for the body trunk of the more distal region and **Supplementary Video 14** for a z-stack image of notochord) and spinal cord (**Supplementary Fig**. **16**)—fins (**Supplementary Video 15** for a z-stack image), muscle fibers and muscle mitochondria (**Fig. 2l**; see **Supplementary Fig**. **17a**–**d** for a detailed analysis), and even tendons (**Fig. 2m**). To confirm that the larvae expanded homogeneously, we measured the length of the sarcomere of skeletal muscle fibers in the anterior trunk and compared it to the sarcomere length measured using electron microscopy images of larvae reported in previous studies. The average sarcomere length measured from the four expanded larvae was 1.95 ± 0.12 μm. We also measured sarcomere length in other body parts, such as the craniofacial regions, posterior trunks, and tails, but no differences were observed. The average sarcomere length measured from two independent electron microscopy imaging studies of larvae was 1.87 ± 0.07 μm.^46, 47^ Considering that these two values were measured from separate specimens and that muscles are more resistant to expansion than other tissues,^48^ the good agreement between these two values (4.3% difference) indicates that the expansion was homogeneous, and the whole-body ExM process preserved not only the morphology of the structures but also their physical dimensions (**Supplementary Fig**. **17e**–**g**).

### Use of whole-body ExM to study anatomical changes in different developmental stages

Whole-body ExM was useful in studying changes in the morphologies of anatomical landmarks during different developmental stages. The eye was chosen as the model organ. The Alexa Fluor 488 channel of expanded zebrafish larvae clearly showed the landmark structures of the eye, including the cornea, lens, retinal layers, and optic nerves (**Fig. 3a**–**c**). In the retinal layers, photoreceptors were uniformly stained with Alexa Fluor 488; however, the lipid-rich outer segments, which consist of a stack of disc membranes, were strongly stained with ATTO 647N (see **Supplementary Video 16** for a z-stack image).^49^ Among these structures, we focused on two cases: fiber cell maturation in the lens and photoreceptor cell maturation in the retina. In the anterior segment of the lens in the 8 dpf larva, highly organized layers of the lens fiber cells, lens epithelium, and cornea were distinguished (**Fig. 3d**). During lens maturation, fiber cells can be classified into two cell types, depending on their degree of maturation: primary lens fiber cells (PFCs) and secondary lens fiber cells (SFCs). PFCs are found close to the core of the lens, and SFCs are found in the more peripheral regions of the lens. PFCs show well-ordered hexagonal cellular arrangements, whereas SFCs show less-ordered arrangements and unique protrusions formed *via* the ball-and-socket process.^50^ We were able to classify these two different types of fiber cells based on their detailed anatomical features (**Fig. 3e**,**f**), such as the hexagonal profile of PFCs and the ball-and-socket junction of SFCs. Photoreceptor maturation in the retinal layer is another interesting example. At 6 dpf, two rows of inner segments were visible (**Fig. 3g**). At 8 dpf, an increase in the length of the outer segments was observed, but the extent of this increase differed depending on the photoreceptor subtype (**Fig. 3h**). In addition, two distinct rows of inner segments became more evident (**Fig. 3h**; see **Supplementary Video 16** for a z-stack image of the 8 dpf larval retina). Based on these morphological features, we could classify the photoreceptor cells into two different subtypes: double cone photoreceptor cells, which are characterized by shorter outer segments (cell i in **Fig. 3h**), and single cone photoreceptor cells or rod photoreceptor cells, which are characterized by a long outer segment (cell ii in **Fig. 3h**). The observations were in agreement with those of previous studies on the zebrafish retina using transmission electron microscopy (TEM).^45, 49–51^

**Fig. 3:**
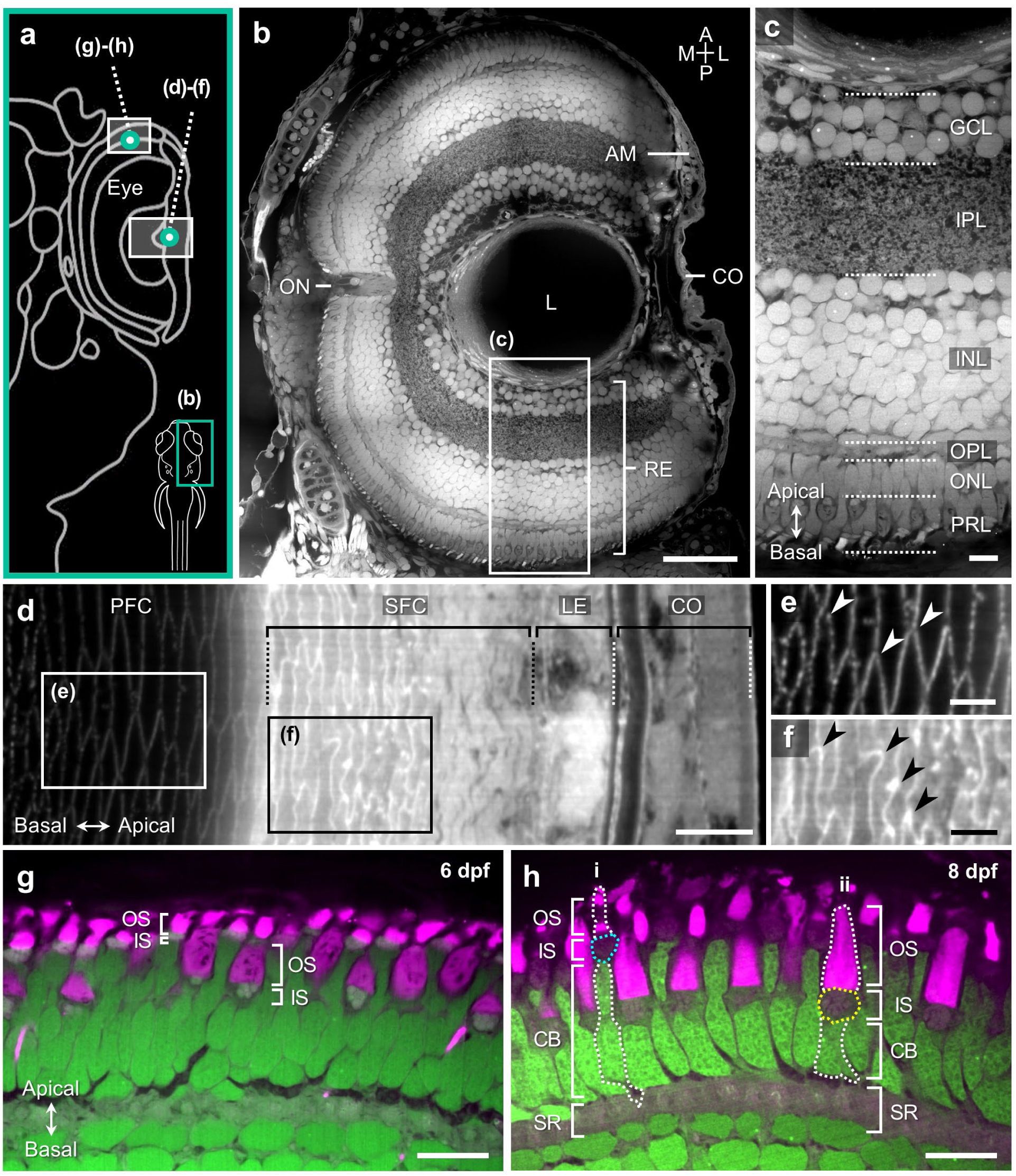
Whole-body ExM imaging of anatomical changes in zebrafish larvae in different developmental stages. (**a**) Schematic diagram of a larval zebrafish depicting locations in **b**–**h**. (**b**) Montage image of the larval eye showing major features of the vertebrate eye, including the angle mesenchyme (AM), optic nerve (ON), lens (L), cornea (CO), and retina (RE) layers. (**c**) Magnified view of the boxed region in **b**, showing the retinal sections organized into highly structured layers, including the ganglion cell layer (GCL), inner plexiform layer (IPL), inner nuclear layer (INL), outer plexiform layer (OPL), outer nuclear layer (ONL), and photoreceptor layer (PRL). (**d**) Confocal microscopy image of the anterior lens of 8 dpf zebrafish larva. PFC, primary lens fiber cell; SFC, secondary lens fiber cell; LE, lens epithelium; and CO, cornea. (**e**) Magnified view of the boxed region in **d** highlighting the detailed anatomical features of the PFCs. Arrowheads, Y-shaped suture pattern between the ends of the cells. (**f**) Magnified view of the boxed region in **d** showing the detailed structure of the SFCs. Arrowheads, ball-and-socket inter-digitations between older fiber cells. (**g**–**h**) Confocal microscopy images of the retinal layers of larval zebrafish at (**g**) 6 dpf and (**h**) 8 dpf depicting photoreceptor cell maturation. Green, Alexa Fluor 488 NHS ester; magenta, ATTO 647N NHS ester. OS, outer segment; IS, inner segment; CB, cell body; and SR, synaptic region. Labels: (**b**–**c**) Alexa Fluor 488 NHS ester, (**d**–**f**) ATTO 647N NHS ester, and (**g**–**h**) Alexa Fluor 488 NHS ester and ATTO 647N NHS ester. Scale bars: (**b**) 50 μm, (**c**) 10 μm, (**d**) 5 μm, (**e**–**f**) 2 μm, and (**g**–**h**) 10 μm. All length scales are presented in pre-expansion dimensions.

### Imaging of the fluorescent proteins of transgenic zebrafish by whole-body ExM

Finally, we tested the compatibility of whole-body ExM with molecular labeling techniques. We first labeled zebrafish larvae with an antibody against HuC/D and proceeded to the whole-body ExM process. After expansion, the antibody staining channel showed the characteristic expression patterns of HuC/D proteins, which were highly expressed in the cytoplasm of the cells in the inner nuclear layer (INL)^52^ (**Supplementary Fig**. **18a**,**b**). The high resolution offered by whole-body ExM enabled the visualization of the localization of this protein inside the cytoplasm; this protein was found outside linear structures strongly labeled with ATTO 647N, presumably the Golgi apparatus or ER (**Supplementary Fig**. **18c**–**e**, see yellow arrowheads). In addition, we also found that genetically encoded GFP signals could be retained during the whole-body ExM process, as shown in **Supplementary Fig**. **18f**,**g**. Whole-body ExM was also compatible with mRNA imaging. As shown in **Supplementary Fig**. **18h**,**i**, the whole-body ExM protocol combined with expansion fluorescent *in situ* hybridization (ExFISH)^20, 53^ visualized the mRNA puncta at an elevated resolution, enabling the nanoscale transcriptional analysis of the whole larvae.

### Expansion of whole mouse embryos

Next, we applied the whole-body ExM protocol to 0.5-mm-thick E11.5–15.5 mouse embryo slices. A 4.2-fold- expanded mouse embryo slice was then imaged with a 10× objective (NA 0.45) at an effective resolution of 158 nm (**Fig. 4a**; see also **Supplementary Video 17**). Again, in the mouse embryo, the fluorescence signals of Alexa Fluor 488 enabled the visualization of all major organs and anatomical landmarks from the expanded slice. In the brain, Alexa Fluor 488 showed strong fluorescence signals in the cortex, midbrain, pons, and choroid plexus. In the face, the otic cavity, trigeminal ganglion, optic cavity, pharynx, lip, and tongue were clearly identified due to their strong Alexa Fluor 488 signal intensities. In the abdomen, the substructures of the heart, lung, stomach, and kidney were identifiable. In this imaging plane, dorsal root ganglions (DRGs) were clearly shown on the dorsal side of the embryo. Relatively tough body parts, such as Meckel’s cartilage, vertebra, and ribs, did not show any signs of tissue damage or cracks. The whole-body ExM protocol enabled more than a fourfold expansion of unsectioned whole mouse embryos of stages up to E15.5 (**Fig. 4b**–**d**).

**Fig. 4:**
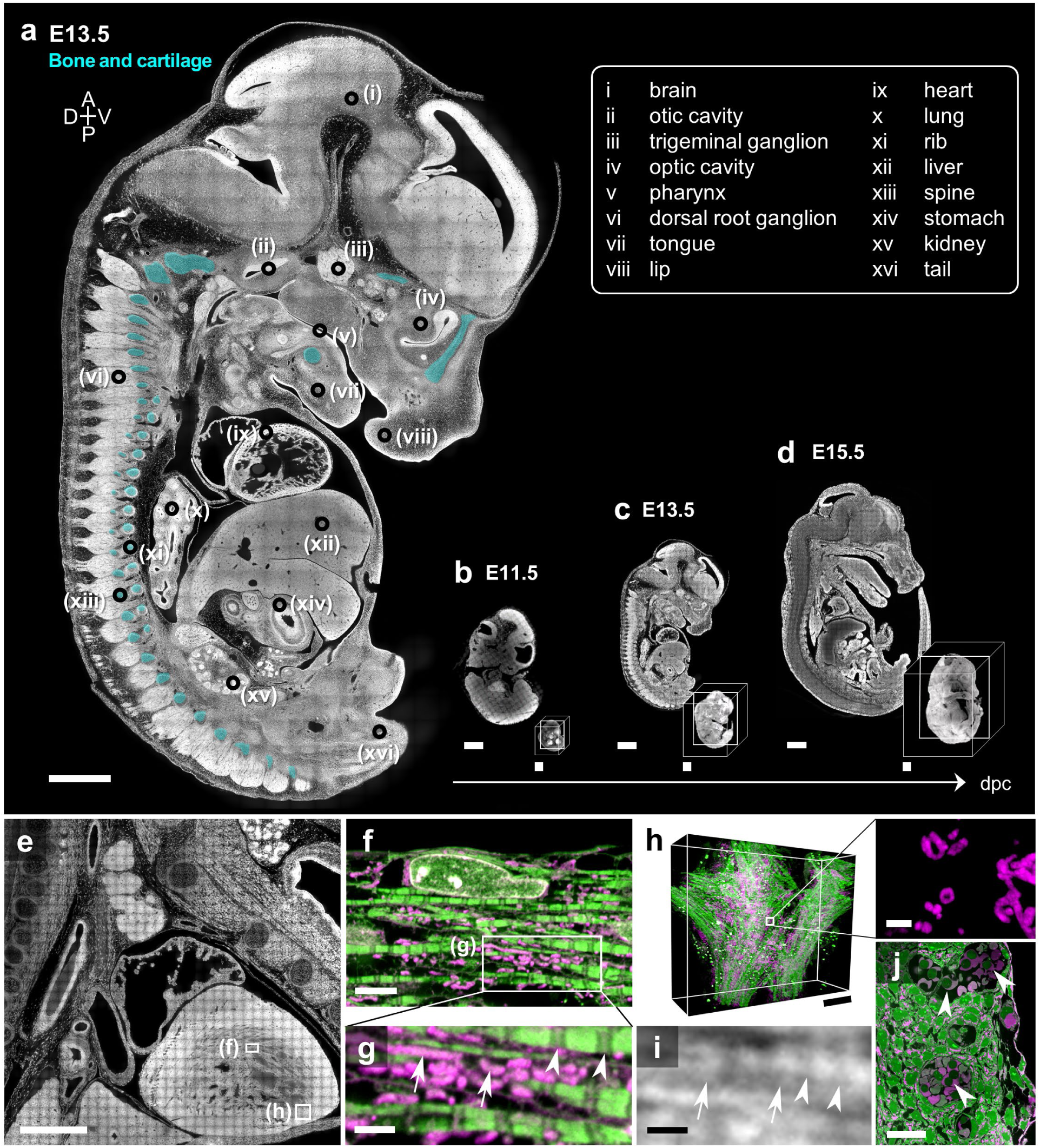
Whole-body ExM imaging of a mouse embryo in different developmental stages and length scales. **(a)** Confocal microscopy image of an expanded E13.5 mouse embryo slice. Colored regions (cyan) indicate bones and cartilage. (**b**–**d**) Confocal microscopy image of expanded mouse embryo slices at different embryonic stages: **b**, E11.5; **c**, E13.5; and **d**, E15.5. The insets show 3D-rendered images acquired from unsectioned whole mouse embryos after expansion. (**e**) Confocal microscopy image of the upper body of an E15.5 mouse embryo slice after expansion. (**f**) Magnified view of the boxed region in **e** showing the morphological details of the myofibrils. (**g**) Magnified view of the boxed region in **f**. Arrows, mitochondria; arrowheads, Z-disc. (**h**) Volumetric view of the boxed region in **e** showing cardiac tissue. The right box displays the magnified view of the mitochondria. (**i**) Confocal microscopy image of myofibrils taken from an unexpanded, BABB-cleared mouse embryo stained with Alexa Fluor 488 NHS ester. Arrows, mitochondria; arrowheads, Z-disc. (**j**) Confocal microscopy image of the coronary arteries of the E13.5 mouse embryo heart. Arrows, RBCs in the arteries. Labels: Gray and green, Alexa Fluor 488 NHS ester; Magenta, ATTO 647N NHS ester. Scale bars: (**a**) 1 mm, (**b**–**d**) 1 mm, (**e**) 500 μm, (**f**) 5 μm, (**g**) 2 μm, (**h**) 20 μm; right box in (**h**) 2 μm, (**i**) 2 μm, and (**j**) 20 μm. All length scales are presented in pre-expansion dimensions.

To confirm that the expansion resulted in an improved resolution, we imaged an unexpanded but cleared mouse embryo and an expanded mouse embryo with the same high NA objective. A 40× objective with a numerical aperture of 1.15 was used, and the theoretical resolution was 259 nm for unexpanded specimens and 62 nm for expanded specimens. Among the various structures, we compared images of the heart myofibrils. Imaging of expanded specimens with the high NA objective visualized the detailed cellular and subcellular structures, as shown in **Fig. 4e**. In the heart, single myofibrils and their characteristic structures, including the Z-disc, H-zone, M-line, and A-band, were resolved, as shown in **Fig. 4f**. More importantly, mitochondria were strongly labeled with ATTO 647N; their locations and morphologies were identifiable (**Fig. 4g**). Due to the increased lateral and axial resolutions, the three-dimensional visualization of myofibrils and mitochondria was possible, as shown in **Fig. 4h** (see **Supplementary Video 17** for the 3D rendered video). However, in the unexpanded but cleared mouse embryos, which were stained with both Alexa Fluor 488 and ATTO 647N, such nanoscale structures were not visible due to the limited lateral and axial resolutions (**Fig. 4i**). The high resolution offered by whole-body ExM would enable more comprehensive studies of heart diseases. Various heart diseases show abnormalities in the structures or alignments of myofibrils.^54–56^ The distinct staining patterns of hydrophilic and hydrophobic fluorophores were useful in detecting coronary arteries and mitochondria (**Fig. 4j**). Red blood cells (RBCs) in the arteries showed strong ATTO 647N signals, possibly due to their hydrophobic pockets;^57^ the coronary arteries inside the heart walls could be identified by the presence of RBCs (see **Supplementary Fig**. **19** for additional images).

The advantage of whole-body ExM is its ability to provide volumetric multiscale imaging of whole embryos and their internal organs. The use of a low-magnification objective, such as a 4× objective (NA 0.2), enables the three-dimensional imaging of organ morphologies with an effective resolution of 343 nm (**Supplementary Fig**. **20a**). Using a 10× objective (NA 0.45), the lateral resolution of 152 nm could be achieved, and the substructures of each organ, such as the individual cells in the heart valves or myocardium, could be resolved (**Supplementary Fig**. **20b**–**d**). Using a high-magnification objective with an NA above 1.0, a lateral resolution of 60 nm could be achieved, and the three-dimensional morphologies and arrangements of subcellular structures, including cellular organelles, were resolved (**Supplementary Fig**. **20e**–**h**). Such pan-length-scale imaging of the whole mouse embryo would enable the study of the changes in organ morphologies and underlying molecular changes in a single model animal. For example, for a large number of heart diseases, changes in the shape, size, and numbers of mitochondria in response to genetic and pharmacological perturbations of fission and fusion proteins have been observed.^58, 59^ TEM has been used as a gold standard method for studying such structural changes in myofibrils and mitochondria.^59^ Various TEM investigations have reported that depending on the type and stage of the heart disease, morphological measures varied accordingly.^59^ Whole-body ExM enables the super-resolution three-dimensional imaging of the whole heart of mouse embryos, providing more comprehensive insight into the diseases.

### The use of whole-body ExM to study the nanoscale details of the spine

We then illustrate the utility of whole-body ExM in imaging the ultrastructure of the spine, which consists of hard body parts, such as the bones, and soft structures, such as the spinal cord. Volumetric imaging of the spinal cord has become a regularly used method in spinal cord research, such as in the field of multiple sclerosis.^60^ For super-resolution imaging, spinal cords were removed from the spinal column and then expanded for such imaging due to the difficulties in expanding bones.^35, 61^ As whole-body ExM enabled the expansion of whole mouse embryos, it visualized the nanoscale details of the spinal cords inside the spines. Using a 4× objective to image expanded embryos, the spinal cord, spinal nerves, supraspinous ligament, and the characteristic structures of the vertebrae, including vertebral bodies, spinous processes, transverse processes, and intervertebral discs, were visualized, as shown in **Fig. 5a**–**c**. Different Alexa Fluor 488 fluorescence intensities were shown for white matter, which showed lower signal intensities, and the gray matter, which showed higher signal intensities (**Fig. 5a**). The high Alexa Fluor 488 signals of gray matter were attributed to the higher densities of nuclei and somas, as shown in **Fig. 5d**. The mosaic image of the spinal cord acquired at one z-plane image showed multiple layered structures having distinctive structures, including the meninges, spinal cords consisting of four layers, and vertebrae (**Fig. 5d**).

**Fig. 5:**
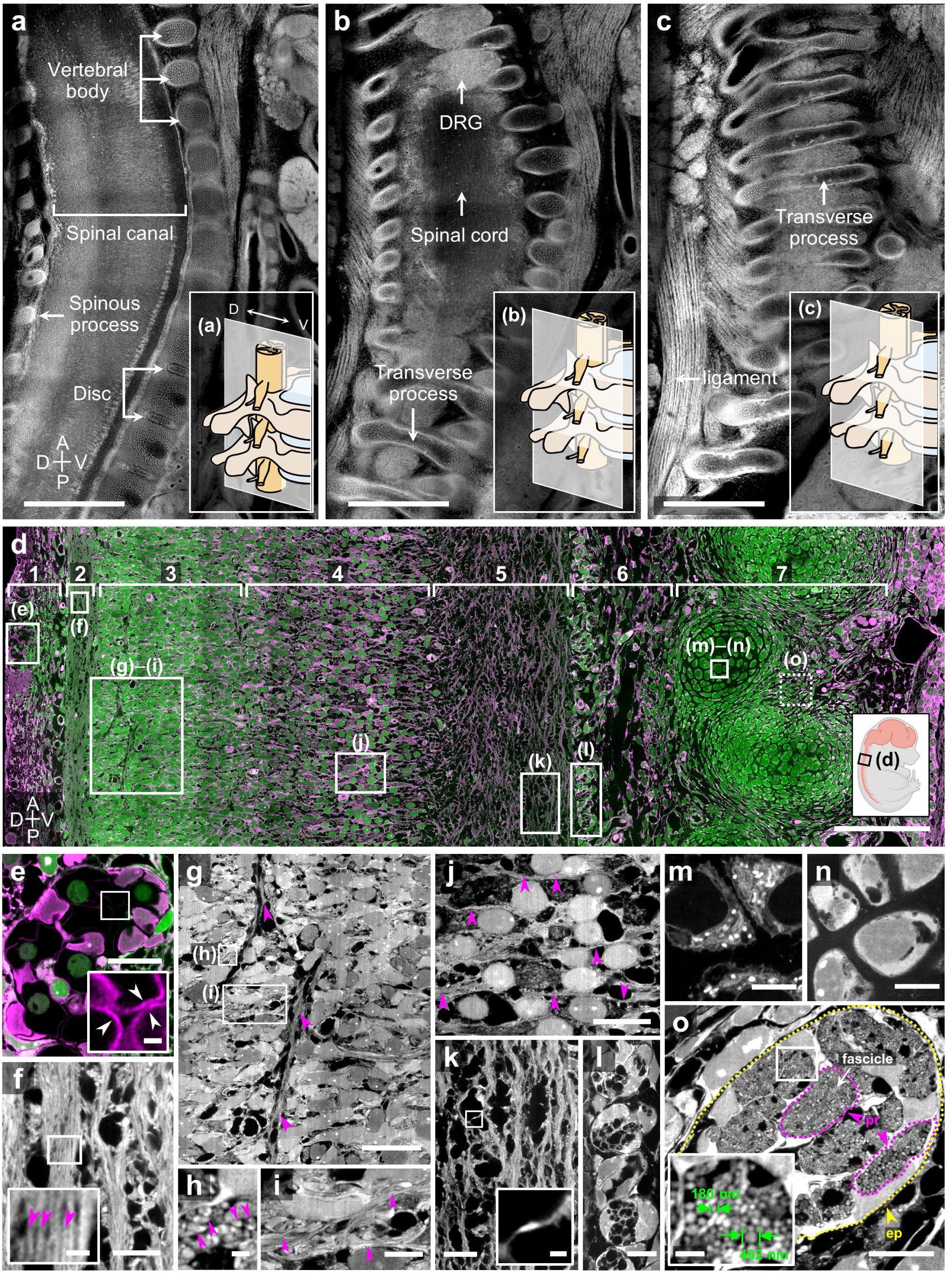
Whole-body ExM imaging of the spine and spinal cord of mouse embryos. (**a**–**c**) Confocal microscopy images of the cervical spine and spinal cord regions taken sequentially from the medial to lateral regions of an expanded E15.5 mouse embryo slice. (**d**) Mosaic image of the thoracic spine and spinal cord regions of the E15.5 mouse embryo after expansion. Green, Alexa Fluor 488 NHS ester; Magenta, ATTO 647N NHS ester. (**e**–**o**) Magnified views of the boxed region in **d** showing detailed morphologies of each layer. (**e**) Blood vessels and red blood cells found in the spinal meninges. The inset displays the magnified view of the boxed region. Arrowheads, membranes of RBCs. (**f**) Axon bundles aligned along the longitudinal axis. The inset displays the magnified view of the boxed region. Magenta arrowheads, single axon fibers. (**g**) Nuclei and axons aligned along the dorsoventral axis. Magenta arrowheads, single axon fibers aligned along the longitudinal axes. (**h**–**i**) Magnified view of the boxed regions in **g** showing axon bundles aligned along the (**h**) horizontal and (**i**) dorsoventral axes. Magenta arrowheads, single axon fibers. (**j**) The region between the gray matter and the white matter of the spinal cord containing fewer nuclei and axons aligned along the longitudinal axis. Magenta arrowheads, single axon fibers. (**k**) Spinal white matter containing thick axon bundles aligned along the longitudinal axis. The inset displays the magnified view of the boxed region. (**l**) A unique membrane organized into a single line found in the outermost layer of the spinal cord. (**m**–**n**) Two types of cells inside the vertebrae showing distinctive staining patterns. (**o**) Cross-sections of the spinal nerves next to the vertebra. Yellow dotted lines, epineurium; magenta dotted lines, perineurium; green, axon diameter. Scale bars: (**a**–**c**) 1 mm, (**d**) 100 μm, (**e**) 10 μm, inset of (**e**) 1 μm, (**f**) 10 μm, inset of (**f**) 1 μm, (**g**) 20 μm, (**h**) 1 μm, (**i**) 5 μm, (**j**–**k**) 10 μm, inset of (**k**) 1 μm, (**l**) 10 μm, (**m**–**n**) 5 μm, (**o**) 10 μm, and inset of (**o**) 1 μm. All length scales are presented in pre- expansion dimensions.

The outermost layer (layer 1 in **Fig. 5d**) was attached directly to the spinal cord and contained connective tissues and vessels, which is consistent with those of previous observations on the spinal meninges.^62^ In this layer as well, the high ATTO 647N staining intensities enabled the identification of RBCs and blood vessels (**Fig. 5e**). The first layer of the spinal cord (layer 2 in **Fig. 5d**), which is closest to the meninges, did not have any nuclei. Instead, axon bundles aligned along the longitudinal axis were found. High-resolution of whole-body ExM enabled the visualization of individual axons in the axon bundles (**Fig. 5f**). In the second layer of the spinal cord (layer 3 in **Fig. 5d**), which is considered the gray matter, a large number of nuclei were found, and axons aligned along a dorsoventral axis were observed (**Fig. 5g**). Relatively sparse axon bundles aligned along the horizontal and dorsoventral axes were also found (**Fig. 5h**–**i**). The third layer of the spinal cord (layer 4 in **Fig. 5d**) contained fewer nuclei; this layer showed higher ATTO 647N signals than the second layer, indicating that this layer contained more cellular organelles (**Fig. 5j**). The fourth layer of the spinal cord (layer 5 in **Fig. 5d**), which is considered the white matter, rarely contained nuclei. Instead, thick axon bundles aligned along the longitudinal axis were found (**Fig. 5k**). In layer 6 (**Fig. 5d**), a uniquely shaped membrane organized in a single longitudinal line was found, and loose reticular connective tissues and vessels were found between the membrane and the vertebrae (**Fig. 5l**). Inside the vertebrae (layer 7 in **Fig. 5d**), two types of cells showing distinctive staining patterns were found, probably due to the differences in the chondrocyte maturation and ossification center development stages (**Fig. 5m**–**n**).

Interestingly, the ultrastructures of the spinal nerves were visualized. From an image acquired at a z-plane different from the image shown in **Fig. 5d**, cross-sections of the spinal nerves were observed next to the vertebra. The characteristic structures of the nerves, including the epineurium around the entire nerve, fascicles, perineurium around fascicles, and individual axon fibers, were shown in the Alexa Fluor 488 channel (**Fig. 5o**). Consistent with previous studies, the axon diameters showed a broad distribution ranging from 150 nm to a few microns.^63^ Prior studies have reported that axon diameter and its distribution are closely related to neuronal function, energy capacity, and information rate, and axon morphology is modulated by the local environment. In those studies, high-resolution 3D imaging of axon bundles and their surroundings is required to fully reconstruct the volume fractions of the respective CNS compartments and thereby attain a robust description of single-axon structure and function.^64^ TEM imaging provides an adequate resolution for the quantitative analysis of the g-ratio, axon diameter, and axon diameter distribution, whereas micro-CT technologies have enabled investigations into axonal morphologies and projections over long distances with their mm-scale coverage.^64, 65^ However, whole-body ExM not only enables the 3D measurement of axonal parameters and trajectory variations over a large scale but also the extraction of the detailed information about the local environment, including volume fractions and the morphologies of cells, vacuoles, vessels, and even organelles, such as mitochondria.

### Visualization of neuronal structures on the face of mouse embryos using whole-body ExM

We then tested the use of whole-body ExM to image neuronal structures over a large volume. An E13.5 mouse embryo that had been sectioned in half along the sagittal plane was stained with an antibody against TUJ1, which stains all axons in the brain and the face.^9^ The embryo slice was then subjected to the whole-body ExM procedure. After expansion, we examined the structures labeled with Alexa Fluor 488 and an anti-TUJ1 antibody in the head of the embryo (**Fig. 6a**). In the Alexa Fluor 488 channel, the trigeminal ganglion and three axon bundle branches emerging from the trigeminal ganglion were apparent (**Supplementary Video 18**). When the expanded trigeminal ganglion was imaged with a high NA objective, the individual axons were resolved, as shown in **Fig. 6b**,**c**. In addition, ciliary structures of choroid plexus epithelium with a width of 150 nm were resolved, as shown in **Fig. 6d**. After the whole-body ExM procedure, the fluorescence signals of antibodies showed signal intensities high enough for nanoscale imaging. In the midbrain, the characteristically aligned axons were visualized in the antibody channel after expansion (**Fig. 6e**). Between the aligned axons, other aligned fibrous structures that were not stained with the TUJ1 antibody were shown in the Alexa Fluor 488 channel (**Fig. 6f**–**g**). More studies would be needed to identify the identities of these structures, but they would be the stem cells with their characteristic elongated structures.

**Fig. 6:**
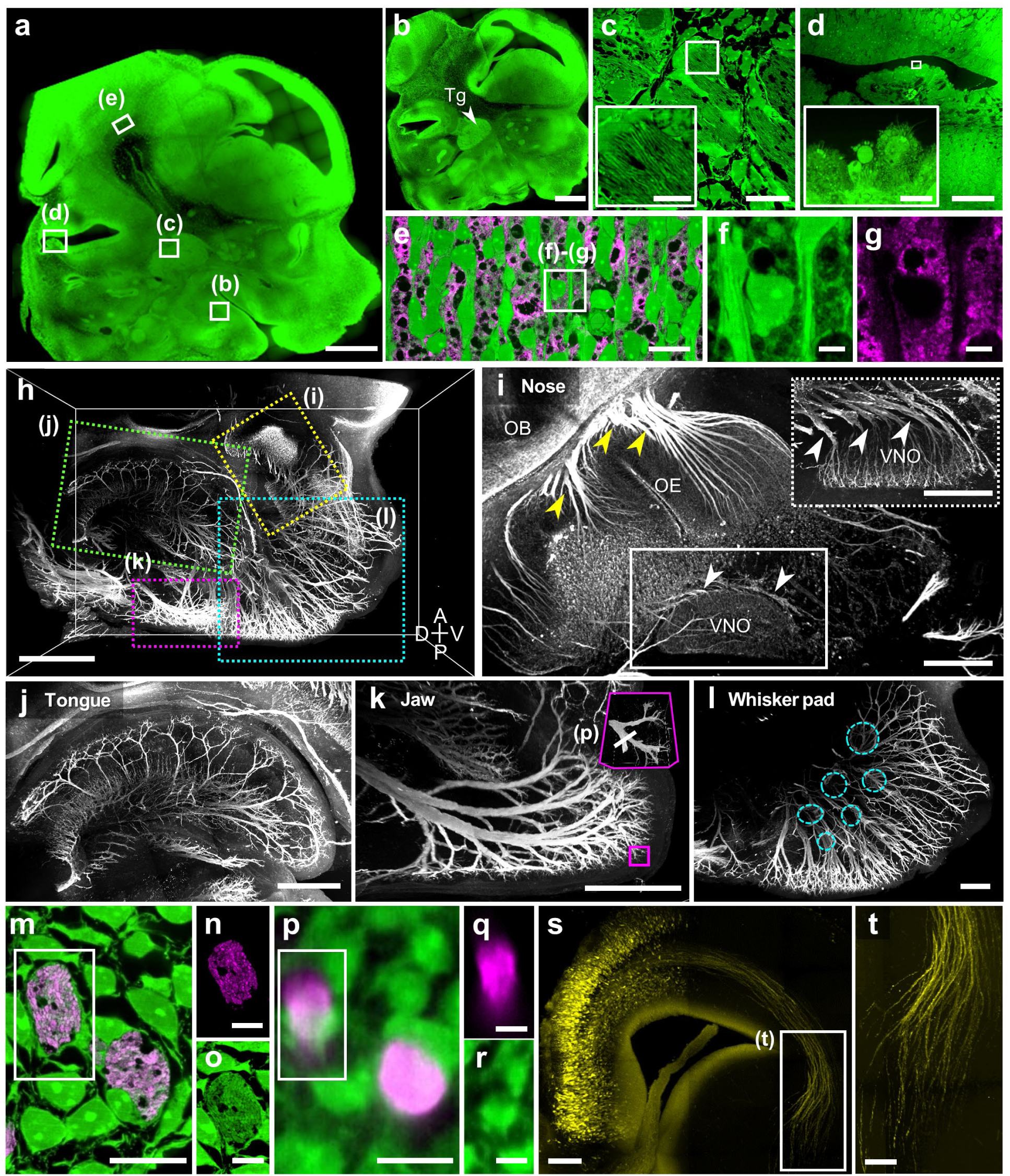
Whole-body ExM imaging of the craniofacial part of mouse embryos. (**a**) Confocal microscopy image of the head of the E13.5 mouse embryo after expansion. Green, Alexa Fluor 488 NHS ester. (**b**) Confocal microscopy image of the specimen shown in **a** acquired at a different height. Tg, trigeminal ganglion. (**c**–**g**) Confocal microscopy image of an expanded, E13.5 half mouse embryo specimen co-stained with anti-TUJ1 (magenta) and Alexa Fluor 488 NHS ester (green). (**c**) Trigeminal ganglion, (**d**) choroid plexus, and (**e**) midbrain. The insets in **c** and **d** display the magnified view of each boxed region. (**f**–**g**) Respective individual channels of the boxed region in **e**. **f**, Alexa Fluor 488 NHS ester staining; **g**, anti-TUJ1 staining. (**h**–**l**) Confocal microscopy images of an expanded mouse embryo with anti-TUJ1 staining. (**h**) TUJ-1 stained nerve networks in facial parts. Yellow dotted box, nose; green dotted box, tongue; magenta dotted box, jaw; and cyan dotted box, whisker pad. (**i**) Nerves network in the nose. The inset (upper right) displays the boxed region (middle) acquired at different heights from the bottom of the specimen and shows details of the vomeronasal nerves in the vomeronasal organ. Yellow arrowheads, main olfactory nerves; white arrowheads, vomeronasal nerves. OB, olfactory bulb; and OE, olfactory epithelium; VNO, vomeronasal organ. (**j**) Nerve networks with lingual, chorda tympani, and hypoglossal nerves inside the tongue. (**k**) Mental nerves in the lower jaw. The inset displays a magnified view of the boxed region with a single branched nerve bundle. (**l**) Infraorbital nerves of the maxillary branch in the whisker pad. Cyan dotted circle, free nerve endings. (**m**–**o**) Confocal microscopy images of a whole-body ExM-processed mouse embryo co-stained with Alexa Fluor 488 NHS ester (green) and anti-TUJ1 (magenta) with its respective individual channels of the boxed region. **n**, anti-TUJ1 staining; **o**, Alexa Fluor 488 NHS ester staining. (**p**–**r**) Confocal microscopy images of a BABB-cleared mouse embryo co-stained with Alexa Fluor 488 NHS ester (green) and anti-TUJ1 (magenta) with its respective individual channels of the boxed region. **p**, anti-TUJ1 staining; **r**, Alexa Fluor 488 NHS ester staining. (**s**) Confocal microscopy image of RFP-positive neurons (yellow) in a whole-body ExM-processed mouse embryonic brain. (**t**) Magnified view of the boxed region in **s**. Electroporated RFP-positive neurons show axonal projections in the developing brain. Scale bars: (**a**) 1 mm, (**b**) 1 mm, (**c**) 10 μm, inset of (**c**) 1 μm, (**d**) 50 μm, inset of (**d**) 5 μm, (**e**) 10 μm, (**f**–**g**) 2 μm, (**h**) 500 μm, (**i**–**l**) 200 μm, (**m**) 10 μm, (**n**–**o**) 5 μm, (**p**) 10 μm, (**q**–**r**) 5 μm, (**s**) 100 μm, and (**t**) 50 μm. All length scales are presented in pre-expansion dimensions.

We further studied neuronal structures on the face. ExM imaging of facial neurons has not been demonstrated before, as this requires the uniform expansion of the skull. We used a 10× objective (NA 0.45) and imaged a volume spanning 2 mm by 2 mm by 3 mm with a resolution of 152 nm within 8 h. As shown in **Fig. 6h**, such imaging visualized all axons in the face, including in the nose, tongue, jaw, and whisker pad. In the nose, axons of the main olfactory nerve in the olfactory epithelium (OE) and olfactory bulbs (OB) were clearly shown (**Fig. 6i**; see **Supplementary Video 19** for a 3D-rendered video). Below the OE, the axons of the vomeronasal organ (VNO) were also shown. In a different z-plane, the characteristic shape of the vomeronasal nerves was clearly visualized. In the tongue, the lingual, chorda tympani, and hypoglossal nerves inside the tongue were shown (**Fig. 6j**). In the jaw, mental nerves were clearly shown (**Fig. 6k**). In the whisker pad, infraorbital nerves were observed (**Fig. 6l**). When the axons were imaged with a high NA objective (40×, NA 1.15), individual axons consisting of the thick axon bundles shown in **Fig. 6k** were clearly resolved in both anti-TUJ1 and Alexa Fluor 488 channels, as shown in **Fig. 6m**–**o** (see **Supplementary Video 19** for a 3D-rendered video). The axon bundles were made of more than 100 axons with a diameter of 400 nm. Even without the anti-TUJ1 channel, the characteristic bundle structures of the axons enabled the identification of axon bundles from surrounding cellular structures in the Alexa Fluor 488 channel. However, in the unexpanded but cleared mouse embryos, individual axons were not resolved both in the anti-TUJ1 and Alexa Fluor 488 channels due to the limited axial resolution, as shown in **Fig. 6p**–**r**. We also tested another antibody, which was an anti-TH antibody, with whole-body ExM. TH staining visualized neuronal projections in the spinal cords and internal organs (**Supplementary Fig**. **21**). In addition to the compatibility between whole-body ExM and antibody staining, we tested whether the FP signals were retained after the whole-body ExM process for mouse embryos. We performed *in utero* electroporation with pCAG-RFP plasmid into E13.5 mouse embryonic brain to express RFP in the projection neurons. At E16.5, this mouse embryonic brain was fixed and processed for the whole-body ExM protocol. After expansion, the RFP signals were highly retained, enabling the visualization of projection neurons in the brain (**Fig. 6s**,**t**). In a separate experiment, we also confirmed that GFP signals were highly retained after the whole-body ExM procedure (data not shown). The compatibility of FPs with the whole-body ExM procedure makes the whole-body ExM a highly attractive tool for studying neuronal projections.

Finally, we asked whether specific cellular structures, such as axon bundles shown in **Fig. 6m**–**o**, can be identified from non-specific protein label channels, such as Alexa Fluor 488 or ATTO 647N channels, without the need to use antibodies using machine learning (ML). We trained a neural network with z-stack data sets that have two channels: the Alexa Fluor 488 and anti-TUJ1. The anti-TUJ1 channel was used as the ground truth of axon bundles and tested to determine whether the neural network could identify axon bundles from Alexa Fluor 488 channels. We tested both 2D and 3D images as ground truth learning data and found that 3D data showed much higher accuracy, as reported in the previous studies on the application of ML to images.^66, 67^ After the training, the neural network successfully marked axon bundles from Alexa Fluor 488 channels, as shown in **Supplementary Fig**. **22**. Considering the rapid staining speed of fluorophore NHS esters, the identification of specific structures from such pan-molecular staining would be a highly useful tool in the study of neuronal projections or morphologies of cellular organelles over a large volume.

## DISCUSSION

In this work, we demonstrated a protocol that enables three-dimensional super-resolution imaging of major anatomical landmarks, proteins, and FPs in whole larval zebrafish and mouse embryos *via* expansion microscopy. The key ideas were the post-digestion EDTA treatment to completely decalcify hard body parts to enable bone expansion and collagenase mixture use to homogenize the digestion speed to prevent crack formation during the digestion process. By using this process, we visualized the nanoscale details of the zebrafish larvae’s and mouse embryos’ anatomical structures from head to tail. In this work, we illustrated that the fluorescence signals of immunostained zebrafish larvae and mouse embryos were retained. Whole-body ExM can be directly used to image the nanoscale details of target proteins that have been studied without expansion. The compatibility of whole-body ExM with FPs makes it highly attractive for studying the morphological changes of organs and cellular organelles, as well as changes in the long processes of specific neurons, with a 60 nm resolution in transgenic models. For example, when applied to a SLICK (Single-neuron labeling with inducible Cre-mediated knockout) mouse line,^68^ whole-body ExM would visualize the distribution and morphologies of mitochondria, as well as their interactions with other organelles, in specific neurons from the cortical layers to the spine over a centimeter scale. Such super-resolution imaging would be highly useful in understanding the molecular mechanism of mitochondria biogenesis or lipid synthesis.^69^ In addition, *Fzf2*-Cre driver lines crossed with reporter lines would allow high-resolution tracing of the cerebrospinal tracts during embryonic development.^70^ When combined with state-of-the-art multiplexed cell labeling techniques, such as Confetti mouse^71^ or Brainbow-based mice,^72–74^ the study early cells’ clonal expansion of early cells over whole embryos would be possible.

When implementing whole-body ExM, considerations regarding imaging time and systems for the expanded whole mouse embryo are needed. Commercially available high NA objectives with a long working distance (e.g., 25× NA 0.95, WD 8 mm)^75^ would give an effective resolution of 67 nm when used to image expanded specimens. Using this objective, the imaging of a fourfold expanded whole 6 dpf zebrafish larva and half of an E13.5 mouse embryo would take two days and three months, respectively. Depending on the required resolution, expanded specimens could be reversibly shrunk in a buffer containing salt before being imaged.^76^ Completely imaging a twofold expanded half- mouse embryo with the same objective at an effective resolution of 134 nm could require 11 days. This resolution is high enough to resolve the 3D morphologies of cellular organelles, such as mitochondria, and individual axons in the axon bundles. The use of recently developed microscopy systems that can image a larger field of view could reduce the overall imaging time by half.^77^ Further reduction in imaging time could be achieved by using fast-evolving light-sheet microscopy techniques, which can acquire high-resolution images at a higher throughput.^31, 78^

Whole-body ExM could be improved in multiple ways. First, whole-body ExM could be used to visualize the sub-10 nm details of protein structures over the entire organism. Recently, expansion factors larger than fourfold, such as 10- or even 20-fold, have been demonstrated using different hydrogels or through multi-round expansion.^17, 24, 28, 34, 79^ Once combined with such more-than-fourfold expansion techniques, whole-body ExM would enable the study of the entire organism at a molecular resolution. Second, large Stoke’s shift fluorophores or spectral unmixing could be used to label multiple proteins, along with hydrophilic/hydrophobic fluorophores. Recently developed blind unmixing techniques, such as PICASSO, would enable the use of spectrally overlapping fluorophores for antibody staining; even fluorophore NHS esters are used in a similar spectral range.^80^ Most importantly, whole-body ExM could be combined with tissue clearing techniques that enable the rapid immunostaining of thick tissues. Recently, multiple technological breakthroughs, such as delipidation,^9, 81^ the use of nanobodies,^5^ and the optimization of staining conditions depending on antigen densities, fluorophore charges, and temperatures,^10^ have accelerated immunostaining. Based on this, the immunostaining of mm-thick specimens,^6, 11, 82^ even whole mouse embryos,^9^ has been achieved. Other rapid staining techniques that strongly fix specimens and accelerate antibody diffusion by applying electric fields could also be used to stain whole mouse embryos.^83–85^ The compatibility of these techniques with the whole-body ExM protocol should also be studied.

The use of machine learning to identify specific structures from fluorophore NHS ester images of expanded specimens would be the subject of further study. Recently, machine learning has been applied to data from label-free optical microscopy imaging,^86, 87^ vibrational imaging of expanded specimens,^88^ and EM imaging^89–92^ for cellular structure segmentation and reconstruction. In applying ML to image data, the most critical part is acquiring a large amount of labeled data, which can be used as ground truth training. As whole-body ExM is compatible with antibody labeling and FPs, it eliminates the need for the manual annotation of specific structures. As observed in various examples, including mitochondria and RBCs in the mouse embryo heart, the simultaneous use of hydrophilic and hydrophobic fluorophores provides chemical information on structures, in addition to the morphological data provided by a single fluorophore. Such dual-labeled data would enable more precise structural identification when used with machine learning. The range of structures that can be identified from expanded whole embryos stained with pan-molecular techniques would be the subject of further study.^26, 27, 29^

## METHODS

### Materials

All chemicals and antibodies were obtained from commercial suppliers, and detailed information is provided in **Supplementary Table 1**. The concentrations of the labeling agents, including antibodies, fluorophore NHS esters, and 4′,6-diamidino-2-phenylindole (DAPI), are listed in **Supplementary Table 2**.

### Ethical regulations

All experimental methods involving mice and zebrafish were approved by the Korea Advanced Institute of Science and Technology Institutional Animal Care and Use Committee (KAIST-IACUC), the Korea Institute of Science and Technology Institutional Animal Care and Use Committee (KIST-IACUC), and the Korea Research Institute of Bioscience and Biotechnology Institutional Animal Care and Use Committee (KRIBB-IACUC). Experiments related to ExFISH labeling in zebrafish were conducted in accordance with the US National Institutes of Health Guide for the Care and Use of Laboratory Animals and approved by the Massachusetts Institute of Technology Committee on Animal Care and Biosafety Committee.

### Primary and secondary antibody preparation

The following primary antibodies were used: goat anti-FITC Alexa Fluor 488, rabbit anti-Homer1, rabbit anti-SLC2A1, rabbit anti-RFP, mouse anti-Bassoon, mouse anti-HuC/D, mouse anti-TUJ1, mouse anti-TH, and rat anti-MBP. The following fluorophore-conjugated secondary antibodies were used: goat anti-mouse CF 405S, goat anti-mouse Alexa Fluor 488, goat anti-mouse Alexa Fluor 546, goat anti-rat Alexa Fluor 488, goat anti-rabbit Alexa Fluor 488, goat anti-rabbit Alexa Fluor 546, goat anti-rabbit CF 633, and donkey anti-mouse Alexa Fluor 546. Detailed information about the antibodies used is provided in **Supplementary Table 2**.

### Mouse organ slice preparation

C57BL/6J mice aged 6–8 weeks were used for organ harvesting. After anesthetization with isoflurane, the mice were transcardially washed with ice-cold 1× phosphate buffered saline (PBS) supplemented with 10 U/mL heparin, followed by perfusion with ice-cold 4% paraformaldehyde (PFA) in 1× PBS. Mouse organs were harvested and fixed in 4% PFA in 1× PBS at 4 °C for 2–6 h. Fixed brains were sliced to a thickness of 150 µm with a vibratome (VT1000S; Leica, Wetzlar, Germany). Other organs were embedded in 4% (w/w) low-gelling-temperature agarose to prepare agarose blocks. Agarose blocks were then sliced to a thickness of 500–1000 µm with a vibratome. After agarose residue was removed from the cut slices, organ slices were stored in 1× PBS with 0.1 M glycine and 0.01% (w/w) sodium azide at 4 °C before use.

### Mouse brain staining and expansion

Samples were blocked and permeabilized with normal goat serum (NGS) blocking buffer (5% [w/w] NGS, 0.2% [w/w] Triton X-100, 1× PBS) for 1–3 h. For brain slices that were imaged for actin, the permeabilized brains were stained with fluorescein phalloidin in 0.1% Triton X-100 in 1× PBS (0.1% PBST) overnight at 4 °C followed by washing in 0.1% PBST three times for 30 min each time. Permeabilized samples were then incubated with primary antibodies in NGS blocking buffer for at least 6 h at 4 °C, followed by washing in 0.1% PBST three times for 30 min each time. Secondary antibodies diluted in NGS blocking buffer were then added for at least 6 h at 4 °C followed by washing in 0.1% PBST three times for 30 min each time. Then, 6- ((acryloyl)amino)hexanoic acid (AcX) treatment, gelation, digestion, and expansion were performed as previously described in the proExM protocol.^22^ For brain slices co-stained with fluorophore NHS ester, either pre-gelation or post-gelation NHS ester staining was performed. Fluorophore NHS ester staining is described in the following section.

### Fluorophore NHS ester staining and expansion of mouse organs

**i) Pre-gelation staining of mouse organs**. Organ slices were permeabilized with 0.1% PBST for 3 h. For the pre-gelation staining of fluorophore NHS ester, organ slices were first stained with the diluted fluorophore NHS ester in 0.1% PBST for 6 h at 4 °C, followed by washing in 0.1% PBST three times for 30 min each time at room temperature (RT; 20–25 °C). Next, stained organ slices were treated with 0.1 mg/mL AcX in 1× PBS for 6 h at RT. AcX-treated organ slices were washed in 1× PBS three times for 30 min each time at RT with gentle shaking. For gelation, organ slices were first incubated with gelation solution (8.625% [w/w] sodium acrylate, 2.5% [w/w] acrylamide, 0.15% [w/w] *N,N*′-methylenebis(acrylamide) [BIS], 2 M NaCl, 0.2% [w/w] ammonium persulfate [APS], 0.2% [v/v] tetramethylethylenediamine [TEMED], and 0.01% [w/w] 4-hydroxy-2,2,6,6-tetramethylpiperidin-1-oxyl [H-TEMPO] in 1× PBS) at 4 °C twice for 30 min each time, followed by incubation at 37 °C for 2 h. Gelled organ slices were then homogenized with 8 U/mL proteinase K (proK) in digestion buffer (1 mM ethylenediaminetetraacetic acid [EDTA], 50 mM Tris-HCl [pH 8.0], and 0.5% Triton X-100, 1 M NaCl) 3–4 times at 37 °C for 12 h each time then 2–3 times at 60 °C for 6 h each time. Complete digestion was confirmed when the digested organ–gel composite was isotropically expanded by approximately 1.5-fold without distortion on its surface.
**ii) Inter-digestion staining of mouse organs**. For inter-digestion staining of organs with the fluorophore NHS ester, permeabilized organ slices were first treated with AcX, as described above. Then, gelation of organs was performed, followed by a single round of digestion with proK for 4–12 h at 37 °C. Then, gelled organ slices were stained with diluted fluorophore NHS ester in 0.1% PBST for at least 6 h at 4 °C, followed by washing in 0.1% PBST three times at RT for 30 min each time. After staining, further digestion was performed, as described above, until complete digestion of gelled organ slices was achieved.
**iii) Post-digestion staining of mouse organs**. For post-digestion staining of organs with the fluorophore NHS ester, permeabilization, AcX treatment, gelation, and digestion were conducted as previously described. Fully digested samples were stained with diluted fluorophore NHS ester in 0.1% PBST for at least 6 h at 4 °C, followed by washing in 0.1% PBST three times at RT for 30 min each time.
**iv) DAPI staining and expansion**. For all three staining methods, fully digested NHS-ester-labeled gels were stained overnight with DAPI diluted in 1× PBS at 4 °C then washed with 1× PBS three times for 30 min each time at 4 °C. After DAPI staining, the gels were washed with an excess volume of deionized water for expansion until the size of the gels plateaued.

### *In vivo* mitochondria labeling

To label mitochondria in parvalbumin (PV)-positive neurons, Cre-dependent adeno-associated virus (AAV) expressing mScarlet followed by mitochondrial matrix-targeting sequence (from cytochrome C subunit VIII) was injected into the PV-Cre mouse line (B6.129P2-Pvalb^tm1(cre)Arbr^/J), as follows. The mice were anesthetized with isoflurane and placed in a stereotaxic frame. Then, 100 nL of 10:1 cocktail of AAV-Jx- synaptophysin-mVenus-T2A-mito-mScarlet was injected into the subthalamic nucleus (STN, AP: +1.62 mm, ML: 1.65 mm, DV: -4.45, -4.55 and -4.65 mm) at a speed of 40 nL/min. Ten days after AAV injection, mice were transcardially perfused, and the harvested brains were fixed with 4% PFA, as mentioned above.

### Zebrafish sample preparation

Wild-type, Casper (*mitfa^w2/w2^; mpv17^a9/a9^*), and *flk-egfp* (=*Tg*(*kdrl*:*EGFP*)) larval zebrafish (*Danio rerio*) aged 3–12 d post-fertilization were used in this study. Larvae were first fixed with fixative at 4 °C for 6 h. Fixed specimens were then kept in 0.1% PBST at 4 °C for 6 h for washing and permeabilization.

### Whole-body ExM of larval zebrafish

(1) **Gelation**. Fixed and permeabilized specimens were incubated in 0.1 mg/mL AcX in 0.1% PBST three times for 6 h each time at 4 °C then washed with 0.1% PBST three times for 30 min each time at 4 °C. The specimens were then incubated with a gelation solution (8.625% [w/w] sodium acrylate, 2.5% [w/w] acrylamide, 0.15% [w/w] BIS, 0.2% [w/w] 2,2’-azobis[2-(2-imidazolin-2-yl)propane] dihydrochloride [VA-044], 0.05% [w/w] Triton X-100, 2 M NaCl, 1× PBS) three times for 6 h each time at 4 °C with gentle shaking. After incubation, each zebrafish was placed between two pieces of #1 cover glass filled with a freshly prepared gelation solution and incubated at 45 °C for 4 h in a humidified, nitrogen-filled chamber.
(2) **Pre-digestion**. After the careful removal of the upper glasses, the cover glasses with attached hydrogels were rinsed with 0.1% PBST multiple times. The excess gel around the specimens was removed using a razor blade, and the resulting rectangle-shaped hydrogels were detached from the cover glasses with a wet paintbrush. The gels were then transferred into a 12-well plate for pre-digestion in 1.5 mL of pre-warmed digestion buffer supplemented with 16 U/mL proK, and incubated three times for 6–12 h each time at 37 °C with gentle shaking. Finally, gels were incubated in staining base buffer (0.1% Triton X-100, 1 M NaCl, 1× PBS) three times for 30 min each time at 4 °C for washing.
(3) **Inter-digestion staining of fluorophore NHS ester**. For NHS-ester staining of zebrafish, the fluorophore NHS ester stock solutions (10 mg/mL in anhydrous DMSO) were stored at -20 °C and then diluted to 1:1000 in cold staining base buffer to prepare the staining solution. Following pre-digestion, the gels were incubated with freshly prepared staining solution for at least 12 h at 4 °C with gentle shaking and then washed with staining base buffer three times for 30 min each time at 4 °C.
(4) **Digestion and post-digestion decalcification**. For further digestion, gels were treated with 16 U/mL proK in digestion buffer at 37 °C 8–12 times for 6 h each time. To completely remove calcium ions that may have been released from digested bone, cartilage, and calcified tissues, fully digested samples were treated with an excess volume of decalcification solution (0.3 M EDTA, 1 M NaCl, 0.1% Triton X-100, pH 8.0) three times for 30 min each time at RT with gentle shaking.
(5) **Expansion**. Decalcified gels were serially washed with decreased concentrations of NaCl (1 M NaCl, 0.8 M NaCl, 0.6 M NaCl, 0.4 M NaCl, and 0.2 M NaCl) for 30 min each at RT. The gels were then washed with an excess volume of deionized water three times for at least 30 min each time at RT and then kept in fresh deionized water overnight at 4 °C for full expansion. Expanded specimens were stored in the dark before imaging.

### GFP imaging of whole zebrafish larvae

Transgenic *flk-egfp* zebrafish larvae at 3 dpf were dechlorinated and then fixed with 4% PFA in 1× PBS for 1 h at 4 °C. After fixation, the larvae were washed three times with 0.1% PBST. To test endogenous GFP signal retention after proteinase digestion, the larvae were processed according to the previously described whole-body ExM protocols. After 54 h of digestion, the specimens were washed with 0.1% PBST multiple times, expanded, and imaged with a confocal microscope.

### ExFISH of whole zebrafish larvae

The zebrafish larvae were processed and expanded according to the proExM and ExFISH protocols.^20, 22, 93^ LabelX was prepared according to the ExFISH protocols.^20, 93^ Briefly, zebrafish larvae were fixed using 4% PFA in 1× PBS for 20 min at 4 °C. After fixation, the larvae were washed twice with Modified Barth’s Saline (MBS) buffer at pH 6.0 and incubated overnight with 0.1 mg/mL of AcX in MBS buffer at RT. For ExFISH processing, incubation with AcX was followed by incubation with homemade LabelX in 3-(*N*- Morpholino)propanesulfonic acid (MOPS) buffer at pH 7.7 overnight at 37 °C. Zebrafish were transferred to a pre-gel solution and incubated overnight at 4 °C. Before gelation, zebrafish were transferred into fresh pre-gel solution containing H-TEMPO, TEMED, and APS in 94:2:2:2 pre-gel solution:H-TEMPO:TEMED:APS and incubated for 1 h at 4 °C before being polymerized in a gelation chamber for 2 h at 37 °C. ^90^ Sample homogenization was performed using 8 U/mL proK in a buffer containing 50 mM Tris-HCl (pH 8.0), 500 mM NaCl, and 40 mM CaCl2 for 10 h at 37 °C. For ExFISH, gelled samples were processed as described before using HCR-initiator-tagged FISH probes (Molecular Instruments Inc.) hybridization for *SNAP*-*25* and *β*-*actin* followed by hybridization chain reaction amplification with HCR hairpin stocks labeled with Alexa Fluor 546 and Alexa Fluor 647 fluorophores. ExFISH- processed samples were expanded in 0.05× saline sodium citrate (SSC).

### Mouse embryo sample preparation

Mouse embryos were isolated on days 11.5–15.5 of pregnancy in C57BL/6J mice and fixed with ice-cold fixative for 24–72 h. Fixed mouse embryos were embedded in 4–8% (w/w) low-gelling-temperature agarose and then sliced to a thickness of 500–1000 µm with a vibratome. After agarose residue was removed from the cut slices, embryo slices were stored in 1× PBS with 0.1 M glycine and 0.01% (w/w) sodium azide at 4 °C before use.

### *In utero* electroporation

The experimental setup and method were designed and performed based on the previously described protocol.^94^ In summary, time-pregnant CD1 mice were anesthetized at E13.5 for laparotomy. Uterine horns were exposed, and embryos were manipulated with approximately 1 µL of endotoxin-free plasmid (pCAG-RFP) with the fast green (0.01%) of the final concentration of 1.5 µg/uL and injected into the lateral ventricle of the embryos with the calibrated microcapillary in a Pneumatic PicoPump setup (PV820, WPI). Five transfer pulses with 43 V, pulse length 50 ms, pulse interval 450 ms, and decay rate 40 were delivered over the uterus on top of the embryo’s head with electrode puddles (Nepa21 TypeII, Nepa Gene). After the completion of electroporation, the uterus was placed back in the abdominal cavity, and the wound was stitched with a suture (W581 Mersilk, Ethicon). The mother was normally reared in the animal facility and euthanized after 3 d to collect embryos at E16.5 and fixed with 4% PFA, as mentioned above.

### Antibody staining of mouse embryos

Fixed half-embryo samples were blocked and permeabilized with NGS blocking buffer for 12 h. Permeabilized samples were then incubated with primary antibodies in NGS blocking buffer for 3 weeks at 4 °C. The antibodies were refreshed once after 24–48 h. Then, samples were washed with NGS blocking buffer 12 times for 30 min each time. Secondary antibodies diluted in NGS blocking buffer were then added for 10 d at 4 °C, followed by washing in NGS blocking buffer 12 times for 30 min each time.

### Whole-body ExM of mouse embryos

**i) Pre-gelation treatment**. Embryos were incubated in 0.1 mg/mL AcX in 0.1% PBST 10 times for 12 h each time at 4 °C, followed by washing with 1× PBS three times for 1 h each time at 4 °C. The specimens were then incubated with a gelation solution four times for 6 h each time at 4 °C with gentle shaking. After incubation, each mouse embryo was placed in the gelation frame made with multiple layers of slide glass, sealed with glue, and sandwiched by two pieces of #1 cover glass. Gelation was performed at 45 °C for 4 h in a humidified, nitrogen-filled chamber.
**ii) Pre-digestion**. After gelation, the excess gel around the specimens was removed using a razor blade. Cut gels were then incubated with a digestion buffer (1 mM EDTA, 50 mM Tris-HCl [pH 8.0], 0.5% [w/w] Triton X-100, and 2 M NaCl) supplemented with 16 U/mL proK for 12 h at 37 °C with gentle shaking. Finally, gels were incubated in staining base buffer (0.1% Triton X-100, 2 M NaCl, and 1× PBS) three times for 30 min each time at 4 °C for washing.
**iii) Inter-digestion staining of fluorophore NHS ester**. Following pre-digestion, the gels were incubated with freshly prepared NHS ester staining solution 10 times for 12 h each time at 4 °C with gentle shaking and then washed with staining base buffer 6 times for 1 h each time at 4 °C.
**iv) Post-digestion with collagenase cocktail and post-digestion decalcification**. NHS ester-stained gels were treated with collagenase cocktail (400 U/mL collagenase type I, 400 U/mL collagenase type II, and 400 U/mL collagenase type IV) in collagenase buffer (1× HBSS, 3 mM CaCl2) at 37 °C 1–2 times for 12 h each time. After then, samples were treated with 16 U/mL proK in digestion buffer at 37 °C 8–12 times for 12 h each time. To completely remove calcium ions, fully digested samples were treated with an excess volume of decalcification solution (0.3 M EDTA, 2 M NaCl, 0.1% Triton X-100, pH 8.0) 2–4 times for 6 h each time at 4 °C with gentle shaking.
**v) Expansion**. Decalcified gels were serially washed with decreased concentrations of NaCl (1 M NaCl, 0.5 M NaCl, and 0.1 M NaCl) supplemented with 0.3 M EDTA and 0.1% Triton X-100 for 1 h each time at RT. Gels were then washed with an excess amount of 1× PBS for around twofold expansion. If necessary, DAPI was stained after the sample treatment. For around 4.2-fold expansion, gels were washed with an excess volume of deionized water, and the deionized water was exchanged until the size of the gels plateaued.

### BABB clearing of mouse embryos

**i) Sample treatment and staining**. Either whole mouse embryos or mouse embryo slices were treated using the same experimental steps for BABB clearing. The fixed specimens were first treated with 0.2% PBST for 2 h. Then, the specimens were dehydrated with increased concentrations of ethyl alcohol in deionized water (30%, 50%, 70%, 90%, and 100%) for 1 h each at RT. Dehydrated specimens were then treated with hydrogen peroxide solution (made with 1/3 dilution of 30% hydrogen peroxide in pure ethyl alcohol) overnight at -20 °C. Then, the specimens were rehydrated with decreased concentrations of ethyl alcohol in deionized water (100%, 90%, 70%, 50%, and 30%) for 1 h each at RT. Following the rehydration, the specimens were permeabilized with NGS blocking buffer for 2 h. Specimens were then treated with primary antibodies diluted in NGS blocking buffer for 48 h at 4 °C, followed by washing with NGS blocking buffer 6 times for 1 h each time. Next, secondary antibodies diluted in NGS blocking buffer were treated for 48 h at 4 °C, followed by washing with NGS blocking buffer 6 times for 1 h each time. Antibody-labeled specimens were then treated with fluorophore NHS ester diluted in 0.2% PBST for 24 h at 4 °C, followed by washing with 0.1% PBST 6 times for 30 min each time. For fluorophore NHS ester labeling only, antibody staining step could be skipped.
**ii) Clearing**. Antibody or fluorophore NHS ester-labeled specimens were dehydrated with increased concentrations of ethyl alcohol in deionized water (30%, 50%, 70%, 90%, and 100%) for 1 h each at RT. The dehydrated specimens were then placed in a confocal dish, and the remaining ethyl alcohol around the specimens was completely removed by evaporation. Then, the specimens were incubated with a BABB solution consisting of benzyl alcohol and benzyl benzoate with a volume ratio of 1:2 for 1 h. The BABB solution was then replaced with a fresh BABB solution. After the specimens became transparent, imaging was performed with a confocal microscope. The imaging of the specimens was performed within 24 h after the BABB solution treatment.

### Determination of the expansion factor

The expansion factor for the mouse organs, zebrafish larvae, mouse embryo slices, and whole mouse embryos was quantified by measuring the gel sizes before and after expansion. Concurrently, the same structural landmarks were imaged pre- and post-expansion, and the expansion factor was calculated using these landmarks. For all experiments in this study, the expansion factor determined by landmarks was approximately 4.2-fold, which is coincident with the expansion factor determined by gels.

### Reproducibility

In this study, more than 100 zebrafish aged 3–12 dpf were used for protocol optimization and confirmation, and more than 30 zebrafish from this cohort were imaged for data acquisition. For mouse embryos, more than 80 embryos at embryonic stages 10.5–15.5 were used for protocol optimization and confirmation, and more than 30 mouse embryos from this cohort were imaged for data acquisition. All images in **Figs**. **1**–**6** and **Supplementary Figs**. **1**–**22** were acquired from at least 30 independent experiments.

### Sample mounting and imaging

For the imaging of expanded hydrogels, the hydrogels were attached to 48 × 60 mm cover glasses to prevent hydrogel drifting during imaging. To ensure firm attachment, the cover glasses were treated for 30 min with 0.01% poly-L-lysine and then washed with deionized water. The hydrogels were then placed on cover glasses and immediately imaged using confocal microscopy. Specimens were imaged using a Nikon Eclipse T*i*2-E (Tokyo, Japan) microscope with either a spinning disc confocal microscope (Dragonfly 200; Andor, Oxford Instruments, Abingdon, UK; CSU-XI; Yokogawa Electrical Corporation, Tokyo, Japan) equipped with a Zyla 4.2 sCMOS camera (Andor, Oxford Instruments) or a laser scanning confocal microscope (Nikon C2+, Nikon). The objectives used were 40× 1.15 NA water immersion lens, 10× 0.45 NA air lens, and 4× 0.2 NA air lens.

### Shading correction

A custom Python script was used to apply the BaSiC algorithm^95^ for flat-field estimation and shading correction on 3D image stacks in the Imaris file format (.ims). For each 3D image stack, ten image slices (2048 px × 2048 px each) were sampled evenly along the z-direction and resized to 512 px × 512 px. BaSiC algorithm with default parameters was applied to the combined list of downsampled slices from total 208 (13 × 16) stacks to estimate flat-field profile. The estimated flat-field profile was then resized to match the original image size. A single flat-field profile was used to correct shading in all image stacks with correction applied to each image slice along the z-directions of the image stack independently. Corrected image stacks were saved as .ims files for further processing, including multi-stack stitching and visualization in Imaris 9.5.

### Inference of axon morphology from fluorophore NHS ester-stained images using deep learning

First, we used a neural network to infer the axonal morphology from the fluorophore NHS ester-stained images. We used three-dimensional images from the facial part of a mouse embryo obtained with fluorophore NHS ester staining as the input and TUJ1 antibody staining as the network target. The training and testing pipelines are illustrated in **Supplementary Fig**. **22a**. We employed a U-Net architecture that consists of a 3D encoder, a 3D decoder, and four skip connections from the encoder to the decoder, as illustrated in **Supplementary Fig**. **22b**. In the 3D encoder, there were four encoder blocks. Each block consisted of a 3(*z*) × 3(*x*) × 3(*y*) convolutional layer, followed by a BatchNorm, a LeakyReLU, and a max pooling layer. The 1(*z*) × 2(*x*) × 2(*y*) max pooling layers were used, except for the second block, which used a 2(*z*) × 2(*x*) × 2(*y*) max pooling layer to increase the receptive field of the network along the z-axis. In the decoder, there were four decoder blocks, each of which contained a trilinear interpolation followed by a 3(*z*) × 3(*x*) × 3(*y*) convolutional layer, a BatchNorm, and a LeakyReLU. The skip connections linked low- and high-level features by adding feature maps. For the loss function, we used binary cross entropy loss. The Adam optimizer was used with a learning rate of 0.0003 and a batch size of two for training the network. The network was implemented using PyTorch. All computations were performed on a workstation with two Intel Xeon Silver 4214R CPUs, 256 GiB RAM, and an NVIDIA GeForce RTX 3090 GPU. For deep learning, we preprocessed z-stack images of a mouse embryo that was stained with fluorophore NHS ester and TUJ1 antibody. The input channel (fluorophore NHS ester labels) was normalized by subtracting the minimum pixel value and then dividing it by the 99th percentile value. The target channel (TUJ1 antibody label) was binarized to generate a axonal morphology mask. Then, we sampled 3D patches with a size of 2(*c*) × 80(*z*) × 416(*x*) × 416(*y*) from the volume as the training data. We augmented the data by flipping and rotating the images by a random multiple of 90°.

## ACKNOWLEDGEMENTS

This work was supported by the Samsung Research Funding & Incubation Center for Future Technology (SRFC- IT1702-09). This work was also supported by the National Research Foundation of Korea (NRF) grant funded by the Korea government (MSIT) (NRF-2021R1C1C1006642) and the Bio & Medical Technology Development Program of the National Research Foundation (NRF) & funded by the Korean government (MSIT) (NRF-2021M3A9I4026318). E.S.B. acknowledges for funding, NIH 1R01EB024261, Good Ventures, Open Philanthropy, Lisa Yang, John Doerr, HHMI, NIH Director’s Pioneer Award 1DP1NS087724, and U. S. Army Research Laboratory and the U. S. Army Research Office under contract/grant number W911NF1510548. We acknowledge Prof. Taeyun Ku and Mr. Young Seo Kim for their help with imaging. We also acknowledge Prof. Myunghwan Choi and Prof. Taejoon Kwon for their helpful discussions. We thank Ms. Su Yeon Kim and Dr. Jang Soo Yuk for their help with transgenic mouse experiments. Furthermore, we thank Dr. Yun-Mi Jeong and Dr. Hoi-Khoanh Giong for their help with the zebrafish experiments. We acknowledge Prof. Hyuk Wan Ko, Dr. Rose G. Long, Prof. Robert Parton, and Prof. Tanya T. Whitfield for their insightful discussions. Cartoons in **Fig. 1a** was created with BioRender.com.

## DATA AVAILABILITY

All experimental data is available upon request to the corresponding authors.

## CODE AVAILABILITY

Computational code related to shading correction and machine learning is available upon request to the corresponding authors.

## AUTHOR CONTRIBUTIONS

J.S., C.E.P., and I.C. performed the experiments and analyses. J.S., C.E.P., and I.C. wrote the paper and contributed to its editing. K.M. performed the expansion of the mouse organs. M.E. and S.H. performed machine learning analysis.

H.J. processed large images for fine stitching. H.-J.C. and E.-S.C. prepared zebrafish samples. A.K. prepared *in utero* electroporated mouse embryos. Y.C. analyzed the anatomical structures of expanded mouse organs. J.S.K., K.D.P., and E.E.J. performed ExFISH imaging of the whole larvae. D.-S.K. contributed to mouse embryo clearing and designed an experimental process with mouse embryos. S.-K.K. prepared *in vivo* mitochondria-labeled mouse brains.

J.K. supervised the processing of large images for fine stitching. J.L. analyzed the anatomical structures of expanded larvae. E.S.B. supervised ExFISH imaging of the whole larvae. Y.-G.Y. supervised the machine learning analysis. J.-B.C. supervised this work. All the authors commented on the manuscript.

