## Supplementary Information for "Nanoscale resolution imaging of the whole mouse embryos and larval zebrafish using expansion microscopy"

**Supplementary Figures 1–22**

**Supplementary Videos 1–19**

**Supplementary Notes 1–6**

**Supplementary Tables 1–2**

**References**

### SUPPLEMENTARY FIGURES

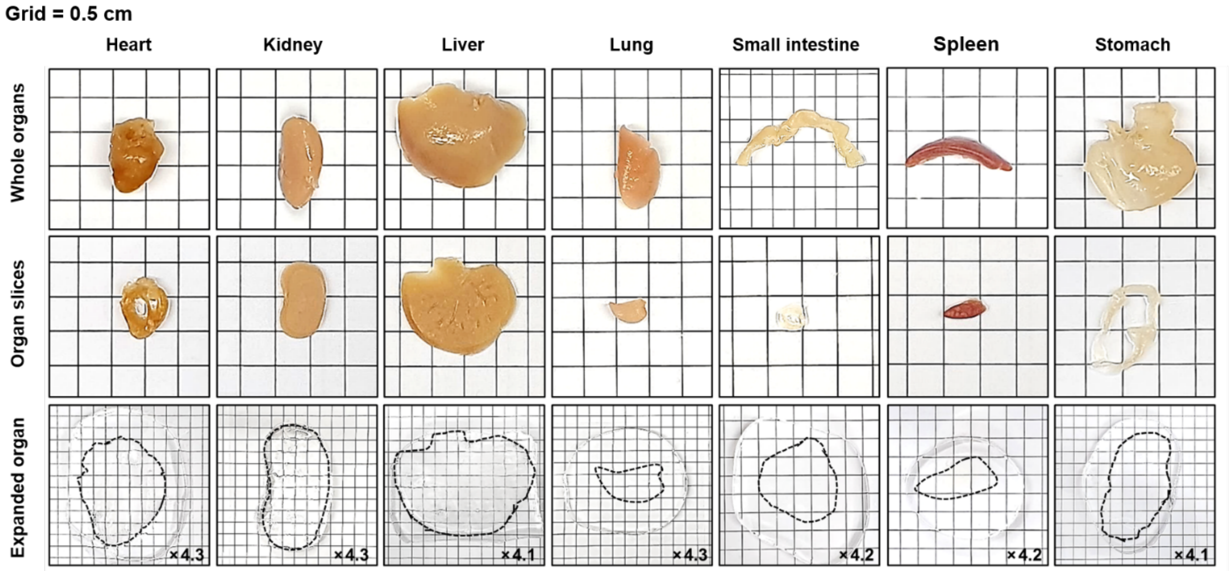


**Supplementary Fig. 1**: **Mouse organs before and after expansion**. The top row, photographs of whole mouse organs; the middle row, mouse organs sliced to a thickness of 1 mm; and the bottom row, mouse organs after repetitive proteinase K treatment and expanding about fourfold in deionized water. Black dotted lines indicate the outer boundaries of the expanded organ slices. Grid size: 0.5 cm.

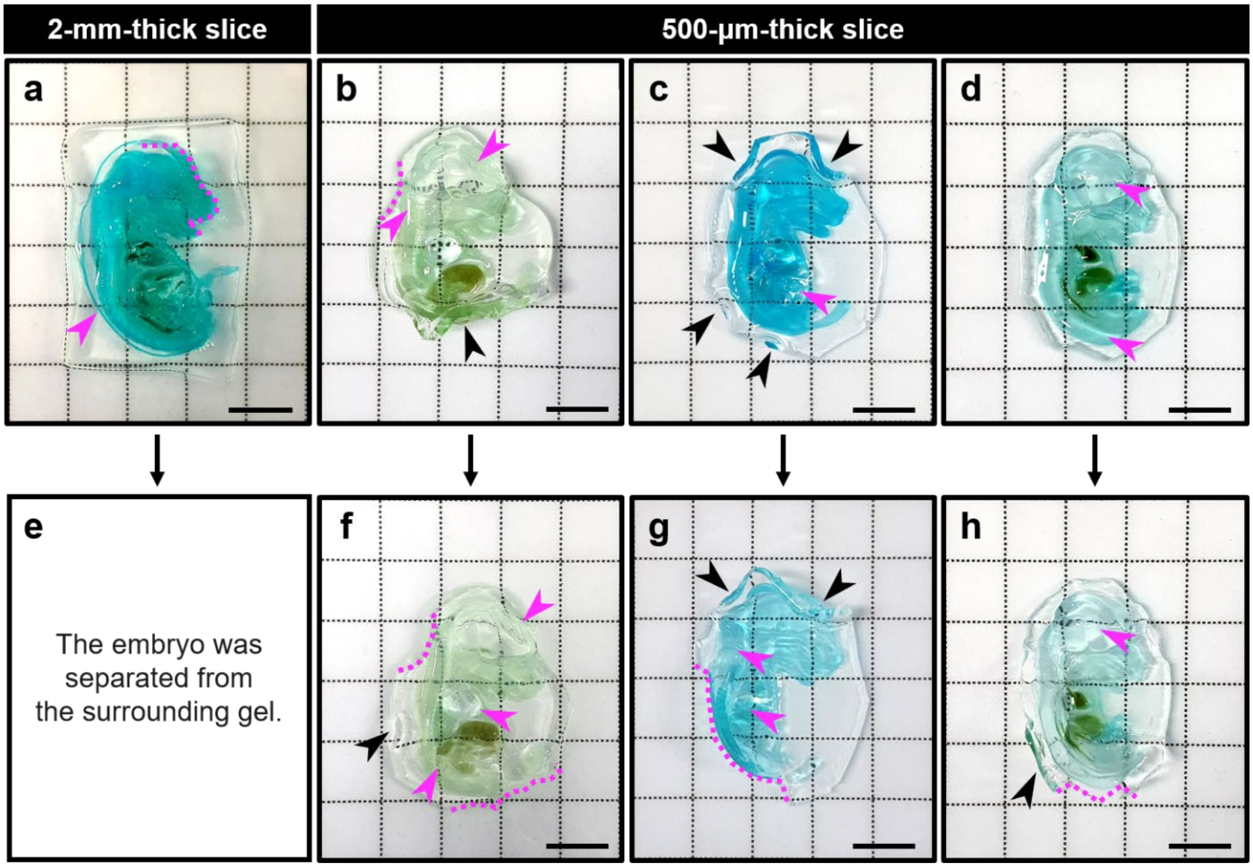


**Supplementary Fig. 2**: **Mouse embryo specimens without digestion kinetics matching**. (**a**–**d**) Representative photographs of hydrogel-embedded E15.5 mouse embryo slices after proteinase K digestion for 12 h. **a**, 2-mm-thick slice; **b**–**d**, 500-μm-thick slices. (**e**) The embryo shown in **a** was separated from the surrounding gel during the further digestion process due to a large expansion factor mismatch (photograph not shown). (**f**–**h**) Photographs of the samples corresponding to **b**, **c**, and **d** after 84 h of proteinase K digestion. Magenta arrowheads, regions of soft tissues with relatively large swellings; Black arrowheads, flipped edges of samples due to heterogeneous expansion; Magenta dotted lines, cracks caused by different expansion ratios during the digestion process. Scale bars: 5 mm.

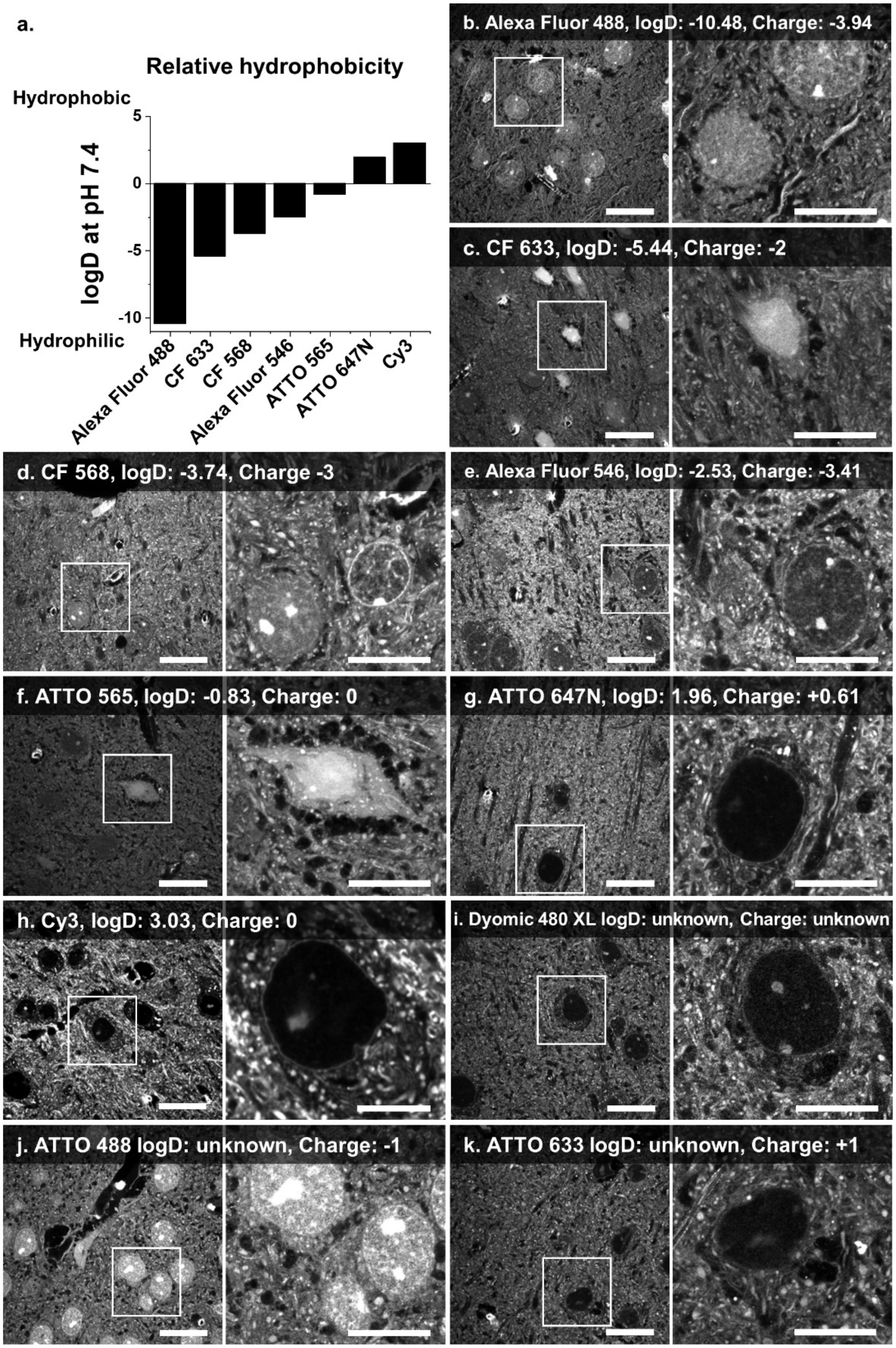
a.
Relative hydrophobicity
Hydrophobic
Hydrophilic
logD at pH 7.4
5
0
-5
-10
Alexa Fluor 488
CF 633
CF 568
Alexa Fluor 546
ATTO 565
ATTO 647N
Cy3
b. Alexa Fluor 488, logD: -10.48, Charge: -3.94
c. CF 633, logD: -5.44, Charge: -2
d. CF 568, logD: -3.74, Charge -3
e. Alexa Fluor 546, logD: -2.53, Charge: -3.41
f. ATTO 565, logD: -0.83, Charge: 0
g. ATTO 647N, logD: 1.96, Charge: +0.61
h. Cy3, logD: 3.03, Charge: 0
i. Dyomic 480 XL logD: unknown, Charge: unknown
j. ATTO 488 logD: unknown, Charge: -1
k. ATTO 633 logD: unknown, Charge: +1

**Supplementary Fig. 3**: **Fluorophore NHS-ester screening in mouse brain**. (**a**) Relative hydrophobicity of fluorescent dyes measured from the diffusivity of the fluorophores. Fluorophores with a higher logD are more hydrophobic than those with a lower logD. Reproduced from Zanetti-Domingues *et al.*[1] LogD was defined as: $logD_{octanol/water} = log([solute]_{octanol}) - log([solute]_{ionized\ water} + [solute]_{neutral\ water})$, which is the ratio of the fluorophore concentrations in water and octanol (non-polar solvent). The larger the logD value of the fluorophore is, the more hydrophobic it is. (**b**–**h**) Fluorophores with known hydrophobicities and charge states. (**b**) Alexa Fluor 488 NHS ester; (**c**) CF 633 NHS ester; (**d**) CF 568 NHS ester; (**e**) Alexa Fluor 546 NHS ester; (**f**) ATTO 565 NHS ester; (**g**) ATTO 647N NHS ester; (**h**) Cy3 NHS ester. (**i**–**k**) Fluorophores with unknown hydrophobicities and charge states. (**i**) Dyomic 480 XL NHS ester; (**j**) ATTO 488 NHS ester; (**k**) ATTO 633 NHS ester. Right images in (**b**–**k**) are magnified views of the boxed region in corresponding left images. Scale bars: (**b**–**k**) 20 µm, and (magnified views in **b**–**k**) 5 µm. All length scales are presented in pre-expansion dimensions.

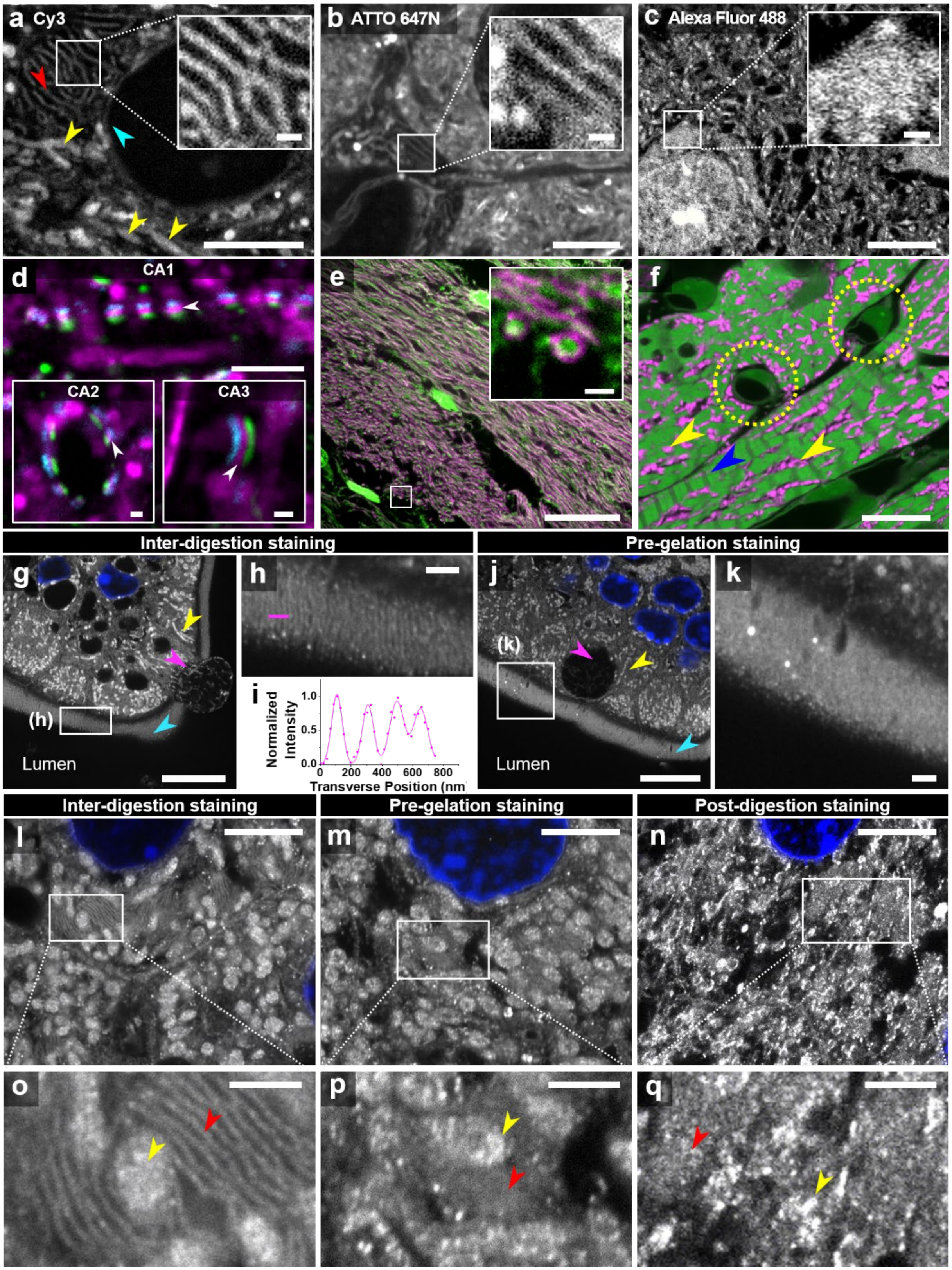

a Cy3
b ATTO 647N
c Alexa Fluor 488
d
CA1
CA2
CA3
e
f
Inter-digestion staining
Pre-gelation staining
g
(h)
Lumen
h
i
Normalized Intensity
1.0
0.5
0.0
0
200
400
600
800
Transverse Position (nm
j
(k)
Lumen
k
Inter-digestion staining
Pre-gelation staining
Post-digestion staining
l
m
n
o
p
q

**Supplementary Fig. 4**: **Analysis of the fluorophore NHS-ester staining pattern in expanded mouse organs**. (**a**–**c**) Confocal microscopy images of fluorophore NHS-ester-labeled mouse brain slice after expansion. Inset images in **a**–**c** display a magnified view of the boxed regions. Red arrowhead in **a** indicates endoplasmic reticulum. Yellow arrowheads in **a** indicate mitochondria. Cyan arrowhead in **a** indicates nuclear membrane. **a**, Cy3 NHS ester; **b**, ATTO 647N NHS ester; **c**, Alexa Fluor 488 NHS ester. (**d**) Confocal microscopy image of mouse brain CA1, CA2, and CA3 in the hippocampus after expansion. Green, Homer1; cyan, Bassoon; magenta, Cy3 NHS ester. White arrowheads in **d** possibly indicate synaptic vesicles at the pre-synaptic end. (**e**) Confocal microscopy image of the mouse brain corpus callosum after expansion. Green, Alexa Fluor 488 NHS ester; magenta, ATTO 647N NHS ester. The inset image displays a magnified view of the boxed region. (**f**) Confocal microscopy image of the expanded mouse heart slice labeled with Alexa Fluor 488 NHS ester and ATTO 647N NHS ester. Green, Alexa Fluor 488 NHS ester; magenta, ATTO 647N NHS ester. Blue arrowhead indicates cardiac muscle cells. Yellow dotted circles indicate capillaries. Yellow arrowheads indicate mitochondria. (**g**–**k**) ExM imaging of the mouse small intestine slice after expansion. Gray, ATTO 647N NHS ester; blue, DAPI. Magenta arrowheads indicate goblet cells. Yellow arrowheads indicate mitochondria. Cyan arrowheads indicate brush border. **g**–**i**, inter-digestion staining; **j**–**k**, pre-gelation staining. (**h**) Magnified view of the boxed region in **g**. (**i**) Multiple-peak Gaussian-fitted line profile of microvilli along the magenta line in **h**. (**k**) Magnified view of the boxed region in **j**. (**l**–**q**) Confocal microscopy images of the mouse liver slice after expansion. Gray, ATTO 647N NHS ester; blue, DAPI. **l**, **o**, inter-digestion staining; **m**, **p**, pre-gelation staining; **n**, **q**, post-digestion staining. (**o**) Magnified view of the boxed region in **l**. (**p**) Magnified view of the boxed region in **m**. (**q**) Magnified view of the boxed region in **n**. Yellow arrowheads in **o**, **p**, **q** indicate mitochondria. Red arrowheads in **o**, **p**, **q** indicate Golgi apparatus. Scale bars: (**a**–**c**) 5 µm, insets in (**a**–**c**) 500 nm, (**d**) 5 µm, insets in (**d**) 250 nm, (**e**) 20 µm, inset in (**e**) 500 nm, (**f**) 10 µm, (**g**) 10 µm, (**h**) 1 µm, (**j**) 10 µm, (**k**) 1 µm, (**l**–**n**) 5 µm, and (**o**–**q**) 1 µm. All length scales are presented in pre-expansion dimensions.

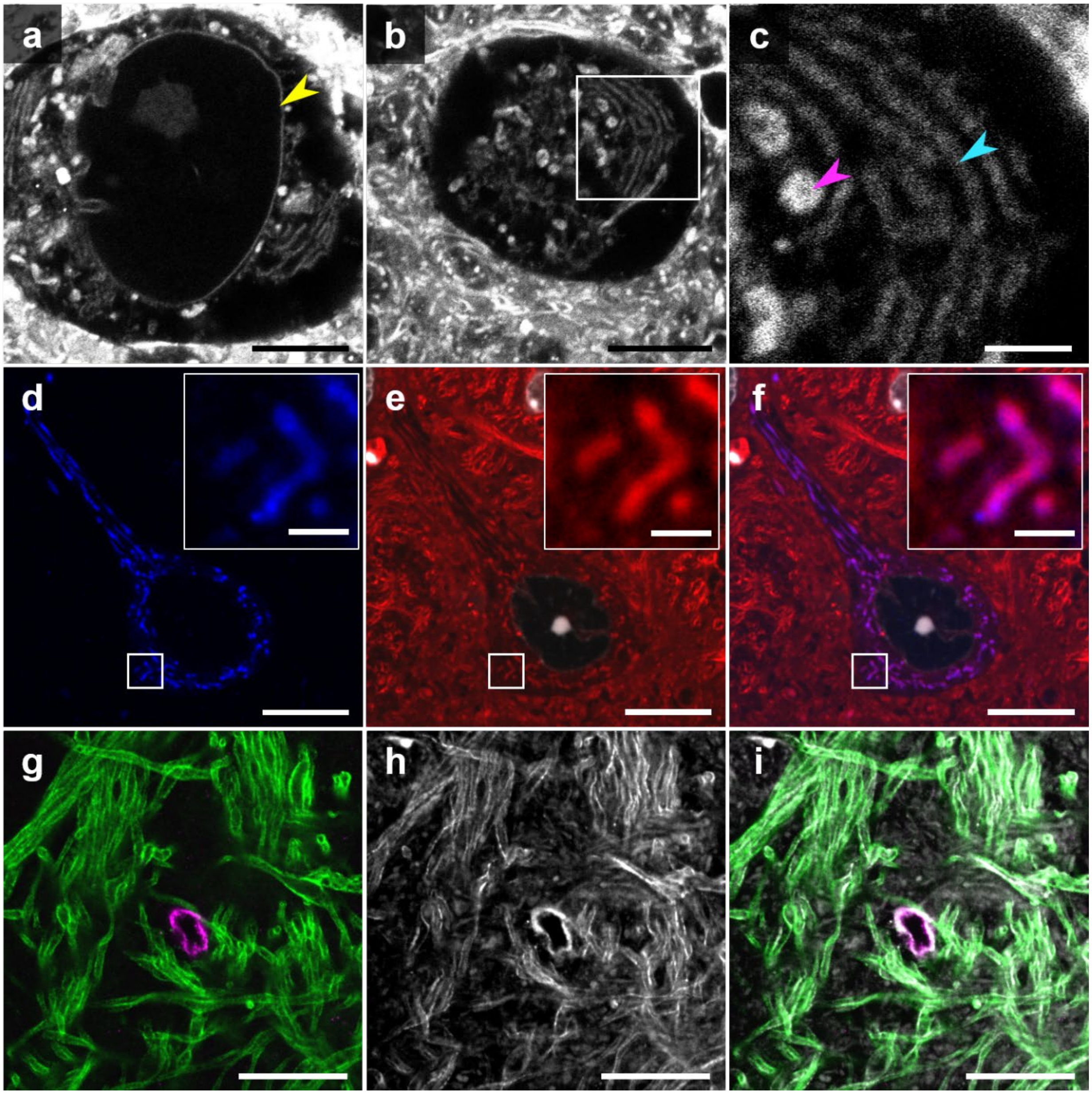


**Supplementary Fig. 5**: **ExM imaging of mouse brain slices labeled with antibodies and fluorophore NHS ester**. (**a**–**b**) Confocal microscopy images of a soma of ATTO 647N NHS-ester-labeled mouse brain DG after expansion. Images in **a** and **b** were acquired at different z-heights. Yellow arrowhead in **a** indicates nuclear membrane. (**c**) Magnified view of the boxed region in **b**. Magenta arrowhead indicates mitochondria. Cyan arrowhead indicates Golgi apparatus. (**d**–**f**) Confocal microscopy images of mouse brain SNr in basal ganglia after expansion. Blue, mScarlet-mitochondria labeled with a rabbit antibody against RFP; red, ATTO 647N NHS ester. (**g**–**i**) Confocal microscopy images of mouse brain cortex after expansion, Green, MBP; magenta, glucose transporter 1; gray, ATTO 647N NHS ester. Scale bars: (**a**–**b**) 5 μm, (**c**) 1 μm, (**d**–**i**) 10 μm, and inset in (**d**–**f**) 5 μm. All length scales are presented in pre-expansion dimensions.

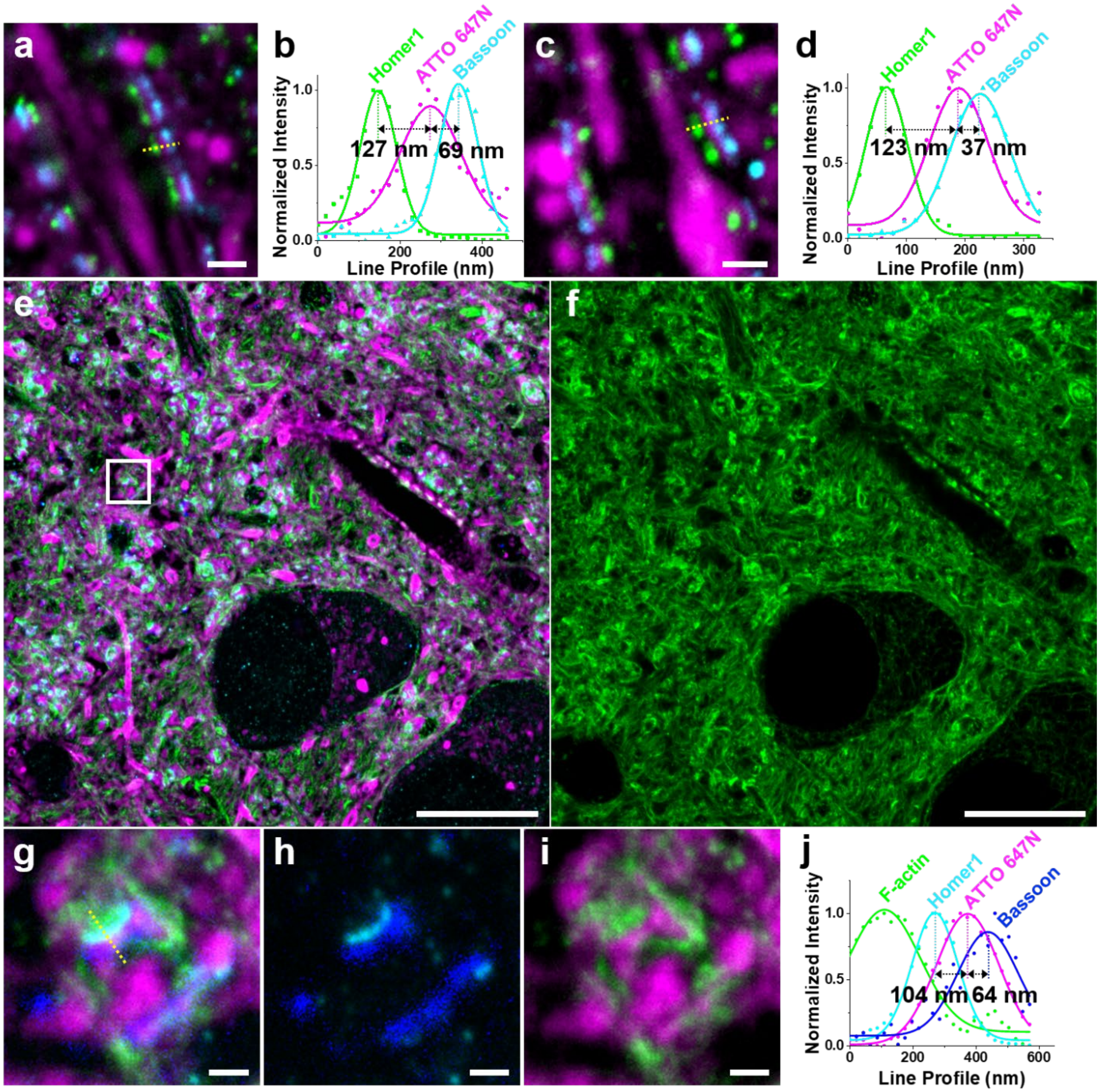
a
b
Homer1
ATTO 647N
Bassoon
Normalized Intensity
127 nm
69 nm
Line Profile (nm)
c
d
Homer1
ATTO 647N
Bassoon
Normalized Intensity
123 nm
37 nm
Line Profile (nm)
e
f
g
h
i
j
F-actin
Homer1
ATTO 647N
Bassoon
Normalized Intensity
104 nm
64 nm
Line Profile (nm)

**Supplementary Fig. 6**: **ExM imaging of mouse brain synapses labeled with fluorophore NHS ester and antibodies**. (**a**, **c**) Confocal microscopy images of mouse brain hippocampus CA1 after expansion. Cyan, Bassoon; magenta, ATTO 647N NHS ester; green, Homer1. (**b**) Line profiles (dots) of Homer1, Bassoon, and ATTO 647N NHS ester along the yellow dotted line in **a**, with a superimposed fit with a Gaussian (solid lines). Peak-to-peak distance: between Homer1 and ATTO 647N NHS ester, 127 nm; between ATTO 647N NHS ester and Bassoon, 69 nm. (**d**) As in **b**, Gaussian-fitted line profiles of Homer1, Bassoon, and ATTO 647N NHS ester along the yellow dotted line in **c**. Peak-to-peak distance: between Homer1 and ATTO 647N NHS ester, 123 nm; between ATTO 647N NHS ester and Bassoon, 37 nm. (**e**) Confocal microscopy image of mouse brain hippocampus DG after expansion. Blue, Bassoon; magenta, ATTO 647N NHS ester; cyan, Homer1; green, actin filaments. (**f**) Same image as in **e**, but only displaying actin filaments. (**g**) Magnified view of the boxed region in **e**. (**h**) Same image as in **g**, but only displaying Homer1 and Bassoon. (**i**) Same image as in **g**, but only displaying actin filaments and ATTO 647N NHS ester. (**j**) Line profiles (dots) of actin filaments, Homer1, Bassoon, and ATTO 647N NHS ester along the yellow dotted line in **g,** with a superimposed fit with a Gaussian (solid lines). Peak-to-peak distance: between Homer1 and ATTO 647N NHS ester, 104 nm; between ATTO 647N NHS ester and Bassoon, 64 nm. Scale bars: (**a**) 500 nm, (**c**) 500 nm, (**e–f**) 10 μm, and (**g–i**) 500 nm. All length scales are presented in pre-expansion dimensions.

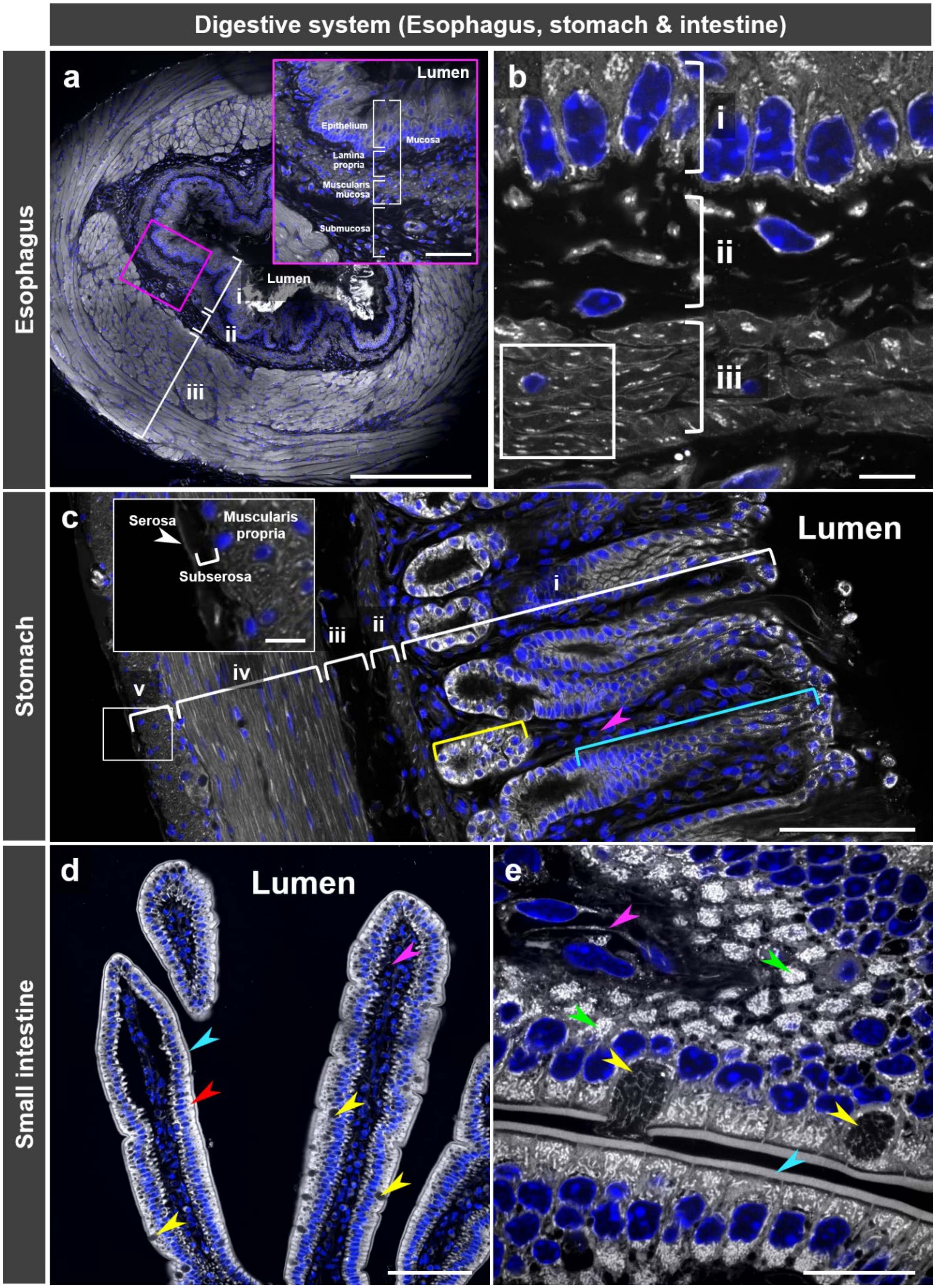
Digestive system (Esophagus, stomach & intestine)
Esophagus
Stomach
Small intestine
a
Lumen
Epithelium
Lamina propria
Muscularis mucosa
Submucosa
Mucosa
Lumen
i
ii
iii
b
i
ii
iii
c
Serosa
Muscularis propria
Subserosa
Lumen
i
ii
iii
iv
v
d
Lumen
e

**Supplementary Fig. 7**: **Mouse digestive system (Esophagus, stomach and small intestine) stained and expanded through the inter-digestion protocol**. (**a**) Confocal microscopy image of a mouse esophagus slice after expansion. Gray, Cy3 NHS ester; blue, DAPI. i, mucosa; ii, submucosa; iii, muscularis externa. Inset: magnified view of the boxed region. (**b)** Confocal microscopy images of a mouse esophagus slice after expansion. Gray, Cy3 NHS ester; blue, DAPI. i, epithelium; ii, lamina propria; iii, muscularis mucosa. (**c**) Confocal microscopy image of a mouse stomach slice after expansion. Gray, Cy3 NHS ester; blue, DAPI. i, mucosa; ii, muscularis mucosa; iii, submucosa; iv, muscularis propria (inner circular layer); v, muscularis propria (outer longitudinal layer). Cyan bracket indicates gastric pit. Magenta arrowhead indicates lamina propria. Yellow bracket indicates gastric gland. Inset shows a magnified view of the boxed region. (**d**) Confocal microscopy image of a mouse small intestine slice after expansion. Gray, ATTO 647N NHS ester; blue, DAPI. Magenta arrowhead indicates lamina propria. Yellow arrowheads indicate goblet cells. Red arrowhead indicates enterocyte. Cyan arrowhead indicates brush border. (**e**) Confocal microscopy image of a mouse small intestine slice after expansion. Gray, ATTO 647N NHS ester; blue, DAPI. Yellow arrowheads indicate goblet cells. Magenta arrowhead indicates lamina propria. Green arrowheads indicate mitochondria. Cyan arrowhead indicates brush border. Scale bar: (**a**) 200 µm, inset in (**a**) 50 µm, (**b**) 5 µm, (**c**) 100 µm, inset in (**c**) 30 µm, (**d**) 100 µm, and (**e**) 20 µm. All length scales are presented in pre-expansion dimensions.

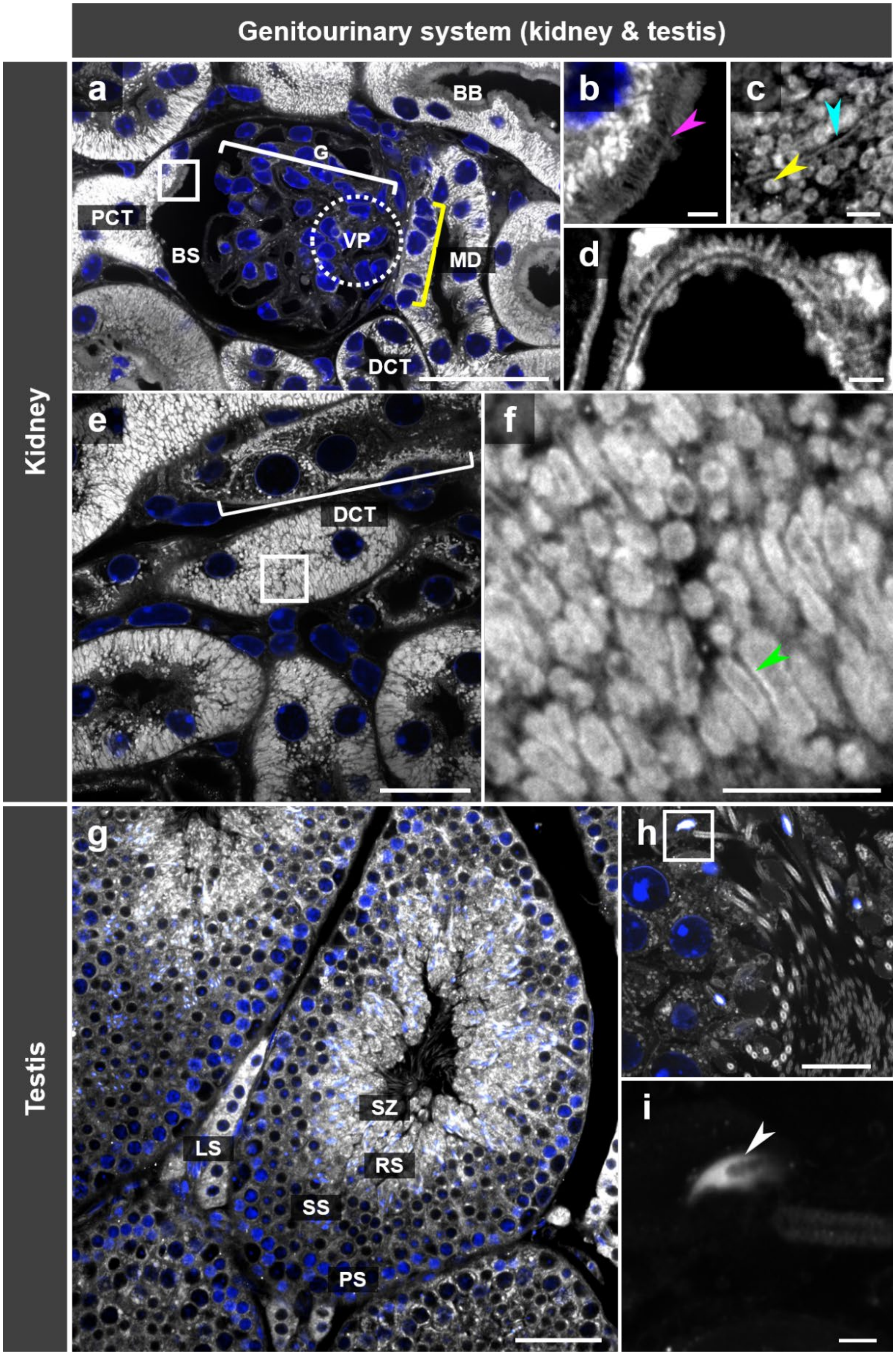

Genitourinary system (kidney & testis)
Kidney
Testis
a
BB
G
PCT
BS
VP
MD
DCT
b
c
d
e
DCT
f
g
SZ
LS
RS
SS
PS
h
i

**Supplementary Fig. 8**: **Mouse genitourinary system (kidney and testis) stained and expanded through the inter-digestion protocol**. (**a**–**f**) Confocal microscopy images of mouse kidney slices after expansion. Gray, ATTO 647N NHS ester; blue, DAPI. G, glomerulus; BB, brush border; BS, Bowman's space; PCT, proximal convoluted tubule; MD, macula densa; DCT, distal convoluted tubule; VP, vascular pole. **a**, **c**, **d** and **e** were taken from different kidney specimens. (**b**) Magnified view of the boxed region in **a**. Magenta arrowhead indicates brush border. (**c**) Mitochondria (yellow arrowhead) and basal striation (cyan arrowhead). (**d**) Foot process. (**f**) Magnified view of the boxed region in **e**. Green arrowhead indicates the outer membrane of a mitochondrion. (**g**–**i**) Confocal microscopy images of mouse testis slice after expansion. Gray, ATTO 647N NHS ester; blue, DAPI. SZ, spermatozoon; RS, round spermatids; SS, secondary spermatocytes; PS, primary spermatocytes; LS, leydig cells. **g** and **h** were taken with different objective lenses from different specimens. (**i**) Magnified view of the boxed region in **h**. White arrowhead indicates sperm head membrane. Scale bars: (**a**) 40 μm, (**b**–**c**) 2 μm, (**d**) 500 nm, (**e**) 20 μm, (**f**) 5 μm, (**g**) 50 μm, (**h**) 10 μm, and (**i**) 1 μm. All length scales are presented in pre-expansion dimensions.

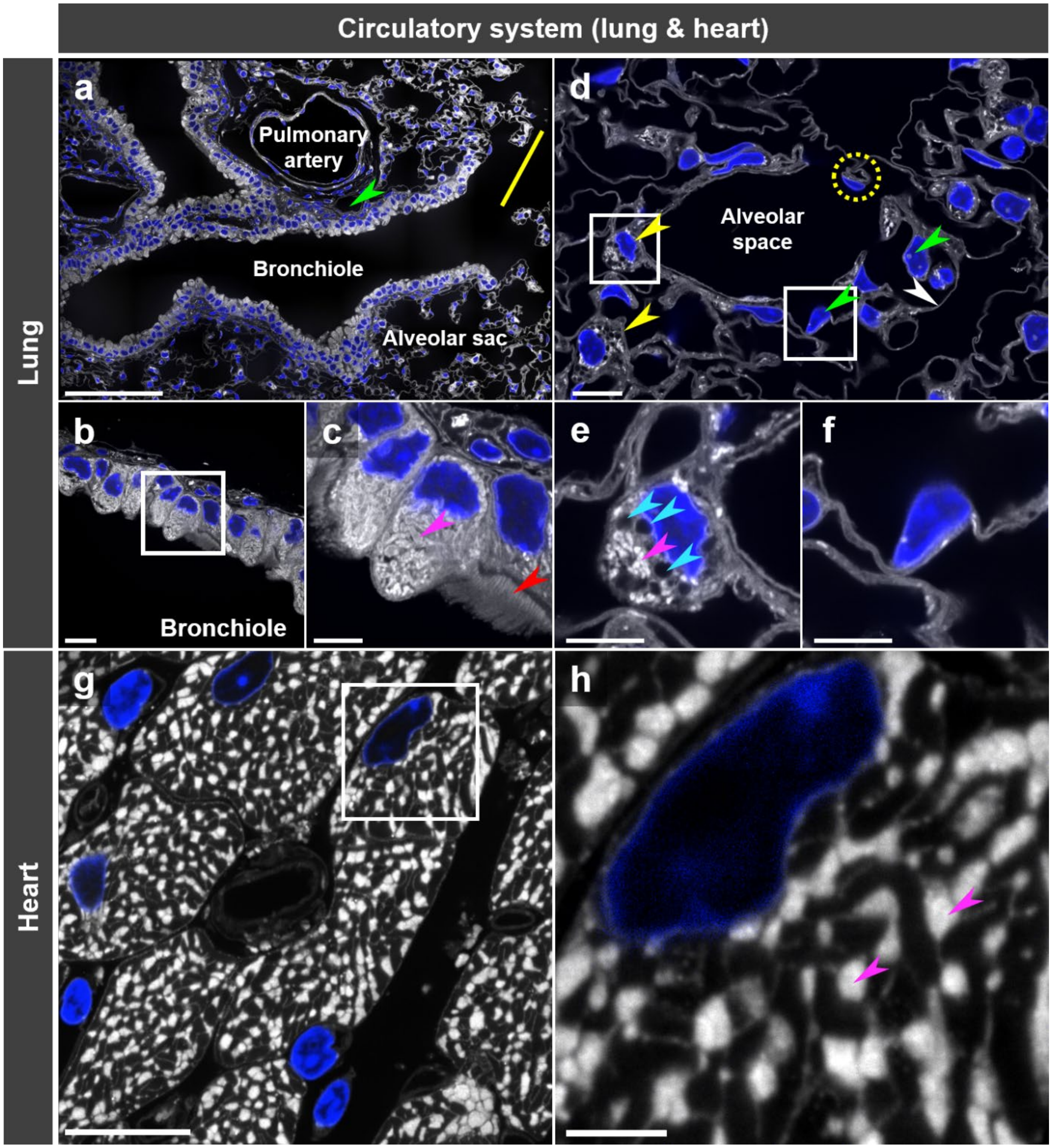
Circulatory system (lung & heart)
Lung
Heart
a
Pulmonary artery
Bronchiole
Alveolar sac
d
Alveolar space
b
Bronchiole
c
e
f
g
h

**Supplementary Fig. 9**: **Mouse circulatory system (lung and heart) stained and expanded through the inter-digestion protocol**. (**a**–**f**) Confocal microscopy images of a mouse lung slice after expansion. Gray, ATTO 647N NHS ester; blue, DAPI. Yellow line indicates alveolar ducts. Green arrowhead indicates blood vessel. (**a**) Lung bronchiole. (**b**) Clara cells adjacent to lung bronchiole. (**c**) Magnified view of the boxed region in **b**. Magenta arrowhead indicates mitochondria. Red arrowhead indicates cilia. (**d**) Confocal microscopy images of the alveolus region in a mouse lung slice after expansion. Gray, ATTO 647N NHS ester; blue, DAPI. Yellow arrowheads indicate alveolar cells (type II). Green arrowheads indicate alveolar cells (type I). White arrowhead indicates cytoplasmic extension of an alveolar cell (type I). Yellow dotted circle indicates an endothelial cell of alveolar capillary. (**e**) Magnified view of the left boxed region in **d**, displaying an alveolar cell (type II). Magenta arrowhead indicates mitochondria. Cyan arrowheads indicate secretory granules (lamellar bodies). (**f**) Magnified view of the right boxed region in **d**, displaying an alveolar cell (type I). (**g**) Confocal microscopy images of mouse heart slice after expansion. Gray, ATTO 647N NHS ester; blue, DAPI. (**h**) Magnified view of the boxed region in **g**. Magenta arrowheads indicate mitochondria. Scale bars: (**a**) 100 μm, (**b**) 10 μm, (**c**) 5 μm, (**d**) 10 μm, (**e**–**f**) 5 μm, (**g**) 10 μm, and (**h**) 2 μm. All length scales are presented in pre-expansion dimensions.

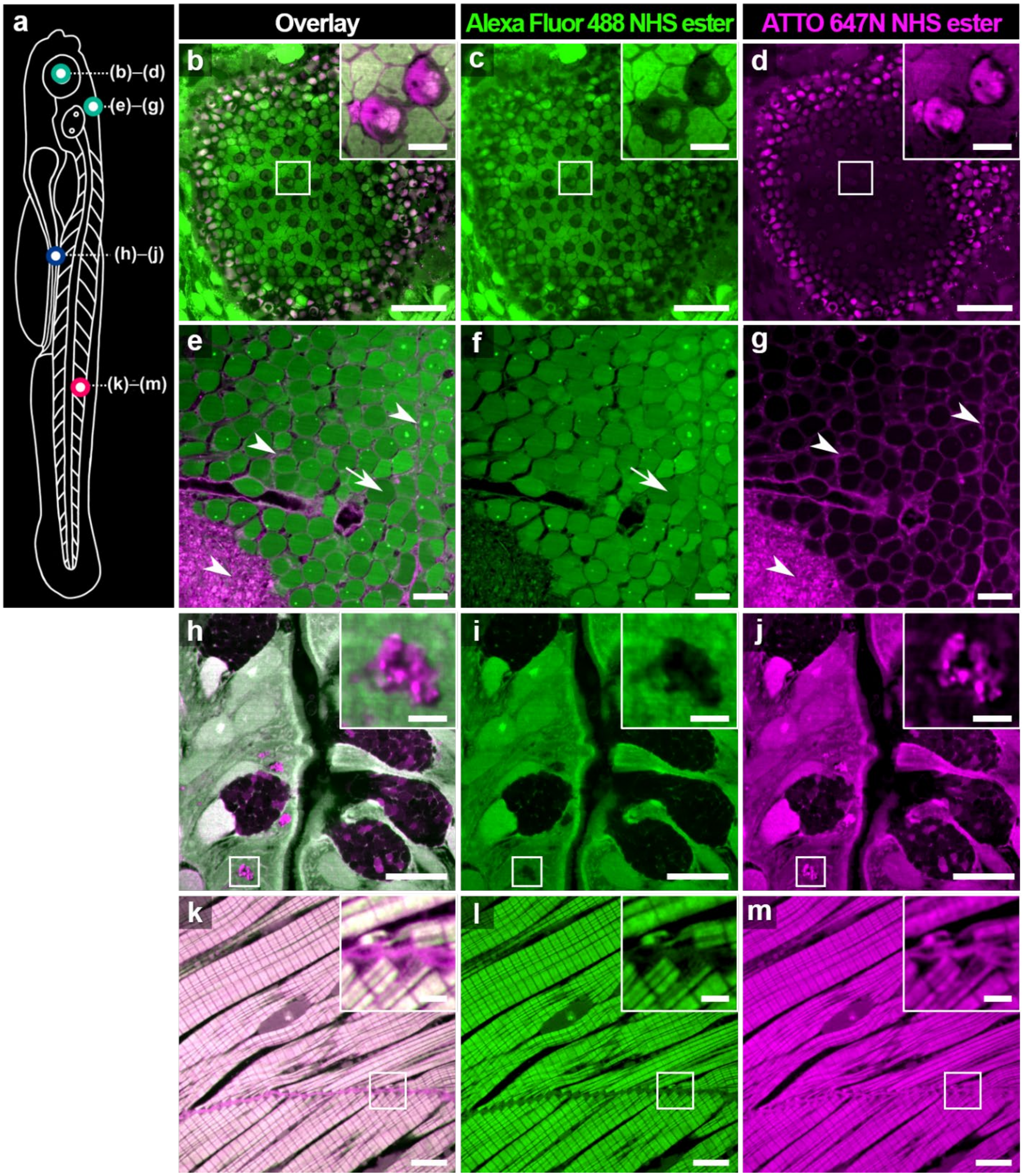
a
(b)–(d)
(e)–(g)
(h)–(j)
(k)–(m)
Overlay
Alexa Fluor 488 NHS ester
ATTO 647N NHS ester
b
c
d
e
f
g
h
i
j
k
l
m

**Supplementary Fig. 10**: **Different staining patterns of Alexa Fluor 488 NHS ester and ATTO 647N NHS ester in larval zebrafish**. (**a**) Schematic diagram depicting the locations where the images shown in **b**–**m** were acquired. (**b**–**m**) Confocal microscopy images of expanded zebrafish at 6 dpf, co-stained with Alexa Fluor 488 NHS ester (green) and ATTO 647N NHS ester (magenta), together with their respective individual channels, including (**b**–**d**) a mosaic arrangement of cone photoreceptors, (**e**–**g**) cell bodies and blood vessels in the brain. Arrowheads in **e**–**g**, cytoplasm; arrows in **e**–**g**, nuclei. (**h**–**j**) goblet cells and secreted mucins in the mid-intestine, and (**k**–**m**) skeletal muscle fibers. Scale bars: (**b**–**d**) 10 µm, inset of (**b**–**d**) 2 µm, (**e**–**g**) 5 µm, (**h**–**j**) 10 µm, inset of (**h**–**j**) 2 µm, (**k**–**m**) 10 µm, and inset of (**k**–**m**) 2 µm. All length scales are presented in pre-expansion dimensions.

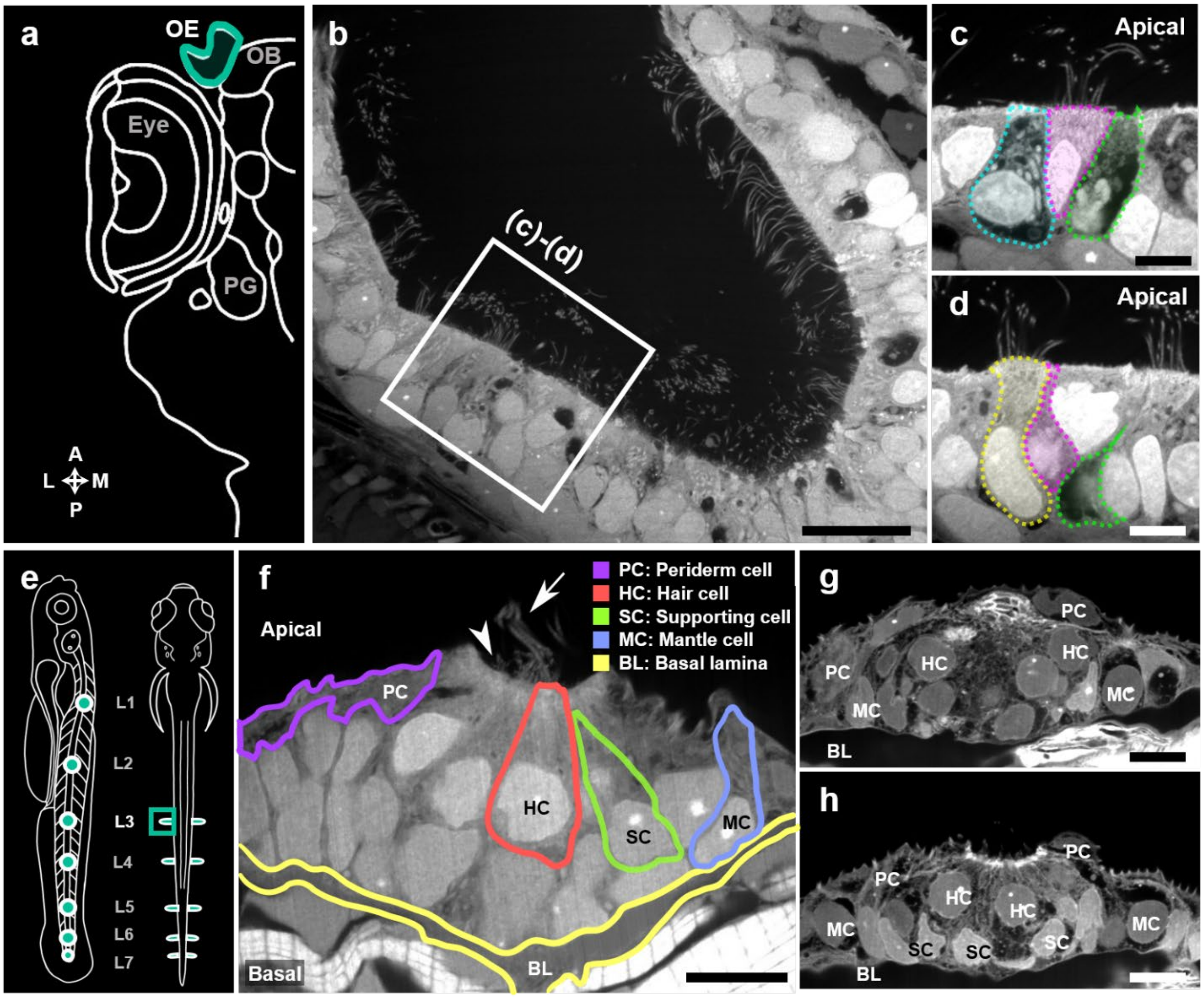


**Supplementary Fig. 11**: **Whole-body ExM imaging of the zebrafish olfactory sensory epithelium and trunk neuromast**. (**a**) Schematic diagram showing the position of olfactory epithelium (OE) in a horizontal section through the zebrafish head. OB, olfactory bulb; PG, preglomerular complex. (**b**) Confocal microscopy image of olfactory sensory epithelium of an expanded zebrafish larva at 6 dpf. (**c**–**d**) Magnified views of the boxed region in **b** with different *z*-planes. Each of the cells that compose the sensory epithelium is colored in a different color. (**e**) Schematic drawing of the neuromast distribution and hair cell orientation in the posterior lateral line viewed from lateral (left) and dorsal (right). (**f**) Confocal microscopy image of the L3 trunk neuromast of an expanded zebrafish larva at 6 dpf, showing the general organization of the neuromast and basal epithelial lamina. A stereocilia bundle (arrowhead) from one of the central hair cells and a kinocilium (arrow) from another protrude into the apical side. (**g**) Confocal microscopy images of the L3 trunk neuromast of an expanded zebrafish larva at 6 dpf. (**h**) Confocal microscopy images of the L3 trunk neuromast taken from the same region as **g**, but at a different *z*-plane. Labels: (**b**–**d**, **f**) Alexa Fluor 488 NHS ester; (**g**–**h**) ATTO 647N NHS ester. Scale bars: (**b**) 5 µm, (**c**–**d**) 2 µm, and (**f**–**h**) 5 µm. All length scales are presented in pre-expansion dimensions.

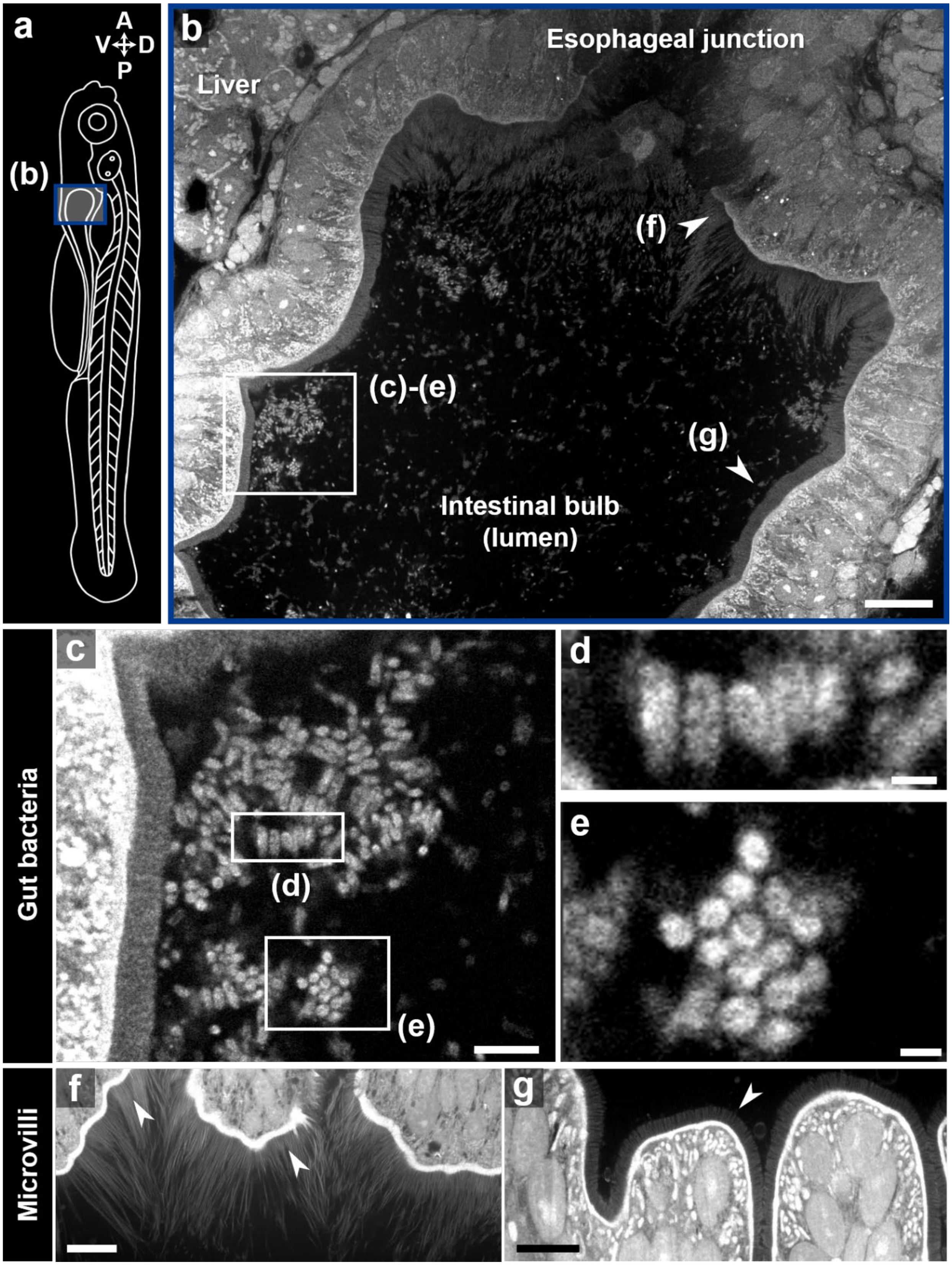
a
A
V
D
P
(b)
b
Esophageal junction
Liver
(f)
(c)-(e)
(g)
Intestinal bulb
(lumen)
Gut bacteria
c
(d)
(e)
d
e
Microvilli
f
g

**Supplementary Fig. 12**: **Whole-body ExM imaging of the zebrafish intestinal bulb**. (**a**) Schematic diagram showing the locations where the images in **b**–**e** were acquired. (**b**) Confocal microscopy image of gut bacteria in the intestinal bulb of an expanded zebrafish at 6 dpf. Bacteria were clustered together in the luminal space, with some remaining close to the host epithelium (arrowheads). (**c**) Magnified view of the boxed region in **b**. (**d**–**e**) Magnified views of the boxed region in **c**, highlighting the rod shape of the bacteria. (**f**) Unusually long microvilli found in the anterior intestinal bulb region adjacent to the esophageal intestinal junction. (**g**) Microvilli with a shorter and uniform length, typical shape of the brush border. All labels: Alexa Fluor 488 NHS ester. Scale bars: (**b**) 20 µm, (**c**) 5 µm, (**d**–**e**) 1 µm, and (**f**–**g**) 10 µm. All length scales are presented in pre-expansion dimensions.

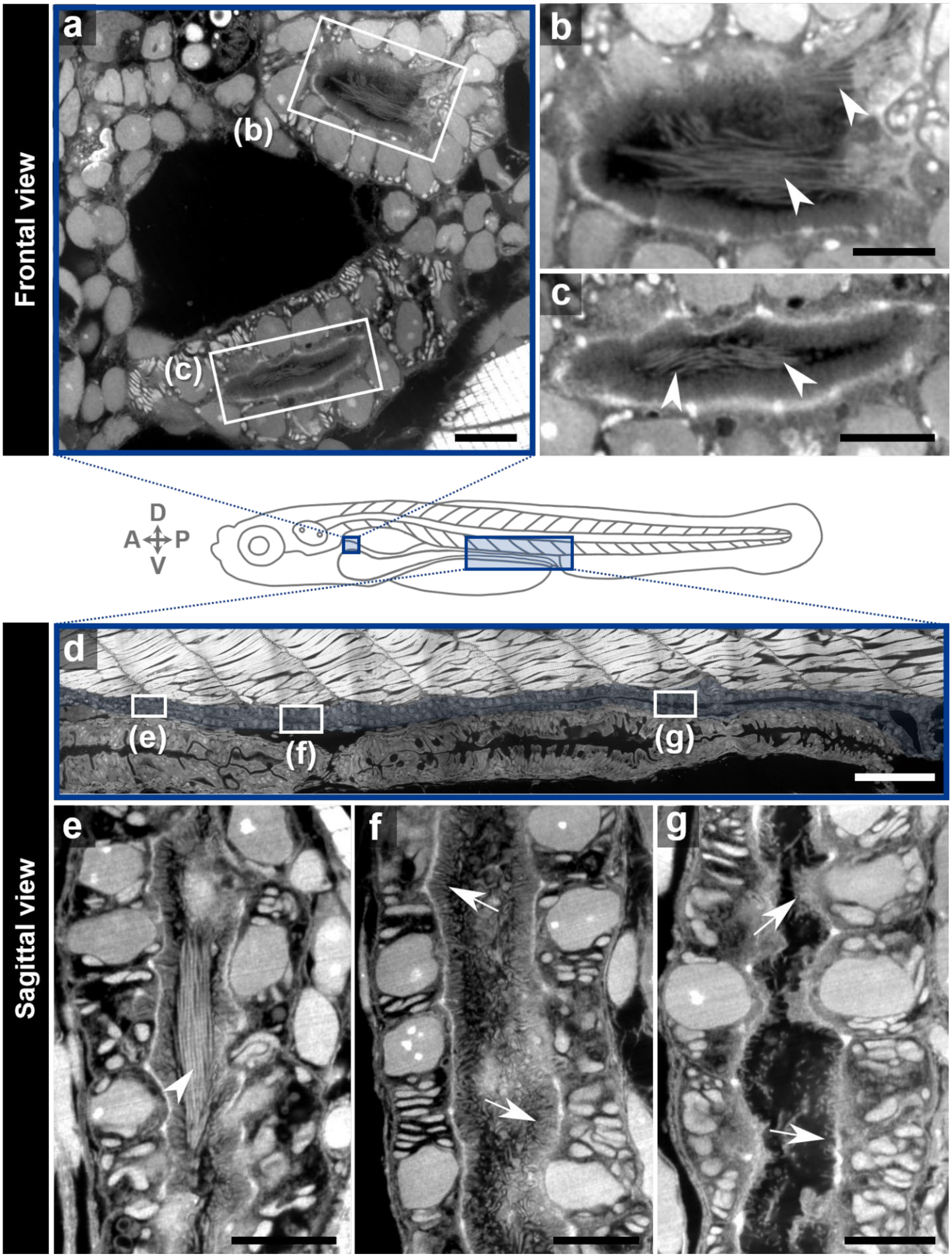
Frontal view
a
(b)
(c)
b
c
D
A
P
V
Sagittal view
d
(e)
(f)
(g)
e
f
g

**Supplementary Fig. 13**: **Whole-body ExM imaging of the zebrafish pronephros**. (**a**) Confocal microscopy image of the pronephric duct (frontal view) of an expanded zebrafish larva at 6 dpf, showing two proximal tubules. (**b–c**) Magnified views of the boxed region in **a**. Arrowheads indicate motile cilia. (**d**) Confocal microscopy image of the pronephric duct (sagittal view) of a 500-μm-long trunk region showing the general morphology of the pronephric duct (colored) located between the swimming muscles and intestine. (**e–g**) Magnified view of the boxed regions in **d**, highlighting apical surfaces of the pronephric duct. Arrowhead, a bundle of cilia; arrows, the apical surface of multi-ciliated cells. All labels: Alexa Fluor 488 NHS-ester. Scale bars: (**a**) 10 μm, (**b–c**) 5 μm, (**d**) 50 μm, and (**e–g**) 5 μm. All length scales are presented in pre-expansion dimensions.

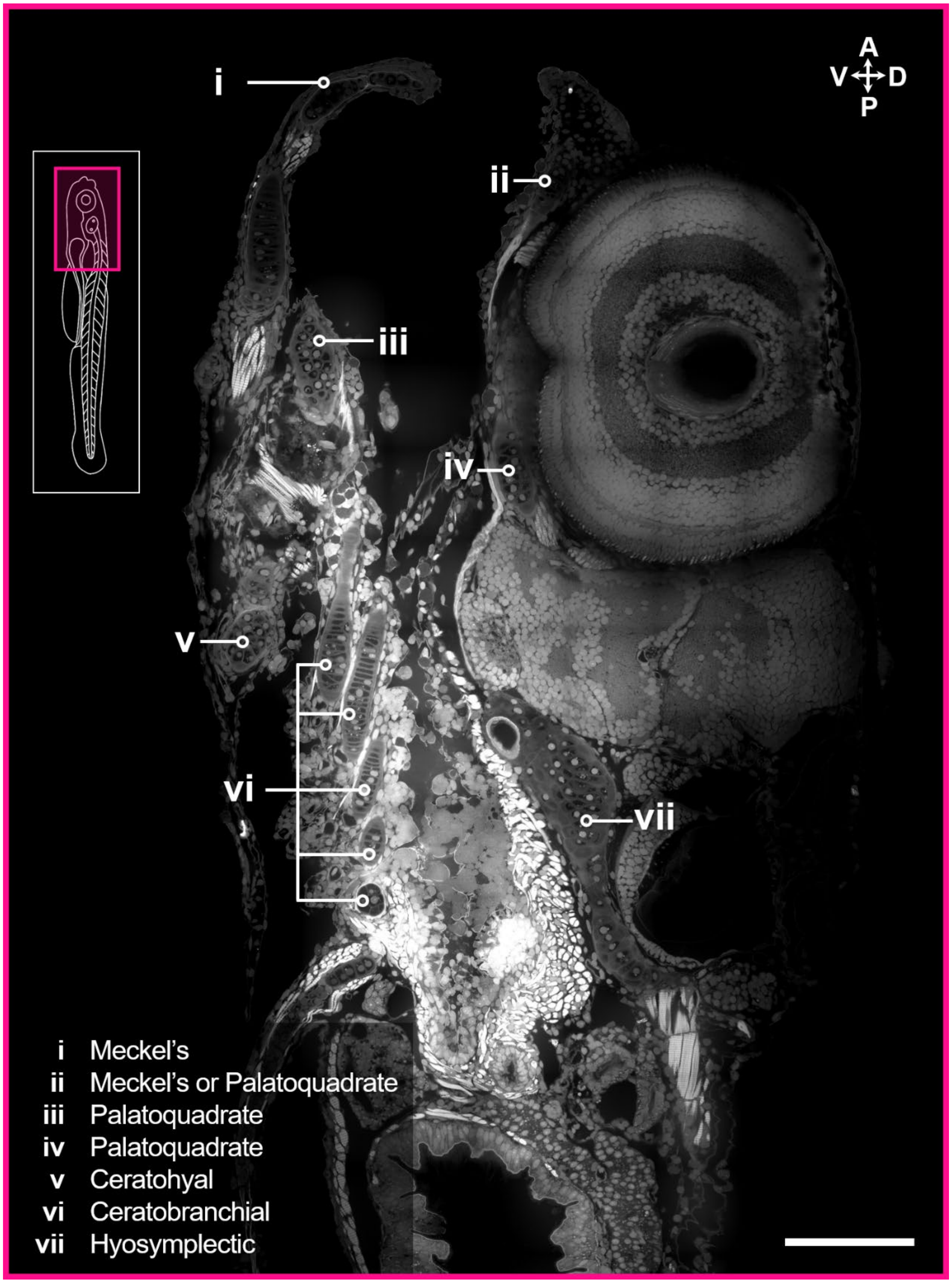


**Supplementary Fig. 14**: **Whole-body ExM imaging of the zebrafish pharyngeal arches**. Confocal microscopy image of the pharyngeal arches (oblique-frontal view) of an expanded zebrafish larva at 6 dpf. Label: ATTO 647N NHS ester. Scale bar: 100 µm. Length scale is presented in pre-expansion dimension.

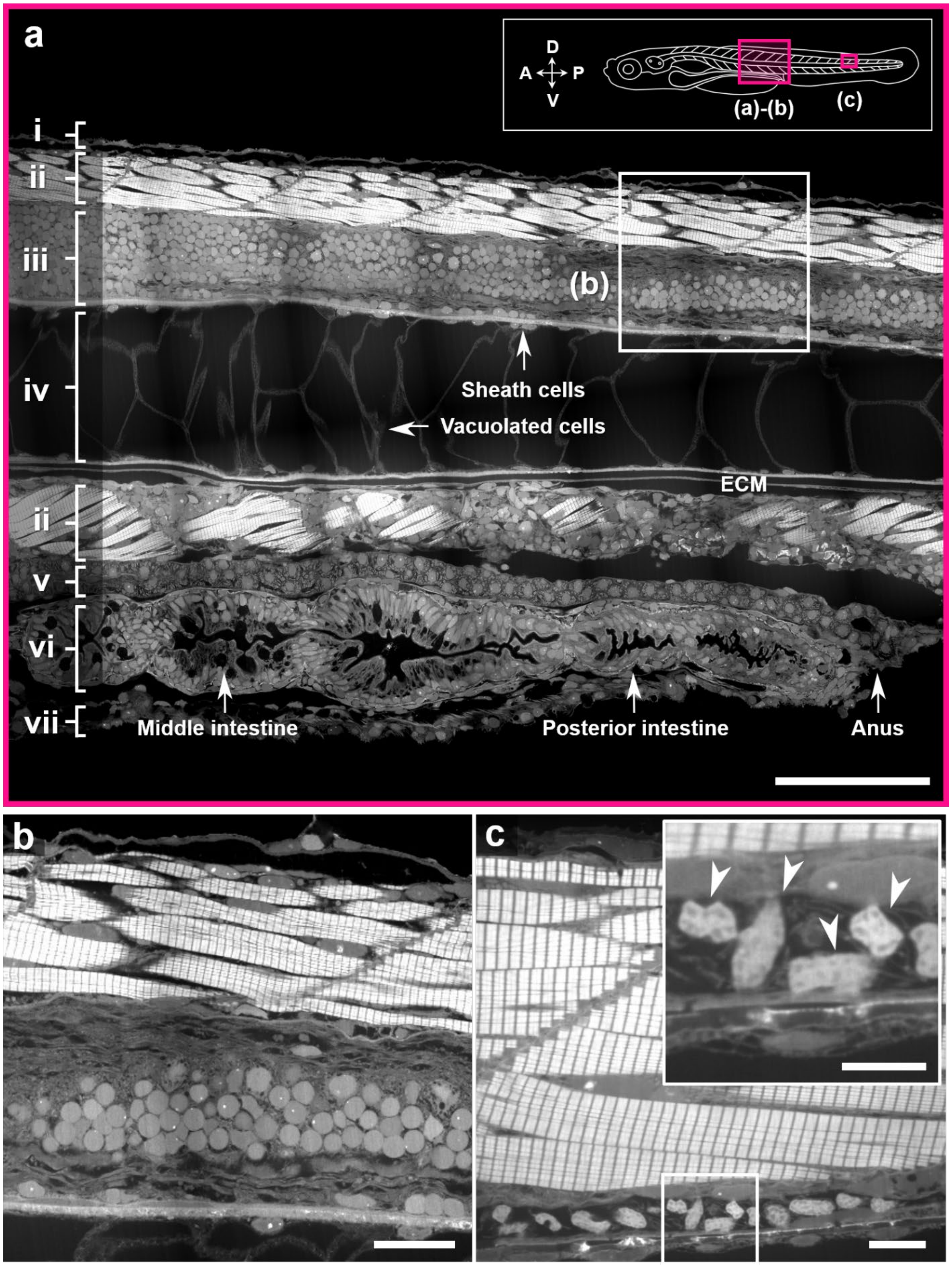
a
D
A
P
V
(a)-(b)
(c)
i
ii
iii
iv
ii
v
vi
vii
(b)
Sheath cells
Vacuolated cells
ECM
Middle intestine
Posterior intestine
Anus
b
c

**Supplementary Fig. 15**: **Whole-body ExM imaging of the zebrafish body trunk**. (**a**) Confocal microscopy image of the body trunk (sagittal view) of an expanded zebrafish larva at 6 dpf. i, dorsal fin; ii, myotomes; iii, spinal cord; iv, notochord; v, pronephric duct; vi, intestine; vii, ventral fin. (**b**) Magnified view of the boxed region in **a**, showing borders of myotome, spinal cord, and notochord. (**c**) The posterior region in the same view, showing dorsal longitudinal anastomotic vessel between myotome and neural tube. Inset shows magnified view of blood cells (arrowheads) abundant within the vessel. All labels: Alexa Fluor 488 NHS ester. Scale bars: (**a**) 50 μm, (**b**) 20 μm, (**c**) 10 μm, and inset in (**c**) 5 μm. All length scales are presented in pre-expansion dimensions.

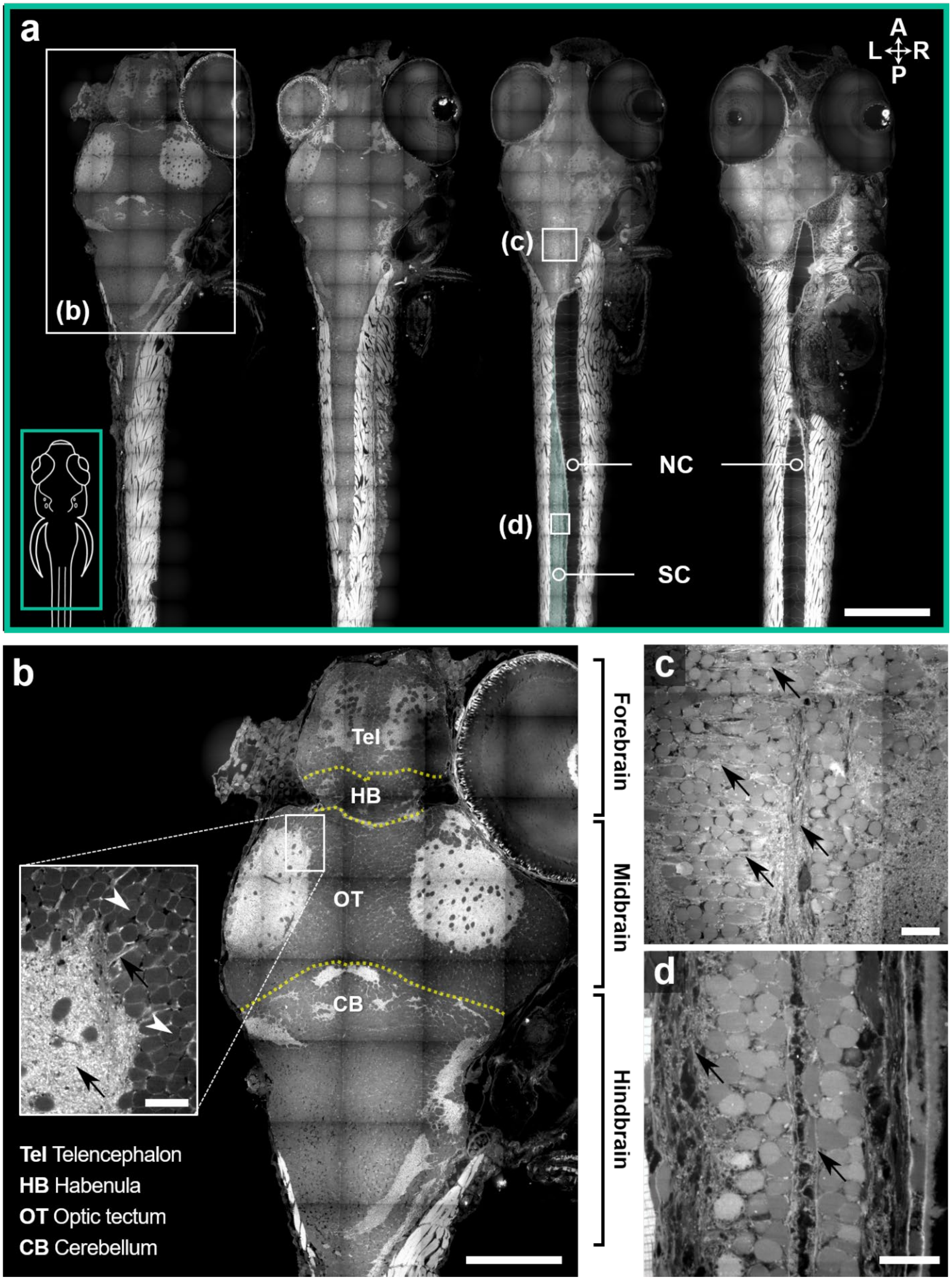
a
A
L
R
P
(b)
(c)
(d)
NC
SC
b
Tel
HB
OT
CB
Tel Telencephalon
HB Habenula
OT Optic tectum
CB Cerebellum
Forebrain
Midbrain
Hindbrain
c
d

**Supplementary Fig. 16**: **Whole-body ExM imaging of the zebrafish brain and spinal cord**. (**a**) Series of montage images taken in a dorsal view at four different *z*-heights, showing the brain and spinal cord of an expanded zebrafish larva at 6 dpf. Notochord (NC) and spinal cord (SC) are marked. (**b**) Magnified view of the boxed region in **a**, indicating well-distinguished anatomical sections of the larval brain. Tel, telencephalon; HB, habenula; OT, optic tectum; CB, cerebellum. (**c**–**d**) Magnified view of the boxed region in **a**, highlighting detailed structures of the brain and spinal cord, respectively. White arrowheads in **b**, nuclei; black arrows in **b**–**d**, cytoplasm. All labels: ATTO 647N NHS ester. Scale bars: (**a**) 200 μm, (**b**) 100 μm, (**c**) 10 μm, (**d**) 15 μm, and inset in (**b**–**d**) 10 μm. All length scales are presented in pre-expansion dimensions.

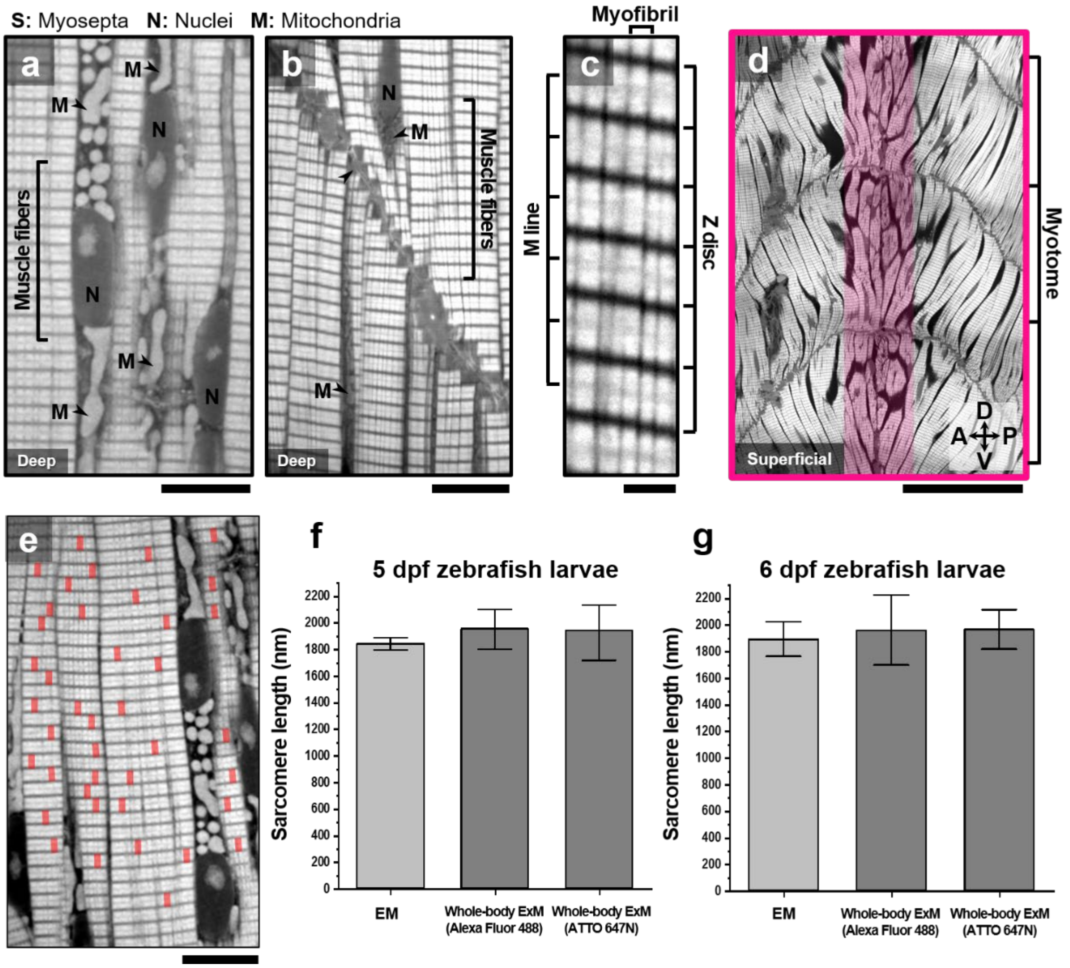


**Supplementary Fig. 17**: **Expansion homogeneity of whole-body ExM**. (**a**–**d**) High magnification view of the muscle fibers depicting a single dorsal. S, myosepta; N, nuclei; M, mitochondria. (**c**) Magnified view highlighting the Z- and M-lines in single myofibrils. Labels: (**a**), (**c**) ATTO 647N NHS ester; (**b**), (**d**), (**e**) Alexa Fluor 488 NHS ester. (**d**) Confocal microscopy image of the chevron-shaped myotomes (lateral view) of an expanded zebrafish larva at 6 dpf. The colored region indicates the horizontal boundary cells in the horizontal myoseptum. (**e**) Representative image showing how the sarcomere lengths were measured from skeletal muscle fibers. The sarcomere length was measured from a Z-disc to a Z-disc. More than 30 sarcomeres were randomly selected in skeletal muscle fibers of larval zebrafish at 5 and 6 dpf. (**f**–**g**) Statistical comparison of the sarcomere length by electron microscopy and whole-body ExM. (**f**) Length of the sarcomere of larval zebrafish at 5 dpf (mean ± standard deviation). EM, 1.84 ± 0.27 µm (n=23); whole-ExM (Alexa Fluor 488 NHS ester), 1.93 ± 0.10 µm (n=35); whole-body ExM (ATTO 647N NHS ester), 1.94 ± 0.11 µm (n=35). (**g**) Length of the sarcomere of larval zebrafish at 6 dpf. EM, 1.89 ± 0.79 µm (n=33); whole-body ExM (Alexa Fluor 488 NHS ester), 1.96 ± 0.14 µm (n=33); whole-body ExM (ATTO 647N NHS ester), 1.96 ± 0.12 µm (n=33). The average sarcomere length measured from two EM data sets in **b** and **c** was 1.87 ± 0.07 µm. The average of sarcomere length measured from four expanded larvae was 1.95 ± 0.12 µm. Scale bars: (**a**,**b**) 10 µm, (**c**) 2 µm, (**d**) 50 µm, and (**e**) 10 µm. All length scales are presented in pre-expansion dimensions.

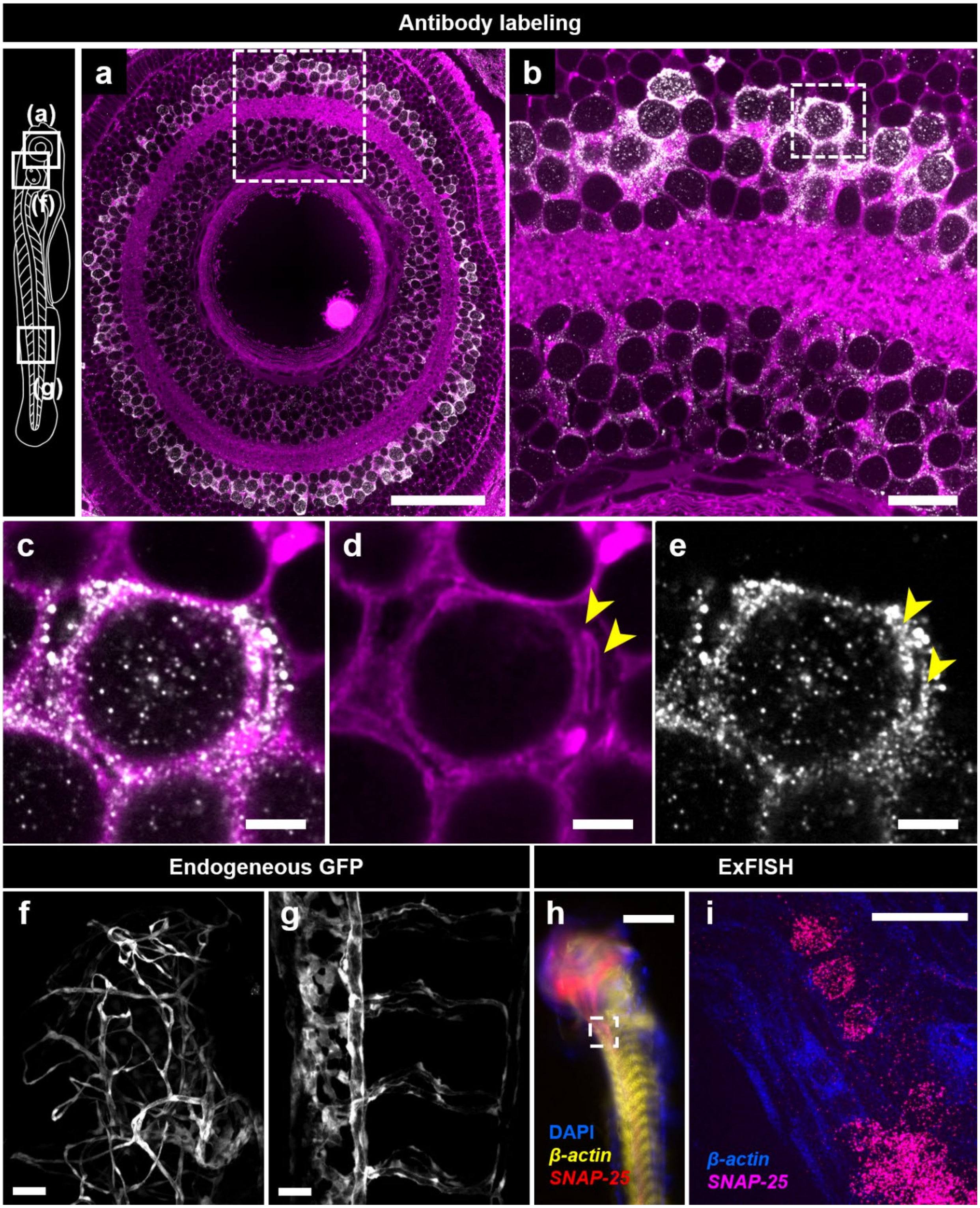
Antibody labeling
(a)
(f)
(g)
a
b
c
d
e
Endogeneous GFP
ExFISH
f
g
h
i
DAPI
β-actin
SNAP-25
β-actin
SNAP-25

**Supplementary Fig. 18**: **Whole-body ExM imaging of zebrafish labeled with antibodies and fluorophore NHS ester and ExFISH**. (**a**) Confocal microscopy images of anti-HuC/D antibody, ATTO 647N NHS-ester-labeled 3 dpf zebrafish larvae after expansion. Gray, anti HuC/D antibody; Magenta, ATTO 647N NHS ester. (**b**) Magnified view of the boxed region in **a**. (**c**–**e**) Magnified view of the boxed region in **b**. (**f**–**g**) Confocal microscopy image of an expanded 3 dpf, *Tg*(*kdrl*:*EGFP*) zebrafish larva showing the endogenous GFP signal retention after 54 h-long digestion. GFP labels vasculature. Location of **f**, **g** are annotated in the schematic image. (**h**–**i**) Confocal microscopy images of 5 dpf zebrafish larvae with ExFISH. DAPI; *β-actin*; *SNAP-25*; image of the three channels. (**i**) Magnified, maximum intensity projected view of the boxed region in **h**. Scale bars: (**a**) 50 µm, (**b**) 20 µm, (**c**–**e**) 10 µm, (**f**) 20 µm, (**g**) 10 µm, (**h**) 500 µm, and (**i**) 50 µm. All length scales are presented in pre-expansion dimensions.

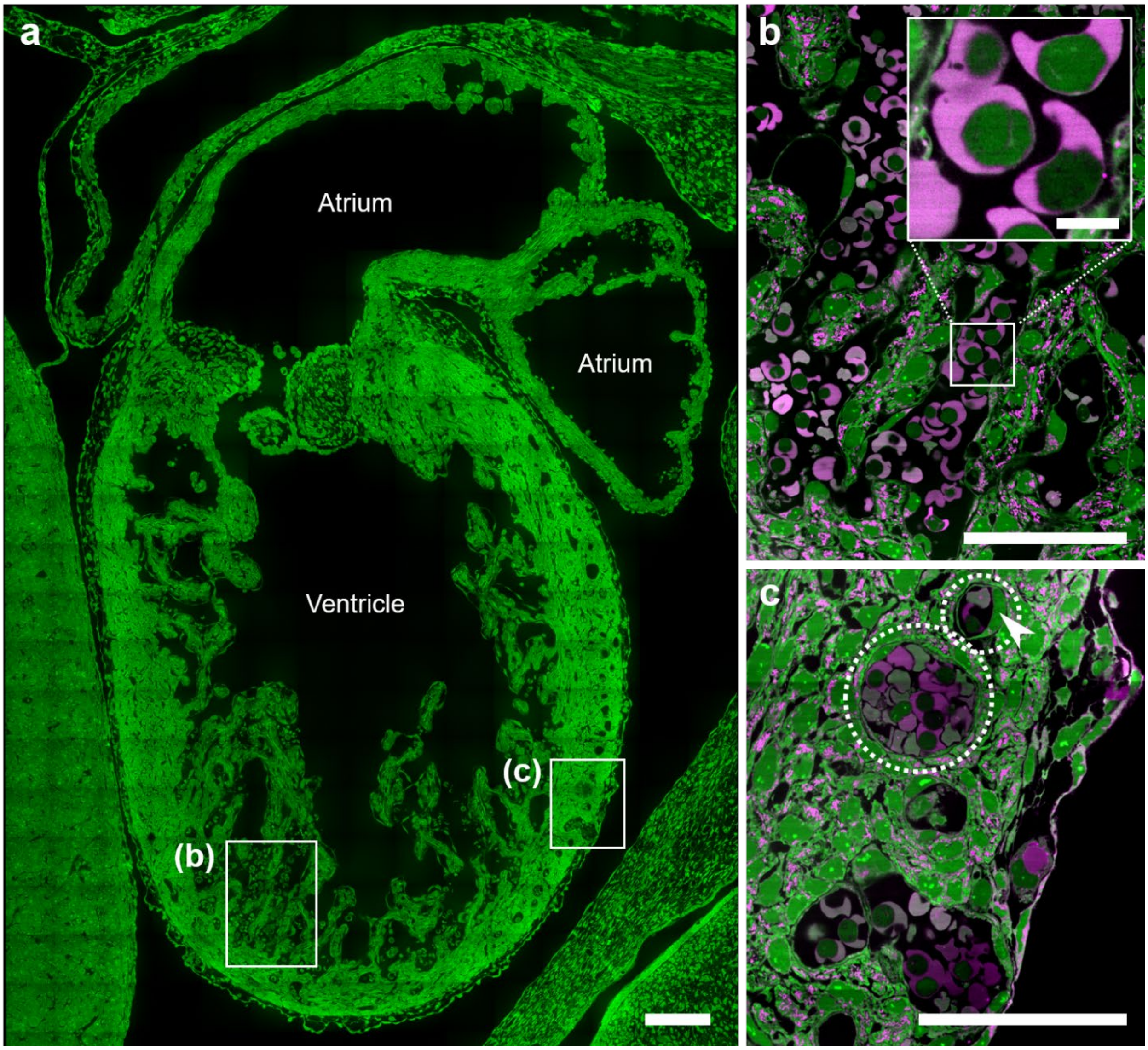


**Supplementary Fig. 19**: **Whole-body ExM imaging of the mouse embryo heart**. (**a**) Confocal microscopy image of the E13.5 mouse embryo heart after expansion. Green, Alexa Fluor 488 NHS-ester; Magenta, ATTO 647N NHS-ester. (**b**–**c**) Magnified views of the boxed regions in **a**, showing blood cells in (**b**) ventricle lumen, and (**c**) blood vessels, respectively. Arrowhead indicates an endothelial cell. Scale bars: (**a**) 100 µm, (**b**) 50 µm, inset in (**b**) 10 µm, and (**c**) 50 µm. All length scales are presented in pre-expansion dimensions.

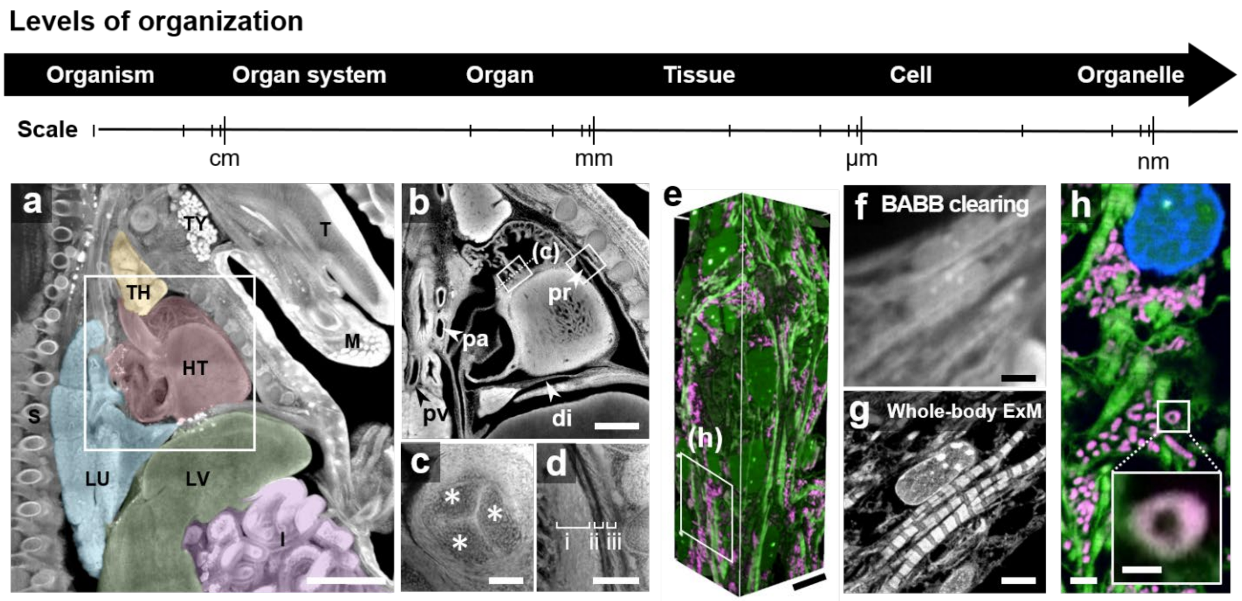


**Supplementary Fig**. **20**: **Whole-body ExM imaging of mouse embryo in different length scales**. (**a**) Confocal microscopy image of body of E15.5 mouse embryo post expansion. Major internal organs are displayed with different colors. T, tongue; M, mandible; TH, thymus; HT, heart; LU, lung; LV, liver; I, intestine. (**b**) Confocal microscopy image of heart and its neighboring tissues of E15.5 mouse embryo. pr, pericardium; di, diaphragm; pa, pulmonary artery; pv, pulmonary vein. (**c**) Cross-sectional view of the boxed region in **b** (left), showing aortic valve. Asterisks indicate three leaflets comprise the valve. (**d**) Magnified view of the boxed region in **b** (right), acquired at different heights from the bottom of the specimen, showing multi-layered pericardium. i, myocardium; ii, serous pericardium; iii, fibrous pericardium. (**e**) Volumetric view of the heart tissue. Magenta, ATTO 647N NHS ester; Green, Alexa Fluor 488 NHS ester. (**f**–**g**) Confocal microscopy images of Alexa Fluor 488 NHS ester-stained heart muscle in (**f**) BABB-cleared mouse embryo and (**g**) expanded mouse embryo, respectively. (**h**) Magnified view of the boxed region in **e**, showing the details of the heart muscle. Inset displays magnified view of the single mitochondria. Magenta, ATTO 647N NHS ester; Green, Alexa Fluor 488 NHS ester. Scale bars: (**a**) 1 mm, (**b**) 500 µm, (**c**–**d**) 100 µm, (**e**) 10 µm, (**f**–**g**) 5 µm, (**h**) 1 µm, and inset in (**h**) 500 nm. All length scales are presented in pre-expansion dimensions.

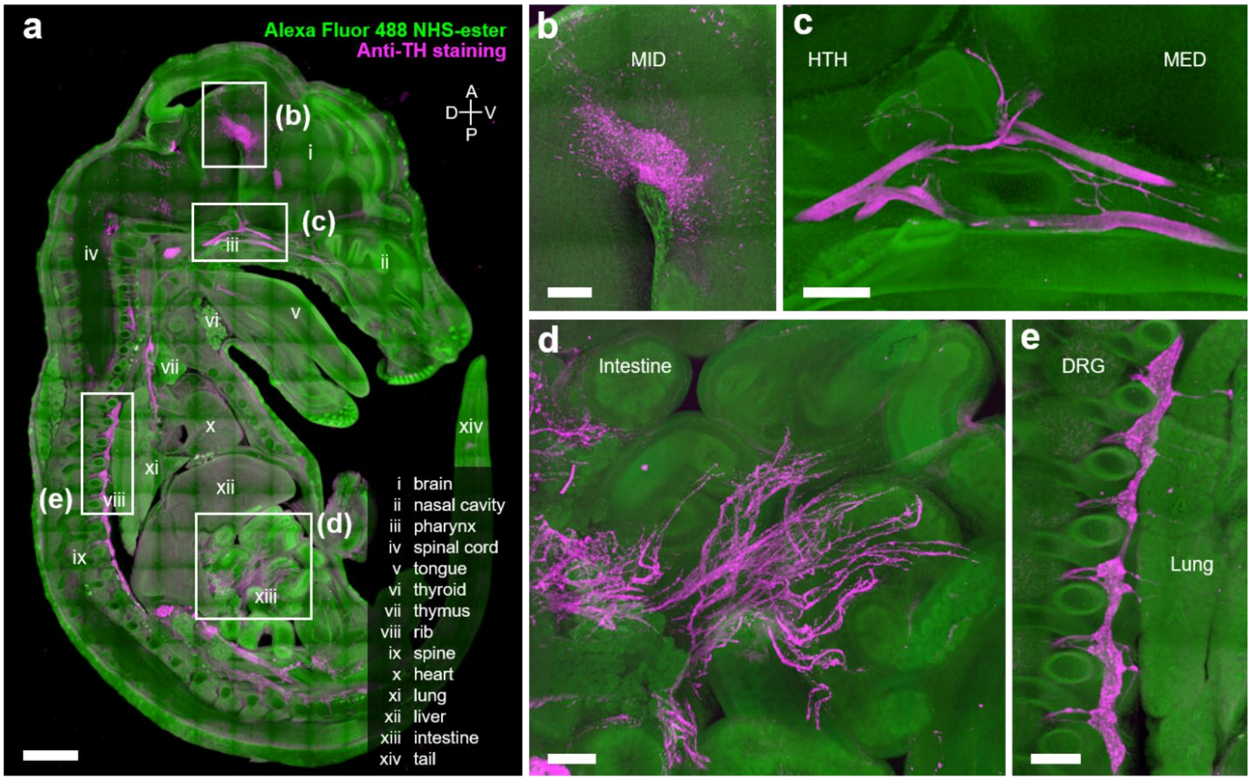


**Supplementary Fig. 21**: **Whole-body ExM imaging of mouse embryo labeled with antibodies and fluorophore NHS ester**. (**a**) Confocal microscopy image of anti-TH antibody and Alexa Fluor 488 NHS-ester-labeled E15.5 mouse embryo slice after expansion. Green, Alexa Fluor 488 NHS-ester; Magenta, anti-TH antibody. (**b**–**e**) Magnified view of the boxed regions in **a**, showing (**b**) brain, (**c**) pharynx, (**d**) intestine, and (**e**) rib, respectively. MID, midbrain; HTH, hypothalamus; MED, medulla; DRG, dorsal root ganglion. Scale bars: (**a**) 1 mm, and (**b**–**e**) 200 μm. All length scales are presented in pre-expansion dimensions.

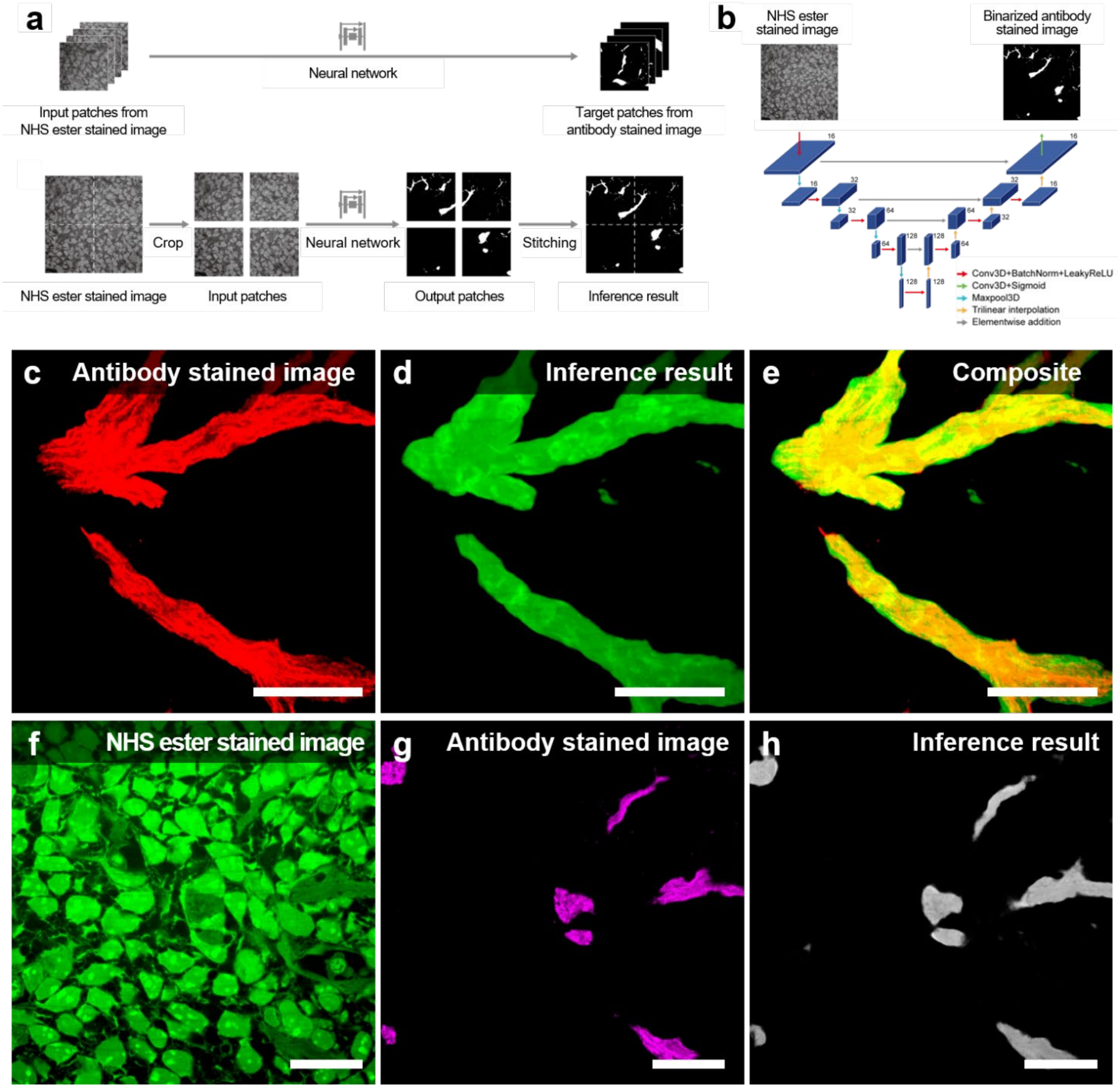
a
Neural network
Input patches from
NHS ester stained image
Target patches from
antibody stained image
Crop
Neural network
Stitching
NHS ester stained image
Input patches
Output patches
Inference result
b
NHS ester
stained image
Binarized antibody
stained image
Conv3D+BatchNorm+LeakyReLU
Conv3D+Sigmoid
Maxpool3D
Trilinear interpolation
Elementwise addition
c Antibody stained image
d Inference result
e Composite
f NHS ester stained image
g Antibody stained image
h Inference result

**Supplementary Fig. 22**: **Axon bundle identification from the non-specific protein label channels using machine learning**. (**a**) Training and testing pipelines. The neural network is trained to infer axonal morphology from fluorophore NHS-ester stained images. Input and target pairs are partitioned from the fluorophore NHS-ester stained channel and the TUJ1 antibody stained channel and used for network training (upper). An unseen fluorophore NHS-ester stained image is partitioned into 3D patches and fed to the network. The inference result over a large volume is obtained by stitching the output patches (lower). (**b**) Network architecture for inference of axonal morphology from fluorophore NHS-ester stained images. Three-dimensional images obtained with fluorophore NHS-ester staining and TUJ1 antibody staining are used as the input and the target, respectively. The network employs a 3D U-net architecture that consists of a 3D encoder, a 3D decoder, and four skip connections from the encoder to the decoder. The skip connections pass the feature maps from the encoder to the decoder. Each blue box corresponds to a multi-channel feature map. The number of channels is noted on the right side of each box. (**c**–**e**) Representative example of inferred axon morphology. **c**, antibody stained image (ground truth), **d**, inference result, **e**, composition of **c** and **d**. (**f**–**h**) Representative example of inferred axon morphology. **f**, fluorophore NHS-ester stained image, **g**, antibody stained image, **h**. inference result. Scale bars: 20 μm. All length scales are presented in pre-expansion dimensions.

### SUPPLEMENTARY VIDEOS

**Supplementary Video 1**. **Z-stack image of an expanded mouse brain slice showing the nanoscale details of mitochondria and ER structures in the brain**. Gray, Cy3 NHS ester.

**Supplementary Video 2**. **Z-stack image of an expanded mouse small intestine slice**. Gray, ATTO 647N NHS ester; blue, DAPI.

**Supplementary Video 3**. **Z-stack image of an expanded mouse kidney slice**. Gray, ATTO 647N NHS ester; blue, DAPI.

**Supplementary Video 4**. **Z-stack image of an expanded mouse lung slice and its three-dimensional view**. Gray, ATTO 647N NHS ester; blue, DAPI.

**Supplementary Video 5**. **Z-stack image of an expanded mouse heart slice**. Gray, ATTO 647N NHS ester; blue, DAPI.

**Supplementary Video 6**. **Montage image of an expanded whole zebrafish larva**. Gray, Alexa Fluor 488 NHS ester; blue, DAPI.

**Supplementary Video 7**. **Z-stack image of an expanded zebrafish olfactory sensory epithelium**. Gray, ATTO 647N NHS ester.

**Supplementary Video 8**. **Z-stack image of an expanded zebrafish trunk neuromast**. Gray, ATTO 647N NHS ester.

**Supplementary Video 9**. **Z-stack image of an expanded zebrafish inner ear.** Gray, Alexa Fluor 488 NHS ester.

**Supplementary Video 10**. **Z-stack image of gut bacteria found in an expanded zebrafish intestinal lumen**. Gray, Alexa Fluor 488 NHS ester.

**Supplementary Video 11**. **Z-stack image of an expanded zebrafish ciliated intestinal epithelium**. Gray, Alexa Fluor 488 NHS ester.

**Supplementary Video 12**. **Z-stack image of an expanded zebrafish mid-intestine**. Green, Alexa Fluor 488 NHS ester; magenta, ATTO 647N NHS ester.

**Supplementary Video 13**. **Z-stack image of expanded zebrafish musculoskeletal tissues**. Gray, Alexa Fluor 488 NHS ester.

**Supplementary Video 14**. **Z-stack image of an expanded zebrafish notochord**. Gray, ATTO 647N NHS ester.

**Supplementary Video 15**. **Z-stack image of an expanded zebrafish pectoral fin**. Gray, Alexa Fluor 488 NHS ester.

**Supplementary Video 16**. **Z-stack image of an expanded zebrafish retina**. Green, Alexa Fluor 488 NHS ester; magenta, ATTO 647N NHS ester.

**Supplementary Video 17**. **Montage image of an expanded mouse embryo slice**. Green, Alexa Fluor 488 NHS ester; gray, ATTO 647N NHS-ester; blue, DAPI.

**Supplementary Video 18**. **Z-stack image of an expanded half mouse embryo after expansion**. Green, Alexa Fluor 488 NHS ester.

**Supplementary Video 19**. **Three-dimensional view of expanded mouse embryos**. Green, Alexa Fluor 488 NHS ester; yellow, anti-TUJ1.

### SUPPLEMENTARY NOTES

**Supplementary Note 1**. **Effect of the hydrophobicity of fluorophores on staining patterns**. We hypothesized that the labeling of protein structures with fluorophore NHS esters would depend on the physicochemical properties of the fluorophores, especially their hydrophobicity. To verify this hypothesis, mouse brain slices were first permeabilized with Triton X-100 and treated with AcX. The slices were then embedded in a swellable hydrogel and digested with proteinase K. Subsequently, the slices were labeled with one of seven fluorophore NHS esters with different hydrophobicities and expanded (**Supplementary Fig**. **3**). As discussed in the manuscript, we found that protein structures labeled with fluorophore NHS esters depended on fluorophores' hydrophobicities. As shown in **Supplementary Fig**. **4a**, Cy3, the most hydrophobic fluorophore from our experimental set, showed globular and linear structures in the cytoplasm that corresponded to cellular organelles. The signal-to-noise ratio of the labeled mitochondria and ER was sufficiently high, allowing clear visualization of their 3D structures, as shown in **Supplementary Video 1**. Another hydrophobic fluorophore NHS ester, ATTO 647N, showed a similar staining pattern (**Supplementary Fig**. **4b**). In contrast, the hydrophilic Alexa Fluor 488 NHS ester strongly labeled the nucleoplasm and showed uniform staining in the cytoplasm (**Supplementary Fig**. **4c**).

**Supplementary Note 2**. **Staining patterns of hydrophobic and hydrophilic fluorophores**. The compatibility of our labeling technique with permeabilization enabled the co-staining of specimens with antibodies and fluorophore NHS esters. When a mouse brain slice was stained with Cy3 NHS ester and antibodies against the pre-synaptic marker Bassoon and post-synaptic marker Homer1, Cy3 NHS ester exhibited a band-like structure located between Homer1 and Bassoon but was more closely located to and overlapped with Bassoon (**Supplementary Fig**. **4d**). Such localization of Cy3 NHS ester close to the pre-synapse can be attributed to the staining of lipid-rich synaptic vesicles at the pre-synapses. We confirmed that the differences in the staining patterns depended on the hydrophobicity of fluorophores in mouse brain and heart slices. We used both hydrophobic and hydrophilic fluorophore NHS esters to label single specimens. In the mouse brain corpus callosum, where many myelinated nerve fibers are present, Alexa Fluor 488 NHS ester labeled the axons of the myelinated fibers, while ATTO 647N NHS ester labeled the surrounding myelin, as shown in **Supplementary Fig**. **4e**. In the mouse heart, Alexa Fluor 488 NHS ester visualized a greater number of anatomical structures than ATTO 647N NHS ester. Alexa Fluor 488 NHS ester visualized the anatomy of cardiac muscle cells, such as the Z-disc, A-band, M-line, and capillaries. In contrast, the ATTO 647N NHS ester showed strong labeling of the mitochondria (**Supplementary Fig**. **4f**). We investigated whether other physicochemical properties of fluorophores, such as their charges, affected the staining patterns. Both ATTO 565 and Cy3 had a zero net charge; however, ATTO 565 was more hydrophilic than Cy3. These two fluorophore NHS esters yielded completely different staining patterns, as shown in **Supplementary Fig**. **3f** and **h**, indicating that hydrophobicity was the key factor determining the labeling pattern produced by the fluorophore NHS esters. In addition, a recent study reported that the hydrophobicity of fluorophores, not their charges, determines the nonspecific binding of fluorophore-conjugated antibodies during single-molecule imaging.[1] The differences in the staining patterns, depending on the hydrophobicity of the fluorophores, indicated that the local environment around the primary amines of proteins would affect the labeling density of the fluorophore NHS esters. Hydrophobic fluorophore NHS esters preferentially labeled lipid-rich structures, such as the nuclear membrane, mitochondria, ER, and myelinated fibers, indicating that the use of a single fluorophore NHS ester would not be sufficient to visualize all anatomically relevant protein structures.

**Supplementary Note 3**. **Inter-digestion labeling with fluorophore NHS esters**. We aimed to determine when it would be optimal to apply fluorophore NHS esters to label more anatomically relevant structures in diverse mouse organs. We compared the images of specimens prepared using three different protocols. First, the specimens were labeled with fluorophore NHS esters and then embedded in a hydrogel. This process was called pre-gelation staining. Second, specimens were embedded in hydrogels, digested with proteinase K briefly (approximately 4 h), labeled with fluorophore NHS esters, and further digested with proteinase K. This process was called inter-digestion staining. Third, the specimens were embedded in hydrogels, repetitively digested with proteinase K, and labeled with fluorophore NHS esters. This process was called post-digestion staining. Inter-digestion labeling demonstrated the best staining performance in terms of the fluorescence intensity and diversity of the labeled structures among the three labeling methods. First, the fluorescence signal intensity of inter-digestion was higher than that of pre-gelation. This was attributed to the bleaching of the fluorophores during the in situ gelation step of the pre-gelation process. In addition, the superior labeling performance of the inter-digestion process could be explained by the exposure of primary amines buried in proteins after the first brief digestion process. During the pre-gelation staining process, the specimens were first treated with fluorophore NHS esters and then with AcX. As both fluorophore NHS esters and AcX reacted with primary amines, some proteins did not have enough fluorophore NHS esters for visualization or AcX for anchoring simultaneously. On the contrary, during the inter-digestion staining process, newly exposed primary amines could be available for labeling, and more diverse structures would be labeled and visualized. Such exposed primary amines would also boost the fluorescence signal by increasing the number of labeling sites. For example, in the mouse intestine, the inter-digestion process visualized individual microvillus of the brush borders that was not clearly observed *via* pre-gelation staining process (**Supplementary Fig**. **4g–k**). In the specimens prepared by the post-digestion staining process, a large number of aggregates were observed, especially in the liver. Such aggregates are the result of the nonspecific adsorption of protein fragments generated by extended proteinase K digestion. In the specimens prepared by the inter-digestion staining process, such aggregates were not observed, indicating that fluorophore-labeled fragments would not form aggregates due to the increased hydrophilicity upon labeling. In the mouse liver, both the pre-gelation and inter-digestion staining processes visualized mitochondria; however, only inter-digestion staining showed the Golgi apparatus (**Supplementary Fig**. **4l–q**).

**Supplementary Note 4**. **The staining patterns of ATTO 647N in the mouse brain**. As discussed in **Supplementary Note 1**, ATTO 647N strongly labeled cellular organelles, such as mitochondria, ER, and Golgi apparatus, in the mouse brain (**Supplementary Fig**. **5a–c**). We confirmed that mitochondria in the cytoplasm—and even those in neurites—were labeled with hydrophobic fluorophore NHS esters using a transgenic mouse line expressing mitochondria-targeted mScarlet (**Supplementary Fig**. **5d–f**). The hydrophobic fluorophore NHS esters strongly labeled the myelinated regions of neurons and blood vessels (**Supplementary Fig**. **5g–i**). Similar to Cy3, ATTO 647N was useful in visualizing synapses in the brain (**Supplementary Fig**. **6a–d**). We recently showed that F-actin is mainly localized in post-synaptic densities.[2] By combining hydrophobic fluorophore NHS ester labeling and ExM imaging of actin, we visualized the four-layered structures of the Bassoon, a hydrophobic fluorophore NHS ester, Homer1, and F-actin (**Supplementary Fig**. **6e–j**).

**Supplementary Note 5**. **ExM imaging of fluorophore NHS ester-labeled mouse digestive organs**. We employed the inter-digestion fluorophore NHS ester staining process to the organs of the mouse digestive system, such as the esophagus, stomach, and small intestine. In the expanded esophagus, the layered structures of the mucosa, which consist of the epithelium, lamina propria, muscularis mucosa, submucosa, and muscularis externa, were clearly identified (**Supplementary Fig**. **7a**). More detailed structures, such as the mitochondria in the epithelium and muscularis mucosa layers, were also visualized (**Supplementary Fig**. **7b**). In the expanded stomach, the stacked layers of mucosa, muscularis mucosa, submucosa, muscularis propria, subserosa, serosa, gastric pit (cyan bracket in **Supplementary Fig**. **7c**), gastric gland (yellow bracket in **Supplementary Fig**. **7c**), and lamina propria (magenta arrowhead in **Supplementary Fig**. **7c**) were clearly identified. In the expanded small intestine, histological structures, including the lumen, brush border, goblet cells, enterocytes, and lamina propria, were observed (**Supplementary Fig**. **7d**). More detailed structures, such as the goblet cells and mitochondria of enterocytes, are clearly shown in

**Supplementary Fig**. **7e** (see **Supplementary Video 2** for a z-stack image).

**Supplementary Note 6**. **ExM imaging of other representative mouse organs labeled with fluorophore NHS esters**. We employed an inter-digestion staining process performed using fluorophore NHS esters for the mouse kidney (urinary system), lung (respiratory system), heart (circulatory system), and testis (reproductive system). In the expanded kidney, major histological structures, such as distal convoluted tubule cells surrounding the glomerulus, macula densa, Bowman's space, vascular pole, brush border, and proximal convoluted tubule cells, were clearly visualized (**Supplementary Fig**. **8a**; see **Supplementary Video 3** for a z-stack image). More detailed structures, such as a single microvillus of the brush border, basal striations, outer membrane of the mitochondria, and individual foot processes, were observed (**Supplementary Fig**. **8b**–**f**). In the expanded testis, the primary spermatocyte, secondary spermatocyte, Leydig cells (magenta arrowhead), round spermatids (red arrowhead), and spermatozoa (**Supplementary Fig**. **8g**,**h**) were clearly shown. Interestingly, mouse sperm was also observed in the testis (**Supplementary Fig**. **8i**). In the expanded lung, the cilia and mitochondria of Clara cells in the bronchiole were observed (**Supplementary Fig**. **9a**–**c**). Moreover, alveolar cell type I (green arrowheads), the cytoplasmic extension of alveolar cell type I (white arrowhead), alveolar cell type II (yellow arrowheads), and secretory granules (cyan arrowheads) of alveolar cell type II in the alveolus of the lung were identifiable (**Supplementary Fig**. **9d**–**f**). Confocal microscopy imaging of the expanded lung enabled the three-dimensional visualization of such histological structures (**Supplementary Video 4**). In the expanded heart, major histological structures, such as the striation, capillaries, and mitochondria between cardiac muscle fibers (**Supplementary Fig**. **9g**–**h**), along with the sarcomere, Z-disc, M-line, I-band, and A-band of cardiac muscles (**Supplementary Fig**. **4f**; see **Supplementary Video 5** for a *z*-stack image), were clearly visualized.

### SUPPLEMENTARY TABLES

| Product name | Vendor | Product number |
|---|---|---|
| **Fixation and staining** | | |
| Agarose, low gelling temperature | Sigma-Aldrich | A9414 |
| DNase/RNase-free distilled water | Invitrogen | 10977-023 |
| Formamide | Sigma-Aldrich | F7503 |
| Glycine | Sigma-Aldrich | 50046 |
| Heparin | Sigma-Aldrich | H3149 |
| Normal donkey serum (NDS) | Jackson Immunoresearch | 017-000-002 |
| Normal goat serum (NGS) | Jackson Immunoresearch | 055-000-121 |
| Paraformaldehyde (PFA) | Electron Microscopy Science | 15710 |
| Phosphate-buffered saline (PBS) | Invitrogen | AM9625 |
| Saline-sodium citrate buffer (SSC) | Invitrogen | AM9625 |
| Salmon sperm DNA | Thermofisher | AM9680 |
| Sodium azide | Sigma-Aldrich | S2002 |
| Triton X-100 | Sigma-Aldrich | T9284 |
| **BABB clearing** | | |
| Benzyl alcohol | Sigma-Aldrich | E7023 |
| Benzyl benzoate | Sigma-Aldrich | 216763 |
| Ethyl alcohol | Sigma-Aldrich | 108006 |
| Hydroxy peroxide solution (30 %) | Sigma-Aldrich | B6630 |

| Product name | Vendor | Product number |
|---|---|---|
| **Expansion microscopy and fluophore NHS-ester staining** | | |
| 2,2'-azobis[2-(2-imidazolin-2-yl)propane] dihydrochloride (VA-044) | Wako Chemicals | 223-02112 |
| 4-hydroxy-2,2,6,6-tetramethylpiperidine-1-oxyl (H-TEMPO) | Sigma-Aldrich | 176141 |
| Acrylamide (AAm) | Sigma-Aldrich | A9099 |
| Ammonium persulfate (APS) | Sigma-Aldrich | A3678 |
| Calcium chloride | Sigma-Aldrich | C1016 |
| Dimethyl sulfoxide (DMSO) | Sigma-Aldrich | 276855 |
| Ethylenediaminetetraacetic acid (EDTA) | Sigma-Aldrich | EDS |
| *N, N′*-methylenebisacrylamide (BIS) | Sigma-Aldrich | M7279 |
| *N,N′*-(1,2-dihydroxyethylene)bisacrylamide (DHEBA) | Tokyo Chemical Industry | D2864 |
| *N,N,N′,N′*-tetramethylethylenediamide (TEMED) | Sigma-Aldrich | T7024 |
| Poly-L-lysine | Sigma-Aldrich | 25988-63-0 |
| Collagenase, Type I | Thermo Fisher | 17100017 |
| Collagenase, Type II | Thermo Fisher | 17101015 |
| Collagenase, Type IV | Thermo Fisher | 17104019 |
| Hanks' Balanced Salt Solution (HBSS) | Thermo Fisher | 14065056 |
| Proteinase K | New England Biolabs | P8107S |
| SE (6-((acryloyl)amino)hexanoic acid) (Acryloyl-X, AcX) | Thermo Fisher | A20770 |
| Sodium acrylate (AA) | Ambeed | A107105 |
| Sodium chloride | Sigma-Aldrich | 71376 |
| *Label IT*® Nucleic Acid Modifying Reagent | Mirus Bio LLC | MIR3900 |
| 3-(*N*-Morpholino)propanesulfonic acid (MOPS) | Sigma-Aldrich | M3183 |
| Tris hydrochloride solution | Sigma-Aldrich | T3038 |
| **Non-chemical supplies** | | |
| No.1 coverglasses | Brain Research Lab | 4860-1 |

**Supplementary Table 1**. **List of reagents used in this study**.

| Product name | Vendor | Product number | Stock conc. | Dilution ratio | Final conc. |
|---|---|---|---|---|---|
| **Primary antibody** | | | | | |
| Goat anti-FITC with Alexa Fluor 488 | Thermo Fisher | A-11096 | 1 mg/mL | 1:100 | 10 µg/mL |
| Mouse anti-Bassoon | Enzo | ADI-VAM-PS003-D | 1 mg/mL | 1:500 | 2 µg/mL |
| Mouse anti-TUJ1(TUBB3) | BioLegend | 801202 | 1 mg/mL | 1:100 | 10 µg/mL |
| Mouse anti-HuC/D(16A11) | Thermo Fisher | A21271 | 1 mg/mL | 1:100 | 10 µg/mL |
| Mouse anti-TH | Immunostar | 22941 | - | 1:1000 | - |
| Rabbit anti-Homer1 | Synaptic Systems | 160 003 | 1 mg/mL | 1:500 | 2 µg/mL |
| Rabbit anti-SLC2A1 | ATLAS | HPA031345 | 0.4 mg/ml | 1:1000 | 0.4 µg/mL |
| Rabbit anti-RFP | Abcam | ab62341 | 0.5 mg/ml | 1:1000 | 5 µg/mL |
| Rat anti-MBP | Abcam | ab7349 | - | 1:100 | - |
| **Secondary antibody** | | | | | |
| Goat anti-mouse CF 405S | Biotium | 20080 | 1 mg/mL | 1:100 | 10 µg/mL |
| Goat anti-mouse Alexa Fluor 488 | Thermo Fisher | A32723 | 1 mg/mL | 1:200 | 5 µg/mL |
| Goat anti-mouse Alexa Fluor 546 | Thermo Fisher | A11030 | 1 mg/mL | 1:200 | 5 µg/mL |
| Goat anti-rat Alexa Fluor 488 | Thermo Fisher | A11006 | 2 mg/mL | 1:200 | 10 µg/mL |
| Goat anti-rabbit Alexa Fluor 488 | JacksonImmuno Research | 111-545-144 | 1 mg/mL | 1:200 | 5 µg/mL |
| Goat anti-rabbit Alexa Fluor 546 | Thermo Fisher | A11035 | 2 mg/mL | 1:200 | 10 µg/mL |
| Goat anti-rabbit CF 633 | Biotium | 20122 | 1 mg/mL | 1:100 | 10 µg/mL |
| Donkey anti-mouse Alexa Fluor 546 | Thermo Fisher | A10036 | 2 mg/mL | 1:1000 | 2 µg/mL |

| Product name | Vendor | Product number | Stock conc. | Dilution ratio | Final conc. |
|---|---|---|---|---|---|
| **Chemical dye** | | | | | |
| DAPI | Sigma-Aldrich | D9542 | 1 mg/mL | 1:1000 | 1 µg/mL |
| Phalloidin-FITC | Thermo Fisher | F432 | 66 µM | 1:40 | 1.65 µM |
| Dyomic 480XL NHS ester | Dyomics | 480XL-01 | 10 mg/mL | 1:1000 | 10 µg/mL |
| Alexa Fluor 488 NHS ester | Thermo Fisher | A20000 | 10 mg/mL | 1:1000 | 2–10 µg/mL |
| ATTO 488 NHS ester | ATTO-Tec | AD 488-31 | 10 mg/mL | 1:1000 | 10 µg/mL |
| CF 568 NHS ester | Biotium | 92131 | 10 mM | 1:500 | 2 µM |
| Alexa Fluor 546 NHS ester | Thermo Fisher | A20002 | 10 mg/mL | 1:1000 | 10 µg/mL |
| ATTO 565 NHS ester | ATTO-Tec | AD 565-31 | 10 mg/mL | 1:1000 | 10 µg/mL |
| CF 633 NHS ester | Biotium | 92133 | 10 mM | 1:500 | 2 µM |
| ATTO 633 NHS ester | ATTO-Tec | AD 633-31 | 10 mg/mL | 1:1000 | 10 µg/mL |
| ATTO 647N NHS ester | ATTO-Tec | AD 647N-31 | 10 mg/mL | 1:1000 | 2–10 µg/mL |
| Cy3 NHS ester | Kerafast | FLP124 | 10 mg/mL | 1:1000 | 2–10 µg/mL |

**Supplementary Table 2. List of labeling reagents and their final concentration for use**.
